# CBP/EP300 acetylates and stabilizes the stress-responsive Heat Shock Factor 2, a process compromised in Rubinstein-Taybi syndrome

**DOI:** 10.1101/481457

**Authors:** Aurélie de Thonel, Johanna K. Ahlskog, Ryma Abane, Geoffrey Pires, Véronique Dubreuil, Jérémy Berthelet, Anna L. Aalto, Sarah Naceri, Marine Cordonnier, Carène Benasolo, Matthieu Sanial, Agathe Duchateau, Anniina Vihervaara, Mikael C. Puustinen, Federico Miozzo, Mathilde Henry, Déborah Bouvier, Jean-Paul Concordet, Patricia Fergelot, Élise Lebigot, Alain Verloes, Pierre Gressens, Didier Lacombe, Jessica Gobbo, Carmen Garrido, Sandy D. Westerheide, Michel Petitjean, Olivier Taboureau, Fernando Rodrigues-Lima, Madeline Lancaster, Sandrine Passemard, Délara Sabéran-Djoneidi, Lea Sistonen, Valérie Mezger

## Abstract

Cells respond to protein-damaging insults by activating heat shock factors (HSFs), key transcription factors of proteostasis. Abnormal levels of HSFs occur in cancer and neurodegenerative disorders, highlighting the strict control of their expression. HSF2 is a short-lived protein, which is abundant in the prenatal brain cortex and required for brain development. Here, we report that HSF2 is acetylated and co-localized with the lysine-acetyl transferases CBP and EP300 in human brain organoids. CBP/EP300 mediates the acetylation of HSF2 on specific lysine residues, through critical interaction between the CBP-KIX domain and the HSF2 oligomerisation domain, and promotes HSF2 stabilization. The functional importance of acetylated HSF2 is evidenced in Rubinstein-Taybi syndrome (RSTS), characterized by mutated CBP or EP300. We show that cells derived from RSTS patients exhibit decreased HSF2 levels and impaired heat shock response. The dysregulated HSF pathway in RSTS opens new avenues for understanding the molecular basis of this multifaceted pathology.

## INTRODUCTION

Since their discovery three decades ago, our way to envision the regulation and roles of the Heat Shock transcription Factor family (HSFs) has been revolutionized. Originally identified and characterized due to their stress-responsiveness and ability to recognize a consensus DNA-binding site, the heat shock element (HSE), HSFs were more recently shown to perform an unanticipated large spectrum of roles under physiological and pathological conditions (Wu, 1995; Abane and Mezger 2010; Akerfelt et al., 2010; Pastor-Gomez et al., 2018). HSFs are activated by a diversity of stressors that provoke protein damage and govern the highly conserved Heat Shock Response (HSR). The HSR contributes to the restoration of proteostasis, through the regulation of genes encoding molecular chaperones, including the Heat Shock Proteins (HSPs; Hartl et al., 2011). HSFs also control immune/inflammatory pathways, metabolism, and, through dysregulation of their protein levels or activity, shape disease susceptibility to cancer, metabolic and neurodegenerative disorders. These pathophysiological roles are performed through altered expression of a broad repertoire of target genes, beyond the *HSPs* (Xiao et al., 1999; Inouye et al., 2007; Dai et al., 2007; Mendillo et al., 2012; Santagata et al., 2013; Anckar and Sistonen, 2011; Jin et al., 2011; Neef et al., 2011; Nakai, 2016; Pastor-Gomez et al., 2017 and 2018). The multifaceted roles of HSFs are achieved by their fascinating plasticity in terms of multi-modular structure and assembly of homo- or heterodimers or trimers, stress- and context-dependent posttranslational modifications, as well as a diversity of partner networks. As a consequence, HSFs act as fine sculptors of transcriptomic and epigenetic landscapes, through dynamic interactions with other transcriptional activators or repressors and chromatin remodelling complexes (Akerfelt et al., 2010; Miozzo et al., 2015; Pastor-Gomez et al., 2018; Raychaudhuri et al. 2014).

The versatile functions of HSFs have been mostly studied with HSF1 and HSF2, two members of the mammalian HSF family, which in human comprises four additional members, *i.e*. HSF4, HSF5, HSFX and HSFY (Pastor-Gomez et al., 2018). The role of HSF1 in acute and severe proteotoxic stress, including exposures to elevated temperatures (42-45°C), has been extensively documented and has become a paradigm for the *modus operandi* of the HSF family. In contrast, HSF2 appears to be responsive to stresses of relevance for chronic or pathological situations, such as fever-like temperatures at 39-41°C (Shinkawa et al., 2011), alcohol (ethanol) exposure (El Fatimy et al., 2014; Miozzo et al., 2018), and prolonged proteasome inhibition (Lecomte et al., 2010; Rossi et al., 2014). Both factors have been associated with different forms of cancer, in a dual manner; HSF1 acts as a potent facilitator of cancer initiation and progression (Dai et al., 2007; Mendillo et al., 2012; Santagata et al., 2013), whereas HSF2 can counteract tumor progression and invasiveness (Björk et al. 2016). The control of HSF1 protein levels is key for its pathophysiological functions, as elevated expression of HSF1 correlates with poor cancer prognosis (Santagata et al., 2011) and decreased expression has been found in different neurodegenerative disorders (Kim et al., 2015; Jiang et al., 2013; Pastor-Gomez et al., 2017; reviewed in Pastor-Gomez et al., 2018). HSF1 and HSF2 are also important players in the physiological brain development and adult brain integrity. Furthermore, we and others have shown that deregulated their activities underlie both neurodevelopmental defects (Kallio et al., 2002; Wang et al., 2003; Chang et al., 2006; El Fatimy et al., 2014; Hashimoto-Torii et al., 2014; Ishii et al., 2017; reviewed in Abane et Mezger 2010, Åkerfelt et al., 2010, Pastor-Gomez et al., 2018) and neurodegenerative processes (Shinkawa et al., 2011; Pastor-Gomez et al., 2017).

The amount of HSF2 protein varies in diverse cellular or embryonic contexts and conditions: both transcriptional and post-transcriptional mechanisms have been shown to regulate *HSF2* mRNA levels (Rallu et al., 1997; Björk et al., 2010). In the developing brain, the protein levels of HSF2 seem to correlate with those of *Hsf2* transcripts, emphasizing the importance of the regulatory mechanisms of *Hsf2* gene expression (Rallu et al., 1997; Kallio et al., 2002; Wang et al., 2003). Moreover, HSF2 is a short-lived protein and its stabilization constitutes an important step controlling the DNA-binding activity of HSF2 (Sarge et al., 1993; Mathew et al., 1998; Kawazoe et al., 1998) and mediating its role in physiological processes and stress responses. HSF2 protein levels also fluctuate during the cell cycle, which further shows that stabilization of HSF2 provides with a critical control step in fine-tuning the HSR (Elsing et al., 2014). Indeed, HSF2 modulates the stress-inducible expression of *HSP* genes, which is primarily driven by HSF1 (Östling et al., 2007). This transient modulatory function of HSF2 is due to the rapid poly-ubiquitination and proteasomal degradation in response to acute heat stress (Ahlskog et al., 2010). While diverse posttranslational modifications (PTMs), such as phosphorylation and acetylation, are well known to control HSF1 stability (Kourtis et al., 2015; Pastor-Gomez et al., 2017; Raychaudhuri et al., 2014; reviewed in Pastor-Gomez et al., 2018), the mechanisms regulating the stability of HSF2 are poorly understood, and given its role in chronic stress, cancer, and physiopathological developmental processes, they are crucial to be elucidated.

The histone/lysine-acetyl transferases (HATs/KATs) CBP (CREBBP, CREB-binding protein; KAT3A) and EP300 (E1A-binding protein p300; KAT3B) control the stability of many transcription factors through their acetylation, including HSF1 (Thakur *et al*., 2013; Raychaudhuri et al., 2014). Heterozygous mutations in one of these KATs lead to Rubinstein-Taybi syndrome (RSTS; Lopez-Atalaya et al., 2014; Spena et al., 2015a). RSTS is a rare disease characterized by multiple congenital anomalies, neurodevelopmental defects, childhood cancer susceptibility, and vulnerability to infections (*CREBBP/CBP* mutation, RSTS1, OMIM #180849*; EP300* mutation, RSTS2; OMIM #613684). Here, we show that HSF2 is acetylated during normal brain development in human organoids, where it is expressed in the same territories as CBP and EP300. We demonstrate that CBP/EP300 mediates the acetylation of HSF2 on specific lysine residues, through critical interaction between the CBP-KIX domain and the HSF2 oligomerisation domain, thereby promoting the stabilization of the HSF2 protein. We then interrogate the functional importance of this regulation in the pathological context of RSTS. We observe a proteasomal-dependent reduction in HSF2 protein levels in cells derived from RSTS patients, which results in impairement of their ability to mount a proper heat shock response. The disruption of the HSR pathway in RSTS highlights the importance of the CBP/EP300-dependent regulation of HSF2 by acetylation and provides a new conceptual frame for understanding the molecular basis of this complex pathology.

## RESULTS

### HSF2 is acetylated and interacts with CBP/EP300 in the developing brain

HSF2 is abundantly expressed in the vertebrate developing brain, where it exhibits spontaneous DNA-binding activity, in unstressed, physiological conditions (Rallu et al., 1997; Kawazoe et al. 1999; Kallio et al., 2002; Wang et al., 2003; Chang et al., 2006). As a first step to determine whether the acetylation of HSF2 was involved in controlling its stability, similarly to the EP300-mediated acetylation of HSF1 (Raychaudhuri et al., 2014), we compared HSF2 and CBP/EP300 expression profiles, and investigated whether HSF2 acetylation could be detected in the developing mammalian cortex, since CBP and EP300 have key roles in neurodevelopment (reviewed in Chan and La Thangue 2001; Lopez-Atalaya et al., 2014).

To the best of our knowledge, the expression of the HSF2 protein in the human developing cortex has not been reported. By generating brain organoids from human embryonic stem cells (Lancaster et al., 2014), we confirmed that, HSF2 mRNAs are present in human brain organoids, as previously reported (Figure S1A; Camp et al., 2015) and observed that *CBP* and *EP300* mRNA are also present. We found that the HSF2 protein was expressed at different stages, from day 20 (embryoid bodies) to day 60 of differentiation, with a profile similar to that of CBP and EP300 (D20 – D60; Figure 1A). By immunofluorescence on D60 organoids, we found that HSF2 was expressed in neural progenitor cells (NPCs; located in areas of dense DAPI-staining), as verified by SOX2 staining (Figure S1B) and in neurons, expressing beta-III tubulin, in regions displaying a cortical-like morphology (Figure 1B and S1B). In addition, we observed co-labeling of EP300 and HSF2 in NPCs (arrowheads) and neurons (arrows) (Figure 1C). Thus, HSF2 and EP300 (and also CBP; see Figure S1C) exhibited similar expression territories (NPCs and neurons; Figure S1B,C). In addition, HSF2, CBP and EP300 were expressed, in a concomitant manner, in the mouse cortex from E11 to E17 (Figure S1D). The similarity of their expression patterns suggested an interaction between HSF2 and CBP/EP300 in the developing cortex. Accordingly, HSF2 was co-immunoprecipitated with CBP and EP300 (Figure 1D and Figure S1E, left panels). We also detected acetylated HSF2 in the developing mouse cortex (Figure 1E and Figure S1E, middle panels) and in D40 human brain organoids (Figure 1F). Similarly, HSF2 was found acetylated in SHSY-5Y neuroblastoma cells, a human cancer cell line of neural origin (Figure S1F). Altogether, these results show that HSF2 is acetylated and interacts with CBP and EP300 in the developing brain.

**Figure 1.**
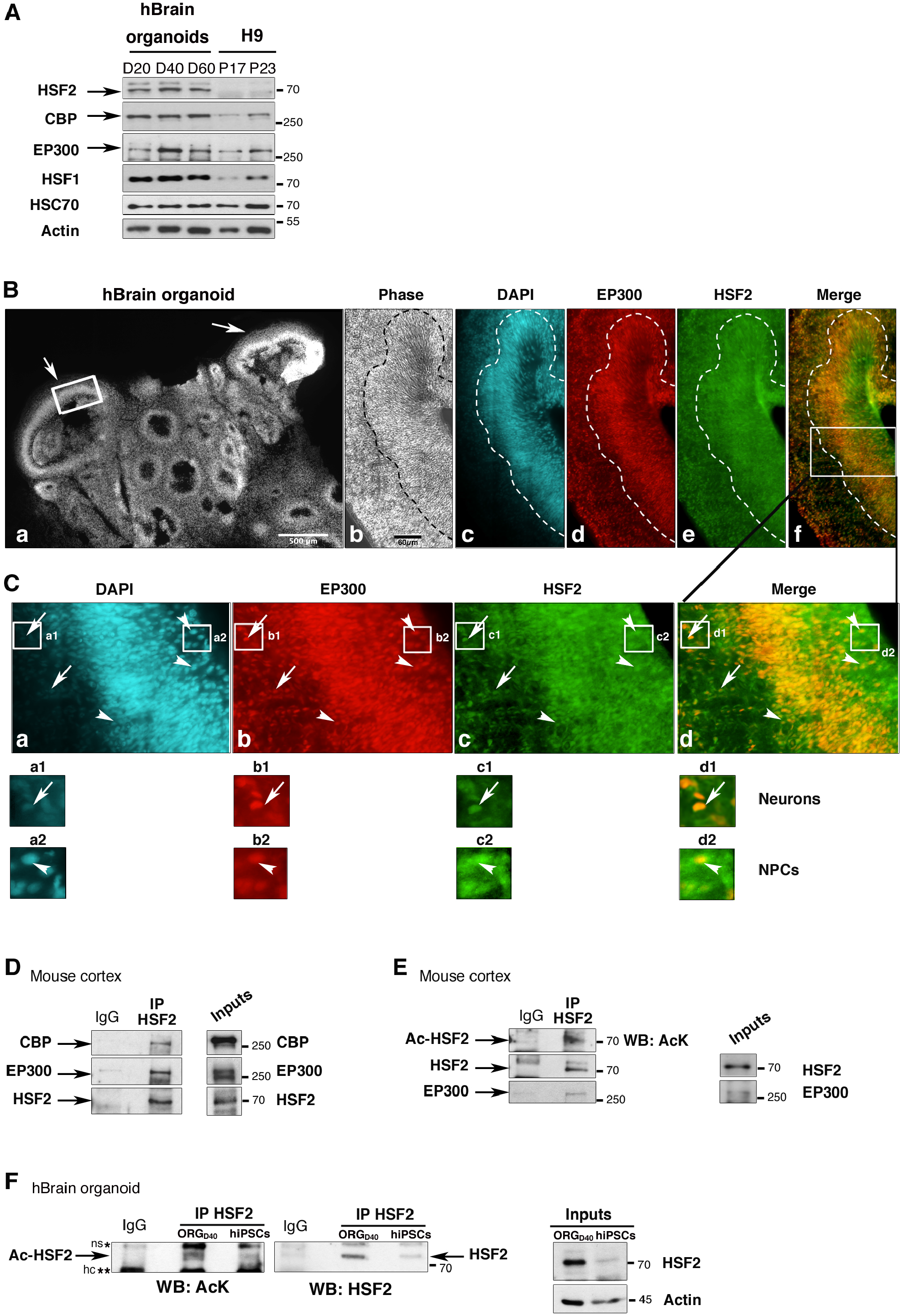
HSF2 expression profiles, acetylation status, and interaction with EP300/CBP in brain development. **(A to C) *HSF2 and CBP/EP300 in human brain organoids exhibit similar expression profiles and territories.*** **(A)** Representative immunoblot of extracts from human brain organoids at day 20, 40, and 60 of *ex vivo* development (D20, D40, D60) and human embryonic stem (ES) cells (H9) at passage 17 and 23. The position of molecular weight markers is indicated (kDa). HSC70 is heat shock cognate protein, which not induced by stress and is commonly used as a loading control. Actin, which increases during organoid differentiation, was also used for comparison. (See also Figure S1A). **(B)** (a) Microscopy epifluorescence images of a D60 human organoid. (a) DAPI-staining of the complete section (image reconstruction), showing structures reminiscent of the developing cerebral cortex (arrows). The thick white rectangle indicates the magnified areas shown in (b-f). (b) Phase contrast; (c) DAPI-staining; (d-f) Immunostaining for EP300 (d, f, red) and HSF2 (e, f, green), and merge (f). The thin rectangle in (f) indicates the area magnified in (**C**). Scale bars: in (a), 500 µm; in (b), 60 µm. **(C)** Magnification of the cortical-like area indicated by the thin rectangle in (**B**, f). (a) DAPI-staining; (b-d) Immunostaining for EP300 (b) and HSF2 detection (c), and merge (d). (a1-d1) and (a2-d2) correspond to magnified regions in the zone of neurons (low DAPI density, Tuj1 positive region, see Figure S1B;e,j) and NPCs (high DAPI density, Sox2-positive region, see Figure S1B;h,i), respectively, indicated by white squares in (a, b, c, d). HSF2 and EP300 are co-expressed in some neurons (long arrows;) and NPCs (arrowheads; dense DAPI-stained regions). (See also Figure S1B; d,e). **(D and E) *HSF2 interacts with EP300 and CBP, and is present in an acetylated form in the developing cortex.*** **(D)** Endogenous HSF2 and EP300 proteins are co-immunoprecipitated in mouse E16 cortical extracts. (See also Figure S1E (E10 stage)). (Left panels) After immunoprecipitation of HSF2, the co-precipitated CBP or EP300 proteins were detected by Western blot analysis WB. (Right panels) total amounts proteins in the input samples. *: IgG heavy chain. Representative immunoblots (n=3 experiments). **(E)** Acetylation of the immunoprecipitated HSF2 protein from E15 mouse cortical extracts was assessed using anti-pan-acetyl-lysine antibody (AcK; see also Figure S1E (E10 stage)). Co-immunoprecipitation of EP300 is shown as a positive control. Representative immunoblots (n= 2 experiments). **(F)** HSF2 is acetylated in human brain organoids. (Left panel) Immunoprecipitation of the HSF2 protein in D40 organoids and immunoblotting with an anti-AcK antibody. (Middle panel) reincubation of the blot with anti-HSF2 antibody. (Right panel) total amounts proteins in the input samples. hIPSCs were used as comparison as they contain with low levels of HSF2. ns*, non specific; hc**, IgG heavy chain. Actin was used as a loading control.

### Analysis of CBP/EP300-mediated HSF2 acetylation

In order to explore the mechanism of HSF2 acetylation, we first examined whether HSF2 was a substrate for acetylation by CBP/EP300, in human HEK 293 cells co-expressing CBP-HA or EP300-HA and GFP- or Myc-tagged HSF2. We found that the immunoprecipitated exogenous HSF2 protein was acetylated by EP300 or CBP (Figure 2A), but that no acetylation was observed in cells transfected by dominant-negative CBP, unable to catalyze acetylation (Figure 2B).

**Figure 2.**
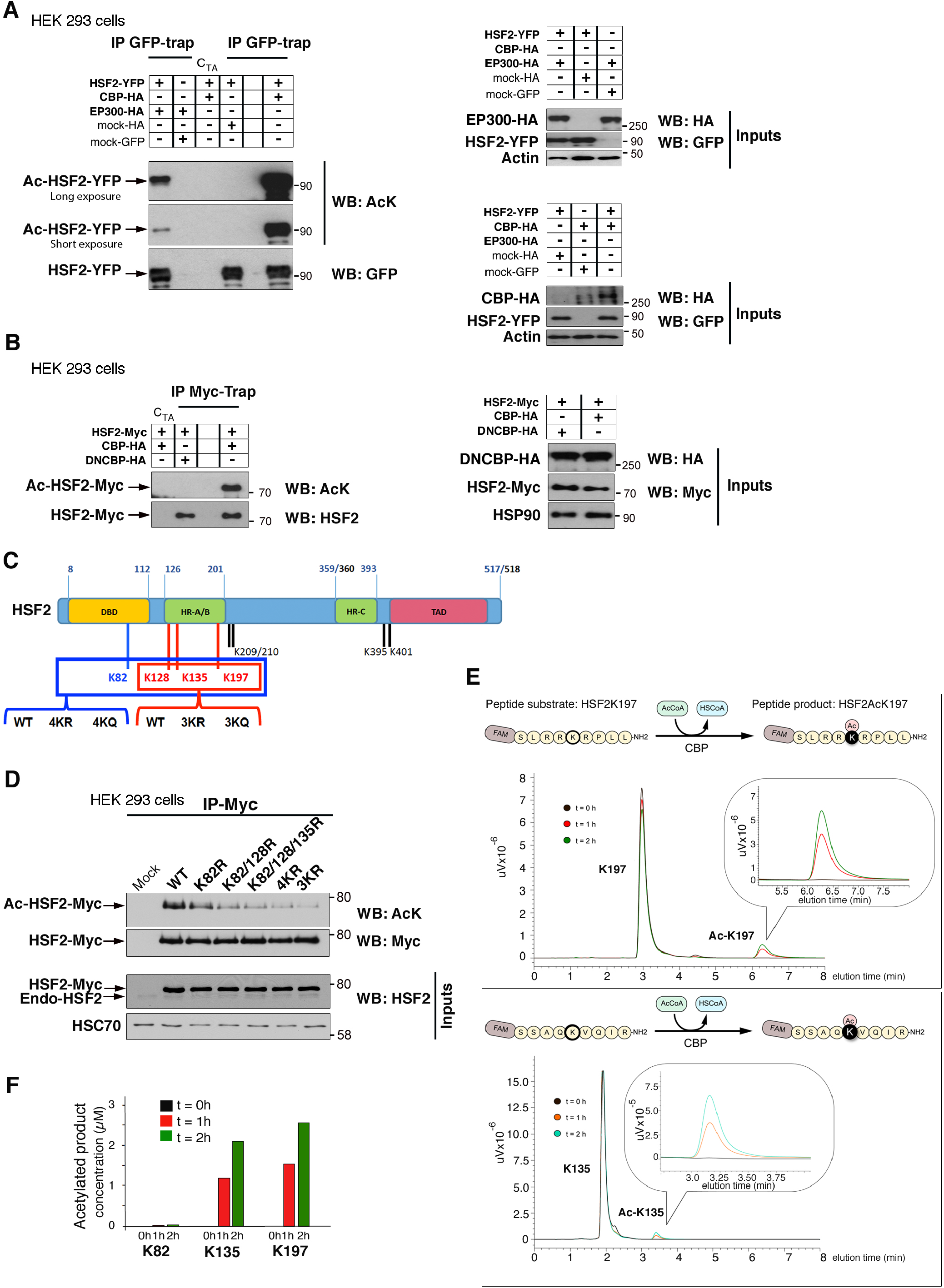
HSF2 is acetylated by CBP and EP300 in normal conditions. **(A) *The ectopically expressed YFP-HSF2 protein is acetylated by exogenous HA-CBP or EP300.*** Representative immunoblots (n=5 independent experiments). HEK 293 cells were transfected with different combinations of YFP-HSF2, HA-CBP, HA-EP300 constructs, and mock-HA or -GFP constructs. YFP-HSF2 was immunoprecipitated using anti-GFP-trap antibody (IP GFP-Trap) and its acetylation status was determined by WB analyses, using an anti-pan-acetyl-lysine antibody (AcK; left panels). Immunoprecipitated HSF2 was detected using an anti-GFP antibody (WB: GFP). The total amounts of the proteins in the input samples were detected with anti-GFP or anti-HA antibodies (inputs; right panels). Actin, loading control. C_TA_:Trap®-A beads were used as a negative control (see Experimental Procedures). **(B) The HSF2-Myc protein is not acetylated by a dominant-negative form of CBP (DNCBP-HA).** HEK 293 cells were transfected as in (**A**), except that HSF2-Myc was used instead of HSF2-YFP, and immunoprecipitation was performed using anti-Myc-trap antibody (IP Myc-Trap). Representative immunoblots (n=2 experiments). C_TA_: Trap®-A beads were used as a negative control (see Experimental Procedures). HSP90: loading control. **(C) Schematic representation of the eight main acetylated lysine residues of the HSF2 protein.** Purified mouse Flag-HSF2, co-expressed with HA-EP300, immunoprecipitated and subjected to MS analysis for detection of acetylated lysine residues. The three lysine residues K128, K135, K197, located in the oligomerization domain (HR-A/B), are enlightened in red and K82, located in the DBD, in blue; the other four lysine residues (K209/K210, K395/K401) are indicated in regular black. The DNA-binding domain (DBD, orange); the oligomerization domain (HR-A/B; green) and the domain controlling oligomerization (the leucine-zipper-containing HR-C; green); as well as the N-terminal domain (activation domain TAD; red) are illustrated. The boundaries of each domain are indicated in bold and blue. Bold and blue numbers correspond to the number of the amino acids located at boundaries of the domains of the mouse HSF2 protein, numbered from the +1 (ATG); the equivalent in the human HSF2 protein, if different, are indicated in bold and black. These four (K82, K128, K135, K197) or three lysine residues (K128, K135, K197) were mutated into glutamines (4KQ or 3KQ, respectively) or arginines (4KR or 3KR, respectively; see also Figure S2A). **(D) The mutations of three or four lysine (K) residues to arginine (3KR or 4KR) or glutamine residues (3KQ or 4KQ) decrease global HSF2 acetylation levels.** HEK 293 cells were co-transfected with EP300-HA and wild-type (WT) or mutated human HSF2-Myc on the indicated lysine residues. After immunoprecipitation of HSF2, using anti-Myc antibody, its acetylation was analysed by WB using an anti-AcK antibody. HSC70, loading control. n=3 independent experiments. (See also Figure S2B,C). **(E) *In vitro* acetylation of HSF2 peptides containing the K135 or K197 residues by recombinant CBP Full-HAT**. Time course of reverse phase-ultra-fast liquid chromatography (RP-UFLC) analysis of the acetylation of HSF2K197 (upper panel) and HSF2K135 peptides (lower panel) by CBP Full-HAT. Aliquots of the reaction were collected at 0 (black), 1 (red) or 2 (green) hours and elution of peptides was monitored by fluorescence emission at 530 nm (excitation: 485 nm, uV: arbitrary unit of fluorescence; see Figure S2D for HSF2K82 peptide). **(F) Quantification of the *in vitro* acetylated HSF2 peptides containing K82, K135, and K197 residues.** The AUC (area under the curve) of the acetylated K82, K135 and K197 peptides was quantified and converted in product concentration using a calibrated curve of various known concentrations of peptides. Note that it was not possible to investigate the acetylation of the HSF2 K128 peptide by CBP, because this peptide was repeatedly insoluble at the synthesis steps (Manufacturer’s information; see also Figure S2D-F).

To identify the acetylated lysine residues in HSF2, we co-expressed Flag-HSF2 with EP300-HA in HEK 293 cells. HSF2 was immunoprecipitated and the acetylation of lysines was analyzed by mass spectrometry (MS). Among the 36 lysine residues of HSF2, we identified eight acetylated lysines: K82 (located in the DNA-binding domain), K128, K135, K197 (all located within the hydrophobic heptad repeat HR-A/B), K209, K210, K395, and K401 (Figure 2C, Figure S2A, and Table S1 and S2). Single point mutations (K82, K128, K135, and K197), or mutation of the doublet K209/K210 to arginine (R, which prevent acetylation), did not abolish global HSF2 acetylation (Figure S2B). This suggests that, in line with our MS data, the acetylation of HSF2 occurs on more than one lysine residue. Indeed, the mutation to either arginine (R) or glutamine (Q) of the three or four lysines K82, K128, K135 and K197, dramatically reduced HSF2 acetylation (Figure 2D and Figure S2C). To dissect the requirement of CBP in the acetylation of HSF2, we used an *in vitro* acetylation assay coupled with HPLC (high-performance liquid chromatography). We found that a synthetic HSF2 peptide containing either K135 or K197 residues was readily acetylated by the purified recombinant full-catalytic domain (Full-HAT) of CBP (Figure 2E; Figure 3A) in an acetyl-CoA-dependent manner, whereas a peptide containing K82 was not (Figure S2D-F). Taken together, our data suggest that HSF2 is acetylated by CBP/EP300 at three main lysine residues, residing in the oligomerization HR-A/B domain: K128, K135 and K197.

**Figure 3.**
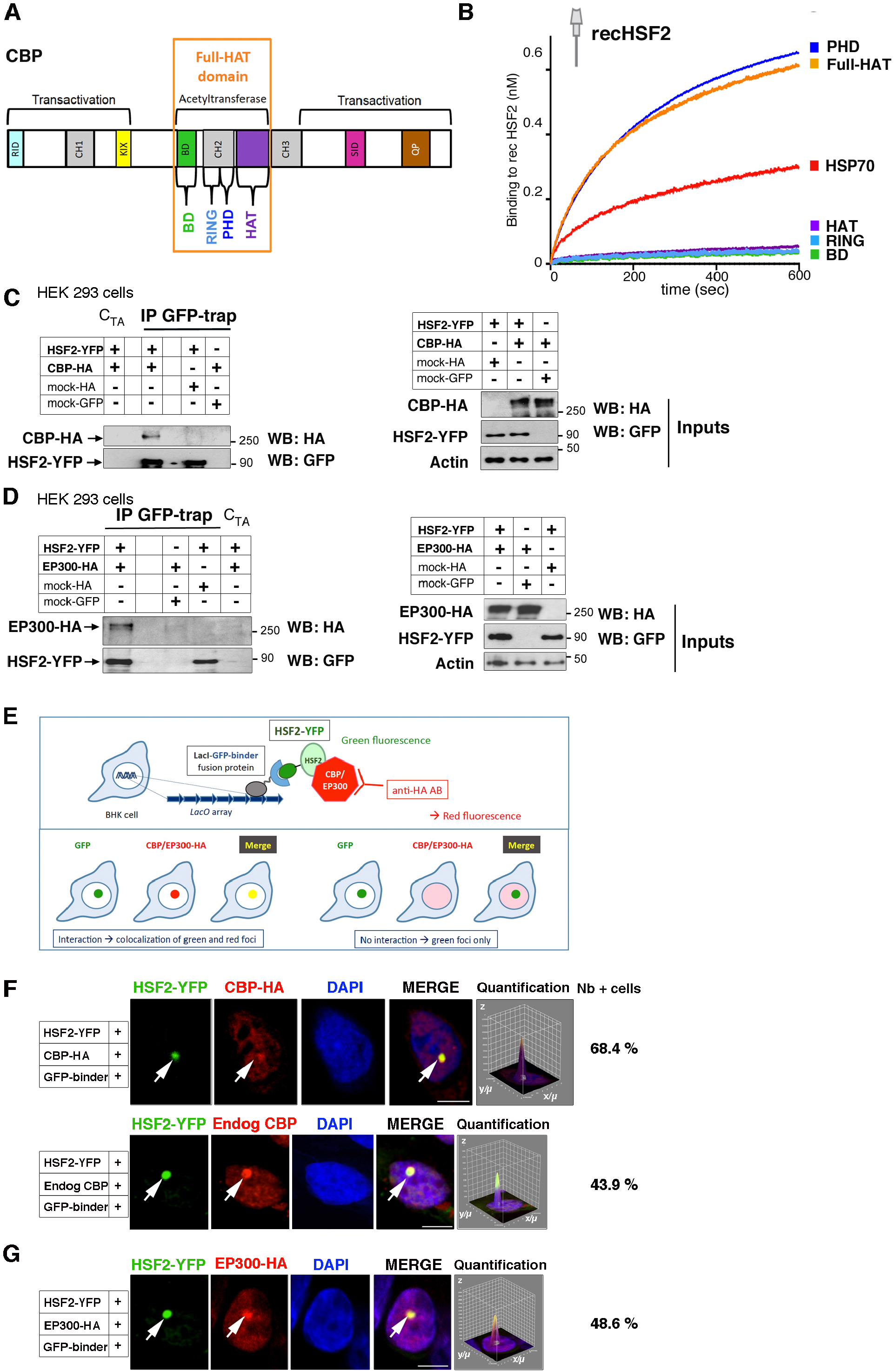
HSF2 interacts with CBP and EP300 in normal conditions. **(A) *Schematic representation of CBP protein domains***. The ability of CBP to bind a very large number of proteins is mediated by several conserved protein binding domains, including the nuclear receptor interaction domain (RID), the cysteine/histidine-rich region 1 (CH1), KIX, Bromodomain (BD), PHD, CH2, HAT, CH3, SID (steroid receptor co-activator-1 interaction domain), and the nuclear coactivator binding domain NCBD (not illustrated here) and QP (Glutamine- and proline-rich domain; Dancy and Cole, 2015; Dyson and Wright 2016). **(B)** Biolayer interferometry measurement of CBP binding to recombinant HSF2. Binding of different His-tagged domains of CBP to immobilized biotinylated recombinant HSF2 on streptavidin sensor tips (recHSF2): the catalytic full domain Full-HAT, or one of the following subdomains: PHD, HAT, RING, or Bromodomain (BD). HSP70 was used as a positive control for binding to HSF2 (Tang et al., 2016; see Figure S3A for determination of the Kd). **(C) *The ectopically expressed YFP-HSF2 protein interacts with exogenous HA-CBP.*** HEK 293 cells were transfected with combinations HSF2-YFP and CBP-HA, or mock-HA (or mock-GFP) constructs. (Left panels) HSF2-YFP was immunoprecipitated using anti-GFP-trap antibody (IP GFP-Trap) and co-immunoprecipitated CBP protein was detected by using anti-HA antibody. The immunoprecipitated HSF2 was detected using an anti-GFP antibody. (Right panels) The total amounts of exogenous HSF2 and CBP proteins in the input samples were detected with anti-GFP and anti-HA antibodies, respectively (inputs). Actin, loading control. Representative immunoblots (n=3 experiments). **(D) *The ectopically expressed YFP-HSF2 protein interacts with exogenous EP300.*** As in (**C**) except that HEK 293 cells were transfected with an EP300-HA construct. Representative immunoblots (n= 3 experiments). **(E)** Principle of the fluorescent-3-hybrid (F3H) assay. **(F) *F3H assay for the visualization of interaction between HSF2-YFP and exogenous CBP-HA, or HSF2-YFP and endogenous CBP*** (Upper and lower panels, respectively). (Left panels) Confocal sections of BHK cells carrying a stably integrated Lac-operator array that were triple transfected with *LacI* fused to the GFP-binder, HSF2-YFP, and CBP-HA constructs. Exogenous and endogenous CBP was detected using an anti-HA or an anti-CBP antibody, respectively (red signal). Chromatin was counterstained using DAPI. All the experiments involving negative controls are shown in Figure S3C,D. White arrows point out localization of HSF2-YFP and CBP-HA at the *LacO* spot. Scale bar, 10 μm. (Right panels) Graphs represent the quantification of the intensity of the two fluorescence signals, visualizing the co-localization of HSF2-YFP and CBP-HA signals to the *LacO* array (x/µ; y/µ; z, signal intensity in arbitrary units). “Number (Nb) of + cells”: quantification of the percentage of cells showing co-recruitment of YFP-HSF2 and CBP-HA or endogenous CBP, in the *Lac0* array. Representative images, n=3 independent experiments for CBP-HA and endogenous CBP. **(G) *F3H assay for the visualization of interaction between HSF2-YFP and exogenous EP300-HA*.** As in Figure 3F, except that exogenous EP300-HA was detected using an anti-HA antibody (red signal). All the negative controls are shown in Figure S3E. Representative images, n=4 independent experiments. Scale Bar, 10 µM.

### HSF2 is a *bona fide* substrate of the core catalytic domain of CBP

Prompted by the finding that a catalytically active CBP is necessary for HSF2 acetylation, we examined whether HSF2 could bind to the core catalytic domain of CBP. The CBP central core catalytic region (“Full-HAT”; Figure 3A) contains the Bromodomain BD, the cysteine/histidine-rich region CH2, and the HAT domain and allows the coupling of substrate recognition and histone/lysine acetyltransferase activity (as in EP300; Delvecchio et al., 2013; Dancy and Cole, 2015; Dyson and Wright 2016). The CH2, in particular, contains a RING domain and a PHD (plant homeodomain; Park et al., 2013). With biolayer interferometry, we observed that the recombinant Full-HAT domain directly interacted with immobilized biotinylated recombinant full-length HSF2 (Figure 3B). Within this region, the recombinant PHD domain, but not the HAT, RING or BD domain, was able to interact with HSF2, in a similar manner as the “Full-HAT” domain (Figure 3B). Interestingly, the interaction of HSF2 with the Full-HAT or the PHD domain was more efficient than with HSP70, which has been reported to interact with HSF2 (Huttlin et al., 2015; Tang et al., 2016). As expected, it is likely that the interaction between HSF2 and the catalytic HAT domain was too transient to be captured in these experiments, because the HAT domain needs other CBP domains to interact with its substrates, including the PHD (Aasland et al., 1995; Bordoli et al., 2001; Kalkhoven et al, 2002). We determined that the K_D_ of HSF2 interaction with the CBP Full-HAT domain was 1.003E^-09^ M (+/-2.343E^-11^; R2= 0.988488; Figure S3A). Our data on HSF2 acetylation and interaction with the Full-HAT domain of CBP, including PHD therein, strongly suggest that HSF2 is a *bona fide* substrate of CBP, and potentially also that of EP300, since their HAT domains display 86% identity.

### HSF2 interacts with CBP and EP300 *via* its oligomerisation HR-A/B domain

Our studies in human brain organoids and mouse cortices indicated that endogenous HSF2 and CBP/EP300 could interact (Figure 1). Similar results were observed in the murine neuroblastoma N2A cells (Figure S3B). We started to dissect the mode of anchorage between these proteins. We first tested whether exogenous proteins, tagged-HSF2 and -CBP/-EP300 could interact. Using HEK 293 cells in GFP-Trap assay, we showed that CBP-HA or EP300-HA co-immunoprecipitated with HSF2-YFP (Figure 3C,D). Second, we confirmed that interaction between these tagged proteins occurred *in cellulo* by an independent imaging technique, the fluorescent three-hybrid assay (F3H; Figure 3E; Herce et al., 2013), using GFP-binder and HSF2-YFP, together with CBP-HA or EP300-HA. In negative control experiments, GFP-binder, which was recruited to the *Lac0p* array locus, was unable to recruit endogenous CBP, CBP-HA, or EP300-HA (Figure S3C-E). Similarly, HSF2-YFP was unable to locate at the *Lac0p* array locus in the absence of GFP-binder (Figure S3C-E). Upon co-transfection with HSF2-YFP, CBP-HA or EP300-HA expression resulted in the formation of a red spot in the nucleus, showing co-recruitment to the HSF2-YFP focus (green spot), in 68.4% and 48.6% of the cells, respectively (Figure 3F, upper panels and 3G). The abundance of CBP in BHK cells allowed us to detect the co-recruitment of HSF2-YFP with endogenous CBP in 43.9% of these cells (Figure 3F, lower panels).

Having characterized the interaction between exogenous tagged proteins, we then asked by which specific domains the anchorage between HSF2 and CBP was facilitated. To determine which HSF2 domains were important for its interaction with CBP, we expressed Flag-HSF2 deletion mutants of different domains. We showed that the deletion of HR-A/B domain (but not of the DBD) led to a marked decrease in HSF2 acetylation (Figure S4A-C; see red arrow). These results were in line with our findings that the major acetylated lysine residues reside within the HR-A/B domain. The deletion of the TAD (transcription activation domain; Jaeger et al., 2016) also resulted in decreased acetylation of the Flag-HSF2 (Figure S4A-C) and was associated with decreased interaction with CBP (Figure S4D). This domain may be important for interaction between CBP and HSF2, since the multivalent interactions between CBP/EP300 and many transcription factors generally involve their TADs (Lee et al., 2009; Wang et al., 2012; reviewed in Thakur *et al*., 2013).

### The presence of KIX motifs in the HSF2 HR-A/B domain promotes binding to the CBP KIX domain

CBP/EP300 interacts with many transcription factors *via* different binding sites, including the KIX domain (kinase-inducible domain interacting domain; Figure 3A, left). The KIX domain contains two distinct binding sites that are able to recognize the “ΦXXΦΦ” KIX motif, where “Φ” is a hydrophobic residue, and “X” is any amino acid residue (Radhakrishnan et al., 1997; Kobayashi et al., 1997; Lee et al., 2009; Zor et al., 2004). Importantly, we identified several conserved, overlapping and juxtaposed KIX motifs in the HR-A/B domain of HSF2 (Figure 4A). We modeled the interaction between the HSF2 HR-A/B KIX motifs and the CBP KIX domain. Based on sequence similarities between the HR-A/B domain, lipoprotein Lpp56, and the transcription factors GCN4, ATF2, and PTRF (Figure S4E; see Experimental procedures), we first developed a structural model of the HSF2 trimeric, triple coiled-coil, HR-A/B domain (Figure S4F; Jaeger et al., 2016). Second, we investigated the possibility of interactions of the KIX recognition motifs in the HR-A/B region with the CBP KIX domain. Best poses suggested that the HR-A/B KIX motif region contacted the so-called “c-Myb surface” within the KIX domain (Figure 4B, Thakur *et al*., 2013), thereby proposing a close interaction of the HSF2 KIX motifs with the tyrosine residue Y650 of CBP (Figure 4C; Figure S4G(b)). We next examined the impact of *in silico* mutations of the K177, K180, F181, V183 residues, which are present within the KIX motifs of HSF2 and involved in the contact with CBP (Figure 4D). Either K177A or Q180A mutation within the HSF2 KIX motifs disrupted HSF2-KIX domain interaction (Figure S4G (e,f); Table S3), in contrast to either F181A or V183A mutation (Figure S4G(c,d)). Finally, we assessed the impact of *in silico* mutation of the Y650 amino acid of the CBP KIX domain, a residue mutated in RSTS patients (Figure 4E). Interestingly, the *in silico* mutation Y650A in CBP profoundly decreased the probability of interaction of the HSF2 KIX motifs with the KIX domain (Figure 4F, upper panel; and Figure S4G (b); Table S3). Using recombinant proteins, we verified that HSF2 directly interacted with the CBP KIX domain in *in vitro* co-immunoprecipitation experiments (Figure 4B) and we confirmed that the Y650A mutation disrupted HSF2 and KIX interaction (Figure 4G). Thereby, we identify Y650 as a residue critical for interaction between the KIX domain of CBP and the KIX motifs within the HSF2 oligomerisation domain.

**Figure 4.**
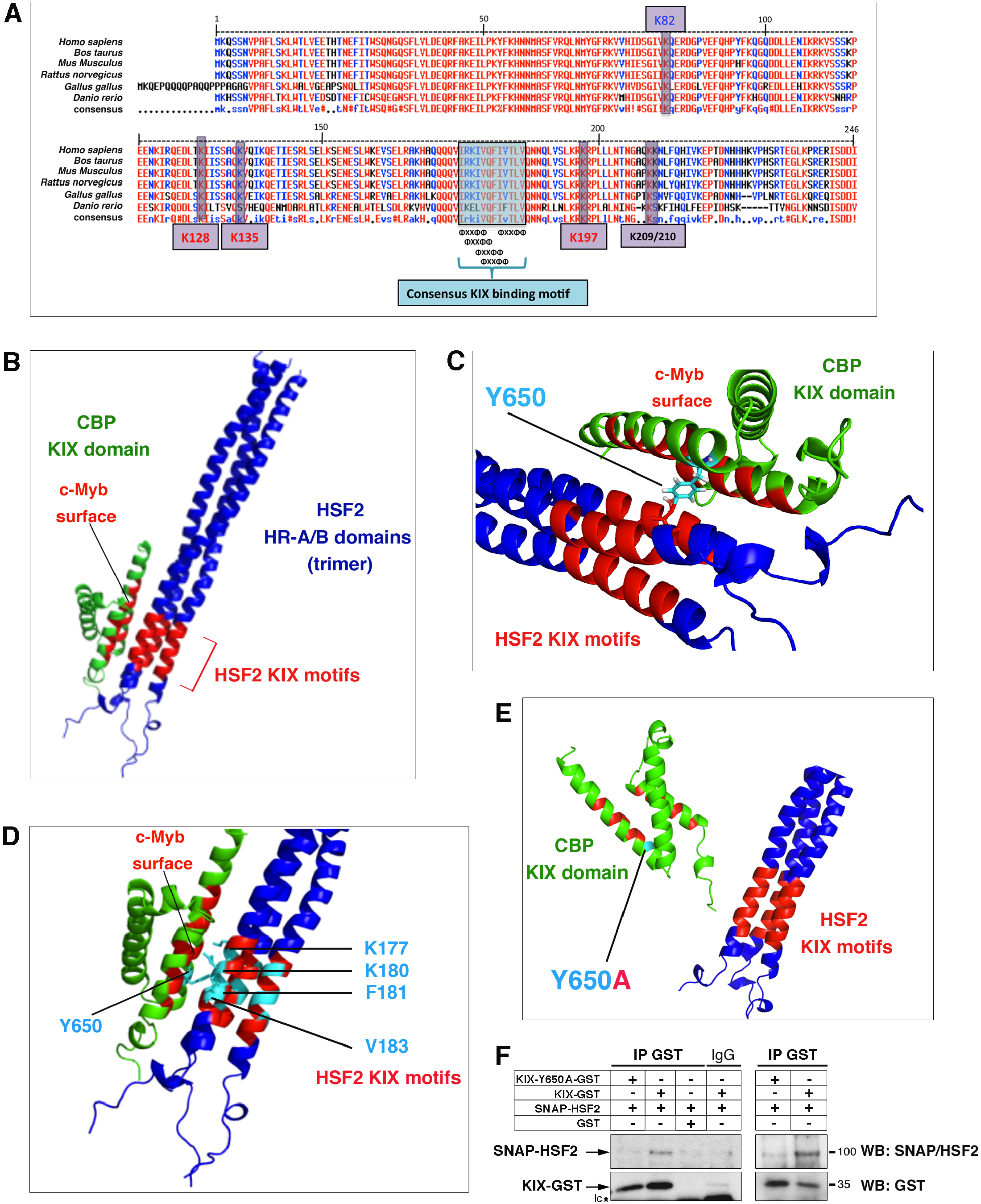
Modelling of CBP and HSF2 interaction. **(A) *Schematic representation of the KIX-binding motifs located in the HSF2 HR-A/B region*.** Conserved KIX-binding motif sequences (“ΦXXΦΦ”) are indicated (blue rectangle). The positions of the very conserved, major acetylated lysine residues are highlighted (purple rectangles): K82 located in the DBD (in blue), K128, K135, and K197 located in the HR-A/B domain (in red), and K209/K210, located downstream the HR-A/B (in black). **(B) *In silico* model structure of CBP KIX domain and HSF2 HR-A/B interaction**. Representation of the HSF2 HR-A/B domains of the HSF2 trimer (in blue), as a triple-coiled coil. The KIX recognition motifs of HSF2 are indicated in red. Representation of the KIX domain of CBP, a triple helical globular domain (in green). The c-Myb surface of the KIX-domain is indicated in red. **(C) *Magnification of the* in silico *representation illustrated in (C)*** showing the contact of the tyrosine residue Y650, located within the c-Myb surface of the CBP KIX domain, to the KIX recognition motifs located within the HSF2 HR-A/B domain. **(D) In silico *representation of the positioning of four residues located within the HSF2 KIX recognition motifs, and of Y650 within the CBP KIX domain that have been analyzed by* in silico *mutation*** (see (D-F) and Figure S4G). **(E) In silico *Y650A mutation disrupts interaction between the HSF2 KIX motifs and the CBP KIX domain*** *Firedock* analysis. (see also Figure S4G(b) for *Zdock* analysis and Table S3). **(F) In vitro *Y650A mutation disrupts interaction between the HSF2 KIX motifs and the CBP KIX domain*** (see also Figure S4G). The recombinant wild-type or mutated KIX-GST and SNAP-HSF2 proteins were produced in bacteria and reticulocyte lysates, respectively, and then subjected to an *in vitro* co-immunoprecipitation using an anti-GST antibody. The left and right panels correspond to two independent experiments. WB with anti-SNAP antibody (left upper panels) or with anti-HSF2 antibody (right upper panels).

### The acetylation of HSF2 governs its stability under non-stress conditions

To explore the functional impact of the CBP/EP300-mediated acetylation of HSF2, we inhibited CBP/EP300 activity in N2A cells, using the specific inhibitor C646 (Bowers et al., 2010; Dancy and Cole, 2015). The pharmacological inhibition of CBP/EP300 decreased the endogenous HSF2 protein levels, which was abolished by treatment with the proteasome inhibitor, MG132 (Figure 5A and Figure S5A). These results showed that the decrease in the HSF2 protein levels was dependent on the proteasomal activity, and that HSF2 was degraded when CBP/EP300 activity was inhibited (Figure 5A), thereby providing the first evidence for acetylation playing a regulatory role in HSF2 stability. To further investigate the role of acetylation in the regulation of HSF2 protein levels, we generated CRISPR/Cas9 *Hsf2*KO U2OS cell lines (2KO; Figure S5B,C) and measured the protein levels of exogenous wild-type HSF2 or HSF2 acetylation mutants, which mimic either constitutively acetylated (3KQ) or non-acetylated (3KR) HSF2 (Figure 5B-E). We first verified that the HSF2 WT, 3KQ and 3KR were expressed at comparable levels (Figure S5D), and capable of binding DNA *in vitro* (Figure S5E). Our *ex vivo* experiments also verified that they were also able to locate into specific subnuclear structures, called the nuclear stress bodies, nSBs (Jolly et al, 1997, 1999, 2004; Rizzi et al., 2004; Sandqvist et al., 2009), whose formation upon stress is associated to the recruitment of HSF1 and HSF2 on pericentromeric repeats, predominantly at the *SatIII* 9q12 locus (Figure S5F). To monitor the decay of a pre-existing pool of HSF2 molecules, we performed pulse-chase experiments using the SNAP-TAG technology (Bodor et al., 2012). A pool of SNAP-HSF2 molecules was covalently labeled by adding a fluorescent substrate to the cells. At t_0_, a blocking non-fluorescent substrate was added, quenching the incorporation of the fluorescent substrate to newly synthesized HSF2 molecules (Figure 5B), allowing us to measure the decay in the fluorescence intensity of the corresponding labeled HSF2 bands. When 2KO cells were transfected with wild-type SNAP-HSF2 (SNAP-HSF2 WT), a ∼50% decay in fluorescence intensity of the corresponding bands was observed within 5 hours (Figure 5C and D). Preventing HSF2 acetylation (SNAP-HSF2 3KR) resulted in a similar decay (Figure 5C and D). In contrast, mimicking acetylation with SNAP-HSF2 3KQ protected HSF2 from decay (Figure 5C and D). Of note, proteasome inhibition with MG132 prevented the decrease in SNAP-HSF2 WT and 3KR fluorescent intensity (Figure 5E). Moreover, we observed that mimicking the acetylation of HSF2 by expressing Myc-HSF2 3KQ limited the poly-ubiquitination of HSF2, when compared to HEK 293 cells expressing either WT or 3KR HSF2 (Figure 5F). (Figure 5F). Altogether these experiments demonstrate that HSF2 acetylation prevents the proteasomal degradation of HSF2.

**Figure 5.**
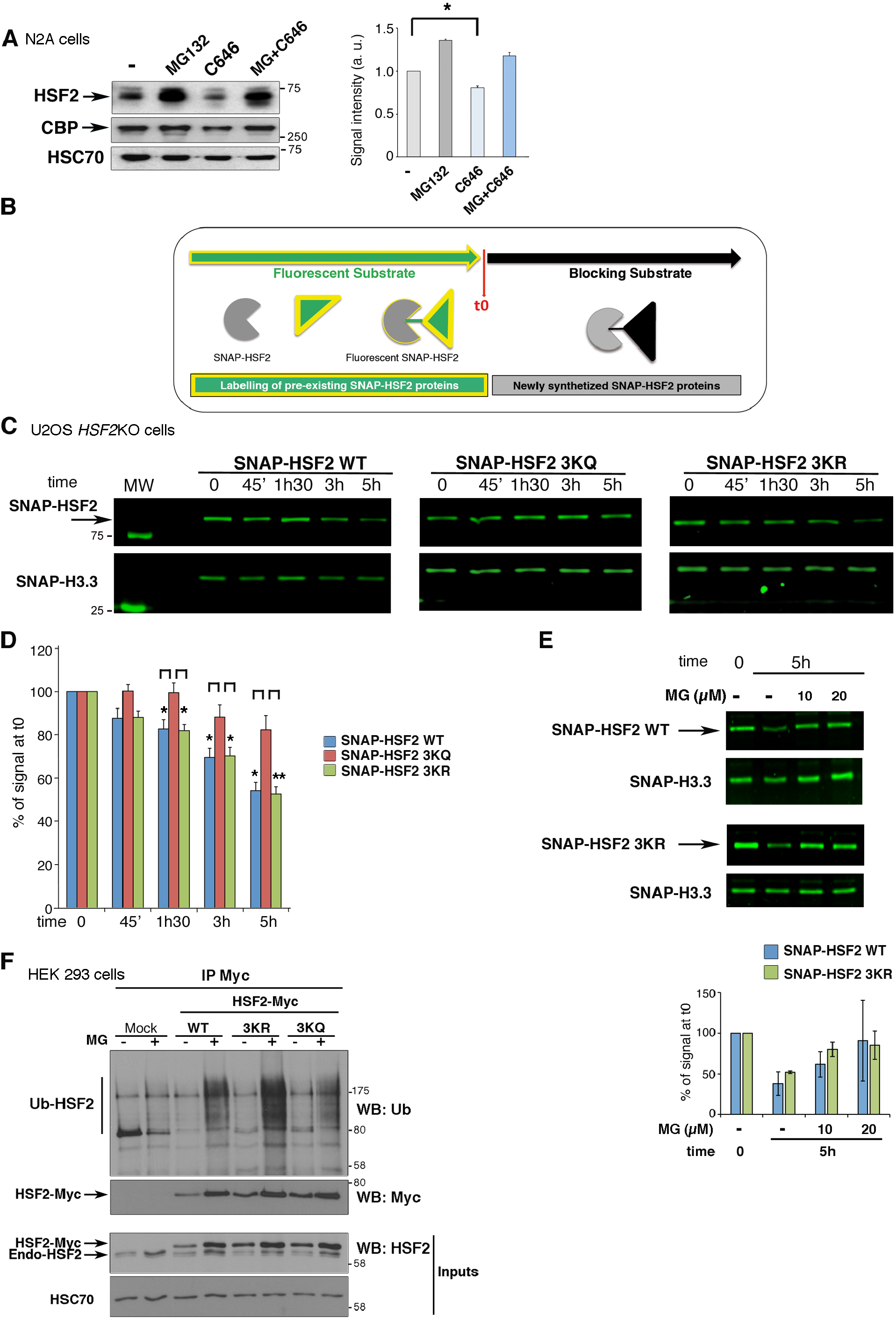
Impact of preventing or mimicking acetylation of lysine residues K128, K135, and K197 on HSF2 protein stability. **(A) *Inhibiting CBP/EP300 decreases HSF2 protein levels, a process counteracted by proteasome inhibition.*** (Left panels) Representative immunoblot of HSF2 and CBP levels upon treatment of N2A cells with the CBP/EP300 inhibitor C646 (40µM for 4 h) and/or with the proteasome inhibitor MG132 (20 µM for 6h). n = 4 independent experiments. (Right panel) Graph corresponding to quantification of the HSF2 signal intensity relative to the vehicle-treated samples (-) and normalized to the loading control HSC70. Standard deviation is indicated. * p<0.05 (relative to non-normalized data). **(B) *Schematic representation of the principle of SNAP-TAG-based pulse-chase experiments***. Cells expressing SNAP-tagged HSF2 are incubated in the presence of a fluorescent substrate, which, at a given time (t_0_), covalently labels the pool of SNAP-HSF2 molecules present in the cell. The addition of a non-fluorescent, blocking substrate prevents further labelling of the newly synthesized HSF2 molecules. It allows the measurement of the decay in the fluorescent signal corresponding to SNAP-HSF2 molecules covalently bound by the fluorescent substrate, thereby allowing an estimation of the decay in HSF2 protein levels. **(C) *Combined mutations of lysine residues K128, K135, and K197 mimicking HSF2 acetylation (3KQ) slow down the decay of HSF2 protein levels, compared with HSF2 WT***. Representative gel analysis of decay in HSF2 protein levels, carrying 3KR or 3KQ mutations, and labelled by the fluorescent SNAP-substrate, in CRISPR-Cas9 *Hsf2*KO U2OS cells. SNAP-H3.3 is used as a loading control. **(D) *Quantification of the fluorescent signal*** corresponding to SNAP-HSF2 WT (blue), 3KQ (red), or 3KR (green), as a measure of SNAP-HSF2 protein decay, relative to the control samples (t0) and normalized to the loading control H3.3. n=7 independent experiments. Standard deviation. * p<0.05;** p<0.01. **(E) *The decrease in SNAP-HSF2 WT and 3KR protein levels depends on proteasome activity***. (Upper panels) as in (**C**), but cells were pre-treated with 10 or 20 µM MG132 for 6 h. n=2 independent experiments. (Lower panels) quantification as in (D). **(F) *The HSF2 3KR mutation, favours HSF2 poly-ubiquitination, whereas 3KQ does not*.** HEK 293 cells were co-transfected with Myc-HSF2 WT, 3KR, or 3KQ, and treated or not with the proteasome inhibitor, MG132 (MG; 20 µM for 6 h). HSF2 was immunoprecipitated using anti-Myc antibody and its poly-ubiquitination status was analysed by WB, using an anti-ubiquitin (Ub) antibody. The protein amounts in the input samples were detected with antibodies against HSF2. HSC70, loading control. (n=3 independent experiments).

### HDAC1 is involved in the destabilization of the HSF2 protein under non-stress and stress conditions

To identify the enzymes that could function as deacetylases for HSF2, we performed an unbiased screen for HSF2 binding protein partners, using a double-affinity TAP-TAG approach (Bürkstümmer et al., 2006; Figure S6A-C). For this purpose, we generated a HeLa-S3 cell line expressing double-tagged HSF2 (or transfected with the empty vector as a negative control) and analyzed nuclear extracts by MS. We identified HDAC1 as one of the protein partners of HSF2 (Figure 6A). In addition to HSF2, we found nucleoporin Nup62 (Figure 6A), a known HSF2 partner that served as a positive control for the quality of our TAP-TAG/MS analysis (Yoshima et al., 1997). We also performed immunoprecipitation of HSF2 in extracts from the mouse E17 cortices, followed by MS analysis, and found both HDAC1 and HDAC2 as HSF2 partners (Figure S6D). Using the F3H approach (Figure 3E; Figure 6B and Figure S6E), and co-immunoprecipitation in GFP-Trap assays (Figure 6C), we confirmed the interaction between HSF2 and HDAC1 in mammalian cell lines. We then evaluated the impact of HDAC1 and other Class I HDACs on the acetylation of HSF2 by expressing CBP-HA in HEK 293 cells (Figure 6D and Figure S6F). HDAC1 overexpression resulted in marked reduction in HSF2 acetylation levels, whereas HDAC2 and HDAC3 had only limited effects, no effect of HDAC8 was detectable (Figure 6D and Figure S6F).

**Figure 6.**
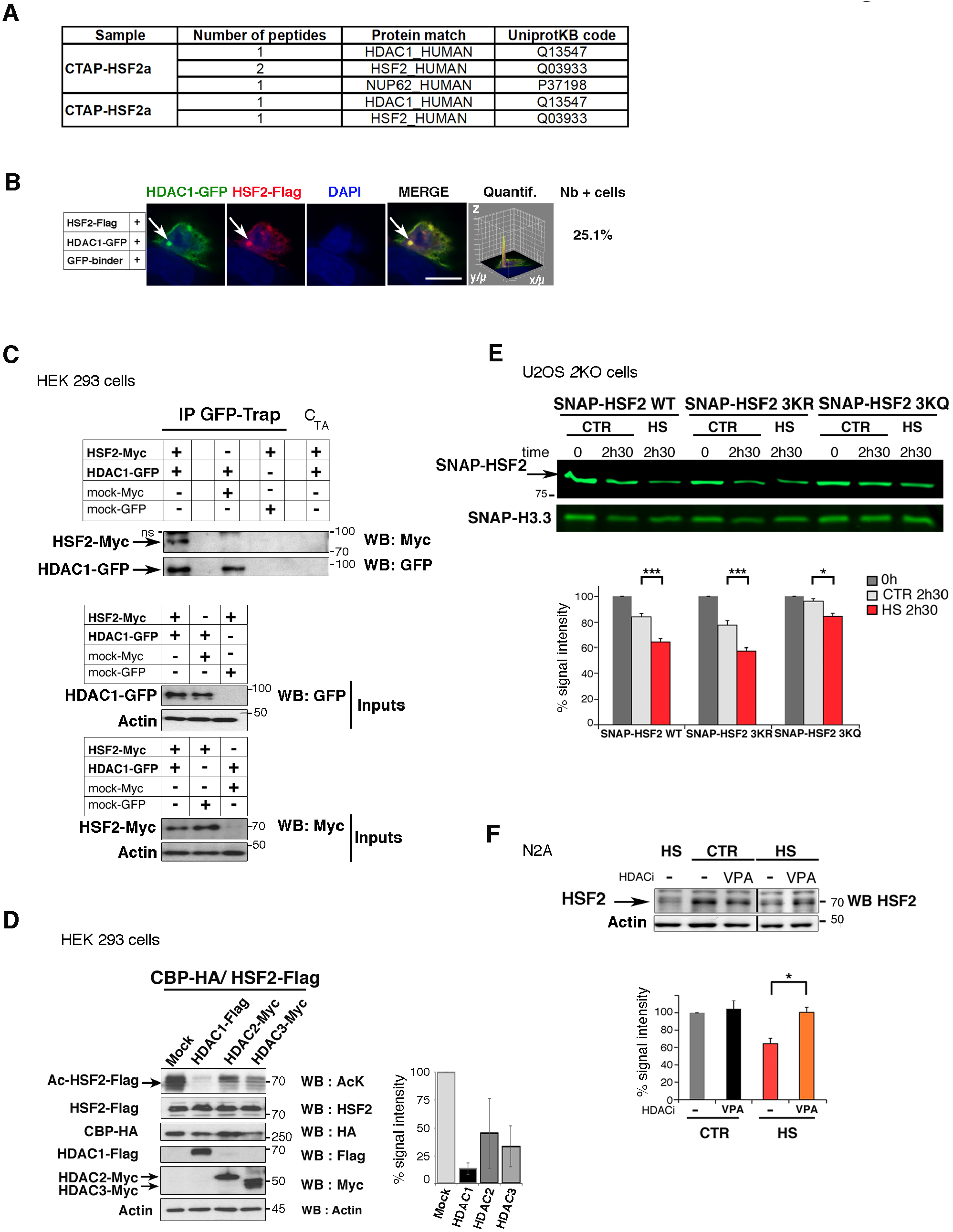
Impact of HDAC1 on HSF2 levels under non-stressed and stress conditions. **(A) *Identification of HDAC1 as a HSF2 protein partner in TAP-TAG/MS analysis in HeLa-S3 cells***. After sequential immunoprecipitation of nuclear extracts of HeLa-S3 expressing CTAP-HSF2α and CTAP-HSF2β, using two tags (G-protein and Streptavidin-binding peptide; see Figure S6A), eluates were analyzed by MS. The number of unique peptides from each identified protein and their UniProt Knowledgebase (UniProtKB) codes are indicated. **(B) *Interaction between ectopically expressed HDAC1-GFP and HSF2-Flag in F3H assays***. As in Figure 3F, except that BHK cells were triple transfected with *LacI* fused to the GFP-binder, HDAC1-GFP, and HSF2-Flag constructs. Chromatin was counterstained using DAPI. All the negative controls are shown in Figure S6E. n=3 independent experiments. Scale Bar, 10 µM. **(C) *The ectopically expressed exogenous HDAC1 interacts with HSF2.*** (Upper panel) GFP-Trap co-immunoprecipitation of HDAC1-GFP and HSF2-Myc in transfected HEK 293 cell extracts. (Middle and lower panels) Immunoblot showing total HDAC1 (WB GFP) or HSF2 levels (WB Myc) in inputs, respectively. n=2 independent experiments. Actin was used as a loading control. ns: non-specific band. **(D) *Overexpression of HDAC1 markedly reduces the acetylation of HSF2 by CBP*.** HEK 293 cells were transfected by the following constructs: CBP-HA and HSF2-Flag, and HDAC1-Flag, HDAC2-Myc, or HDAC3-Myc and the acetylation status of the HSF2-Flag protein was checked by using an anti-AcK antibody. n=5 independent experiments. Actin was used as a loading control. **(E) *The stability of HSF2 3KQ is increased upon HS, compared with HSF2 WT or HSF2 3KR***. (Upper panel) Representative gel analysis of HSF2 protein decay in a SNAP-TAG pulse-chase experiment. As in Figure 5C, except that cells were submitted to HS at 42°C for the indicated times. (Lower panel) Quantification of the fluorescent signal intensity, relative to the control samples (t0) and normalized to the loading control H3.3. n=7 independent experiments. Standard deviation. * p<0.05; *** p<0.001. SNAP-H3.3 was used as a loading control. **(F) *Class I HDAC inhibitor VPA prevents the decrease in endogenous HSF2 protein levels induced by HS***. N2A cells were pretreated or not with 1 mM valproic acic (VPA) for 3 h and subjected to HS 42°C for 2h30. (Upper panel) Representative immunoblot. (Lower panel) Quantification on the signal intensity normalized to actin levels (n=4 independent experiments). * p<0.05.

Because heat shock (HS) provokes the degradation of the HSF2 (Ahlskog et al., 2010), we thus analyzed the impact of acetylation on the heat-shock-induced decay of HSF2, using the SNAP-TAG technology. Mimicking the acetylation of the three major acetylated lysine residues mitigated the decay of fluorescence intensity of SNAP-HSF2 3KQ induced by HS, compared to SNAP-HSF2 WT or 3KR (Figure 6E). The impact of HDAC inhibition on endogenous HSF2 was investigated in N2A cells. We first verified that HS was able to induce HSF2 decay also in N2A cells, although it occurred at a slower rate than in HeLa or HEK 293 cells (Figure S6G; Ahlskog et al., 2010). Treatment with 1 mM of the Class I inhibitor VPA dampened the decline in HSF2 protein levels in N2A cells exposed to HS (Figure 6F and Figure S6H). This indicates that Class I HDAC activity participates to the degradation of HSF2 upon HS, likely through HSF2 deacetylation. We then verified that HS increased HSF2 poly-ubiquitination in HEK293 cells, as previously reported (Ahlskog et al. 2010; Figure S6I, mock transfection). To investigate whether HDAC1 could favor HSF2 poly-ubiquitination, likely through HSF2 deacetylation, we examined the impact of overexpression of a dominant-negative form of HDAC1 on HSF2 ubiquitination. We showed that, indeed, the increase in HSF2 poly-ubiquitination upon HS was mitigated in HEK 293 cells transfected with dominant-negative HDAC1 (Suppl. Figure S6I). As a whole, our results support a role of HDAC1 (and possibly other Class I HDACs) in the destabilization of HSF2 under normal and stress conditions, through HSF2 poly-ubiquitination and proteasomal degradation.

### Declined HSF2 protein levels in the Rubinstein-Taybi Syndrome (RSTS)

To determine the functional impact of CBP and EP300 on HSF2 levels in a pathological context, we compared the amounts of HSF2 protein in cells derived from either healthy donors (HD) or RSTS patients, which are characterized by autosomal-dominant (heterozygous) mutations in the *CBP* or *EP300* genes (see Figure S7A for a description of the mutations). We used human primary skin fibroblasts (hPSFs), at early passages to avoid putative compensation processes during *ex vivo* culture (see Experimental Procedures). We verified the effect of mutated *CBP* or *EP300* in RSTS hPSFs by showing that the amount of acetylated lysine residue K27 in histone H3 (AcH3K27) was reduced in both cases, compared to healthy donors (patient 1 [P1] and patient 2 [P2], respectively, Figure S7B). We observed that HSF2 protein levels were markedly decreased in hPSFs from RSTS patients carrying either *CBP* or *EP300* mutations (Figure 7A-C). HSF2 levels were restored to levels that were comparable to those of healthy donors (HD) when these hPSFs had been treated with the proteasome inhibitor MG132 (Figure 7A and B). However, Class I HDAC inhibition by VPA could not restore the HSF2 levels in RSTS*_CBP_* or RSTS*_EP300_* hPSFs, although HD and RSTS cells displayed similar levels of HDAC1 (Figure 7B; Figure S7C-E). Notably, the stability of HSF2 was impaired both in the RSTS*_CBP_* patient carrying a mutation in the Full-HAT domain of CBP and in the RSTS*_EP300_* patient carrying a deletion of the KIX domain of EP300. This finding suggests that both domains are required for the regulation of HSF2 stability, which is in line with our results (Figure 3–5). Altogether these results demonstrate that the proteasomal turnover of HSF2 is increased in RSTS hPSFs, carrying mutated either *EP300* or *CBP*, thereby strongly indicating that EP300 and CBP are key regulators of HSF2 protein stability.

**Figure 7.**
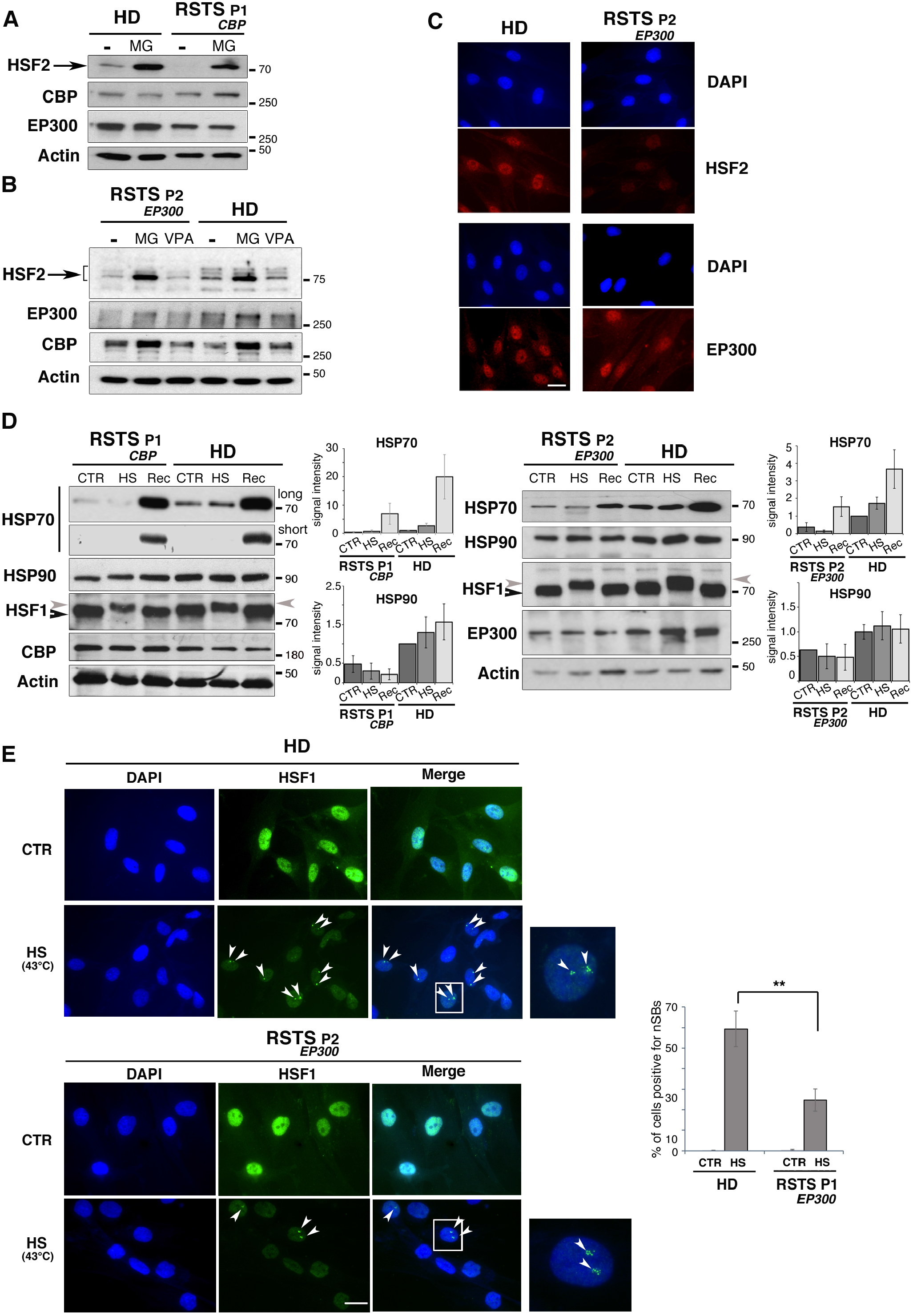
Altered HSF2 protein levels and dysregulated stress response in cells from RSTS patients. **(A) *HSF2 levels are reduced in RSTS_CBP_ hPSFs (patient P1), but restored in the presence of the proteasome inhibitor MG132***. See Figure S7A for the description of the patient’s mutation. Representative immunoblots. Cells were treated with 20 µM MG132 (6 h) and subjected to immunoblot analysis. n=3 experiments. Actin was used as a loading control. **(B) *HSF2 levels are reduced in RSTS_EP300_ hPSFs (patient P2), but restored in the presence of the proteasome inhibitor MG132.*** See Figure S7A for the description of the patient’s mutation. Cells were treated with 20µM MG132 (6 h) or 1mM of the HDAC inhibitor VPA (3 h) and subjected to immunoblot analysis. In contrast to MG132, the HDAC inhibitor VPA does not restores HSF2 levels. n=3 experiments. **(C) *HSF2 staining is reduced in RSTS_EP300_ hPSFs.*** Representative immunofluorescence experiments. n=3 experiments. Scale Bar, 10 µM. **(D) *Reduced HSP basal levels and induction by heat shock in RSTS, compared to HD hPSFs.*** (Left panels) Representative immunoblots from ***RSTS_CBP_*** [P1]. CTR, control conditions; HS, heat shock conditions (1 h at 42°C); Rec, recovery at 37°C for 2 h. Two exposure times for HSP70, long and short. Quantification of the HSP70 and HSP90β (HPS90AB1) signal intensity in immunoblots, normalized to actin. (Right panels) as in left panels, but from ***RSTS*_EP300_** [P2]. n=3 experiments. The grey arrow heads point to the hyperphosphorylated and thereby shifted HSF1 band. **(E) *Altered formation of nSBs by HS in RSTS*_EP300_ *hPSFs.*** (Left panels) Representative pictures of cells in control (CTR) or heat shock conditions (HS at 43°C for 1 h; arrowheads point to nuclear stress bodies [nSBs]). The white rectangles point out two examples of the magnified cells, positive for nSBs. (Right panel) Quantification of the percentage of fibroblasts positive for nSBs, from 100 – 150 cells in n=3 different experiments. RSTS *hPSFs* were compared to HD *hPSF.* Standard deviation*s*. ** p<0.01. Scale Bar, 10 µM.

### Impaired heat shock response in RSTS cells

HSF1 is the essential driver of the acute heat shock response (HSR) in mammals (McMillan et al., 1998). Although dispensable for the HSR in most cellular contexts (McMillan et al., 1998), HSF2 acts as a fine tuner of the HSR (Östling et al., 2007; Elsing et al., 2014), which determines the magnitude to which the applied heat stress induces *HSP* gene expression. Therefore, we evaluated the ability of RSTS cells to mount a HSR. In the absence of heat stress, we observed that RSTS hPSFs displayed lower amounts of HSP70 and HSP90 than their HD counterparts (Figure 7D). Furthermore, RSTS hPSFs exhibited limited capacity in inducing HSP70 accumulation upon HS and during the recovery phase from heat stress (Figure 7D). Importantly, this limited induction did not result from impairment of HSF1 activation, since HSF1 was activated by HS in RSTS hPSFs, as assessed by its slowed mobility shift in SDS-PAGE (see arrowheads in Figure 7D). This shift is a hallmark of HSF1 hyperphosphorylation, which, although not required for HSF1 activation, accompanies the induction of HSF1 transactivation potential (Sarge et al., 1993; Budzyński et al., 2015; reviewed in Anckar and Sistonen, 2011). As mentioned above, HSF1 and HSF2 do not only control the transcription of the *Hsp* genes in response to acute heat stress, but they also upregulate the transcription of *Sat III* 9q12 heterochromatin regions, where nSBs are formed. We therefore used nSBs as a read-out for assessing the HSR integrity in RSTS cells and observed that the stress-inducible formation of nSBs was reduced by more than 50% in RSTS*_EP300_* hPSFs when compared to their HD counterparts (Figure 7E at 43°C and S7F at 42°C). A similar reduction in the formation of nSBs was observed in RSTS lymphoblastoid cells (LBs; Figure S7G).

The functional importance of the regulation of HSF2 stability by CBP and EP300 is therefore highlighted in the RSTS context, under non-stress and stress conditions, which might constitute an interesting novel reading key for this complex disease.

## DISCUSSION

During the last decade, HSFs have been associated with a wide spectrum of pathophysiological conditions, and the specific roles of HSFs, either individually or in combination with each other or with other transcription factors, are of great biomedical interest. Especially, the mechanisms by which the expression levels of HSFs are regulated, in a context-dependent manner, have remained poorly understood. However, there is a wealth of documented cases where either excessive or insufficient HSF protein levels favor the development or progression of devastating diseases, including cancer, neurodevelopmental, and neurodegenerative disorders. In this study, we reveal a novel mechanism that regulates the stability of HSF2 protein. Using different cellular systems and pathophysiological conditions, we found that the KATs CBP and EP300 catalyze the acetylation of three highly conserved lysine residues K128, K135, and K197, located in the HR-A/B oligomerization domain, thereby contributing to the stability of the HSF2 protein. Moreover, we demonstrate the importance of this regulation both under normal and stress conditions, as well as in the pathological conditions of the Rubinstein-Taybi Syndrome (RSTS), which is a rare but highly detrimental disease.

### HSF2 is acetylated by CBP/EP300 in various contexts including brain development

In addition to ectopically/exogenously expressed proteins, HSF2 is acetylated by the overexpression of CBP or EP300 in cell systems. Importantly, we were able to detect the acetylation of the endogenous HSF2 protein in human and murine neural embryonic tissues and cell lines (Figure 1 and S1), showing that HSF2 acetylation is not restricted to one specific cell context. As does HSF2, CBP and EP300, and more generally HATs play significant roles during neurodevelopment (reviewed by Lopez-Atalaya et al., 2014). HSF2, CBP and EP300 exhibit similar expression patterns along the differentiation of human brain organoids, as they do in the developing mouse cortex. This is, to the best of our knowledge, the first report on the expression profiles of HSF2, CBP, and EP300 proteins in a model of the human developing brain as well as on the interactions between HSF2 and CBP/EP300. Our results point out the physiological importance of HSF2 acetylation and suggest that the acetylation of HSF2 might be a key event involved in the abundant expression and important role of HSF2 in cortical development. Accordingly, it is interesting to note that, our MS analysis revealed HDAC1 as an HSF2 partner in E17 cortices, at which stage the HSF2 protein levels are markedly dowregulated (Figure S7; El Fatimy et al., 2014).

### Mode of HSF2 interaction with CBP/EP300

In our attempt to determine the molecular and structural basis for interaction between HSF2 and the acetylating enzymes CBP and EP300, we first show that the full-length HSF2 protein interacts with the CBP core catalytic domain *in vitro*, confirming that HSF2 is a *bona fide* substrate of CBP. In addition, HSF2 strongly interacts with the PHD domain, located within the catalytic core of the CBP/EP300 proteins (Delvecchio et al., 2013). Interestingly, mutations in the PHD domain of CBP/EP300 have been identified in RSTS patients (Kalkhoven et al., 2003). Based on the cell-based analyses combined with the *in vitro* and *in silico* analyses, we found that the HR-A/B oligomerization domain, but not the DNA-binding domain, is necessary for the interaction of HSF2 with CBP and for HSF2 acetylation. Importantly, we show that the HR-A/B domain specifically interacts with the KIX domain of CBP/EP300.

The CBP and EP300 KIX domain serves as a docking site for the binding of many transcription factors and contributes to the properties of CBP/EP300 to act as a molecular bridge, stabilizing the interactions between specific transcription factors and the transcription machinery (Parker et al., 1996; reviewed in Thakur et al., 2014). According to our *in silico* analyses, the KIX-binding motifs that we identified in the HR-A/B oligomerization domain of HSF2 are necessary for its specific interaction with the KIX domain of CBP/EP300, and thus for HSF2 acetylation. In support of the close interaction between the HSF2 KIX recognition motifs and the CBP KIX domain, we show that the mutation of the tyrosine residue Y650 in the CBP KIX domain disrupts HSF2-CBP interaction in *in silico* and *in vitro* experiments. Likely a similar mode of interaction between the HSF2 KIX motif and the homologue of Y650 in EP300 can be expected (Kauppi et al., 2008). Interestingly, this mutation has been identified in RSTS, and associated with a severe neurodevelopmental phenotype (Spena et al., 2015b). Moreover, our *in silico* analyses indicate that the HR-A/B KIX motifs in HSF2 bind to the c-Myb site of the KIX domain. Indeed, the CBP or EP300 KIX domain can, simultaneously and in a cooperative manner, bind two polypeptide ligands (from two transcription factors), on two distinct surfaces, which have been historically called the “c-Myb” and the “MLL” (*Mixed Lineage Leukemia* protein) sites (Goto et al., 2002; Campbell and Lumb, 2002; reviewed in Thakur et al., 2014). This suggests that the binding of another transcription factor via the “MLL site” might potentially modulate the interaction between HSF2 and CBP, through its the KIX domain.

In addition to the HR-A/B oligomerization domain, we also identify the HSF2 TAD (Jaeger et al., 2016) as a potentially important domain for HSF2 interaction with CBP. This result is reminiscent of the TADs of other transcription factors that bind the KIX domain (reviewed in Thakur et al., 2014). Moreover, two different domains of the same transcription factor can simultaneously bind the KIX domain (Lee et al., 2009; Wang et al., 2012). We therefore hypothesize that HSF2 could simultaneously interact with CBP, through two distinct domains, the HR-A/B and the TAD domains. In addition, the dimeric or trimeric coil-coiled structure of HSF2 might also broadens the possibility of establishing multiple contacts with CBP (and most likely EP300), through KIX or other CBP domains, known to also interact with TADs, which paves the way for future studies.

### Dynamics of HSF2 acetylation by CBP/EP300, deacetylation by HDAC1, and degradation

Based on our data, the acetylation of HSF2 by CBP/EP300 limits its proteasomal degradation, which has been observed for other transcription factors, such as p53, STAT3, and HIF1alpha (Grossman, 2001; Jain et al., 2012; reviewed in Yang and Seto, 2008; Geng et al., 2012). However, acetylation does not seem to act by directly preventing the poly-ubiquitination of the three HSF2 lysine residues, K128, K135, and K197. Indeed, only combined mutations of these lysines to glutamines (3KQ), but not to arginines (3KR), prevent HSF2 proteasomal degradation. In addition, 3KQ mutation decreases HSF2 polyubiquitination, whereas 3KR does not (Figure 5). Previous proteome-wide quantitative analyses of the ubiquitin-modified protein have revealed that theubiquitination of HSF2 occurs on multiple residues spanning over the HSF2 protein, including K51, K151, K210 and K420, in addition to K128, K135, and K197. Most of these sites reside in the HR-A/B domain or its vicinity, suggesting a crosstalk between acetylation and ubiquitination (Kim et al., 2011; Wagner et al., 2011; Akimov et al., 2018; www.phosphosite.org). It is important to note that the published studies have not assessed the functional impact of these ubiquitination events on HSF2 turnover (reviewed by Gomez-Pastor et al., 2018). Moreover, we showed that mimicking the acetylation of the lysine residues K128, K135, and K198 limits the degradation of HSF2, also under heat shock conditions. In parallel, we identify HDAC1 as a major lysine deacetylase involved in HSF2 deacetylation and the control of HSF2 proteasomal degradation, both under basal and stress conditions, which may provide an explanation for the rapid degradation of HSF2 by APC/C in response to heat shock (Ahlskog et al., 2010).

Finally, future studies are warranted to determine whether other enzymes (HATs/KATs or HDACs) regulate HSF2 acetylation, deacetylation and thereby stability, and modify other lysine residues that we found acetylated in our MS analysis, as it is the case for HSF1 (Westerheide et al., 2009; Zelin et al., 2012; Raychaudhuri et al., 2014; reviewed by Miozzo et al., 2015).

### HSF2 destabilization and impaired heat shock response in RSTS syndrome

The accelerated turnover of the HSF2 protein in primary cells derived from the RSTS patients mutated either for *CBP* or *EP300* is counteracted by proteasome inhibition. This finding confirms the functional importance of HSF2 interaction with the two KAT3 family members, CBP and EP300, and provides further evidence for the role of acetylation on HSF2 stability. RSTS is a rare genetic, autosomal dominant neurodevelopmental disorder, characterized by intellectual disability, heart and skeleton malformations, and elevated susceptibility to infections as well as childhood cancers (Spena et al., 2015). Of the clinically diagnosed RSTS cases, the two identified genes mutated or deleted represent 60% for *CBP* and 8-10% for *EP300*. One mutated allele of *CBP* or *EP300* is sufficient to provoke this severely disabling disease. This is surprising given that, not only a wild-type copy of the affected gene, but also the two wild-type alleles encoding the other closely related KAT3 are present in the patients. The causal mutations in RSTS patients are extremely diverse and can affect protein expression, protein-protein binding or catalytic activity, which results in loss of specificity for interacting partners and/or substrates. The mutated *CBP (or EP300)* allele *via* a rupture of this subtle equilibrium could thereby exerts a “dominant-negative” effect and compromise compensation by the wild-type copies of the other KAT3 (Merk et al., 2018; Lopez-Atalaya et al., 2014). Accordingly, the presence of either a catalytically inactive *CBP* allele or an *EP300* allele, deleted for the KIX domain, is sufficient to influence the proteasomal turnover of HSF2 in RSTS cell system. These results suggest that the presence of the dominant-negative mutated allele impairs HSF2 acetylation. In addition, VPA was unable to restore the HSF2 levels in RSTS cells, likely because HSF2 is not acetylated in these cells, which reinforces the hypothesis that the integrity of EP300 and CBP function is critical for the control of HSF2 levels in cells derived from RSTS patients.

HSF2 is known to modulate the intensity of the heat shock response (HSR) by fine-tuning the expression of *HSP* genes and the formation of nSBs (Östling et al., 2007; Sandqvist et al., 2009). This occurs through the concomitant binding of HSF2 and HSF1, which involves the formation of heterotrimers, to the regulatory regions of the *HSP* genes and *SatIII* loci (Alastalo et al., 2003; Östling et al., 2007; Sandqvist et al., 2009). In the context of the RSTS model, HSF2 seems to be profoundly deregulated, whereas HSF1 is only slightly affected, and we find that the basal levels of HSP protein are reduced, including HSP70. In response to heat shock, the magnitude of HSP70 induction is clearly impaired, indicating that HSF2 is an important regulator of the HSR in the hPSFs cells, which is in line with our earlier results (Östling et al. 2007). Moreover, we also observe that the reduced HSF2 levels are associated to a decrease in the formation of nSBs in RSTS cells. It is therefore possible that the HSR is regulated at two different levels by the proteasome, providing an exquisite and sophisticated way to control cell proteostasis, *via* the stabilization of: 1) HSF1 in an EP300-dependent manner (at least in some cell systems; Raychaudhuri et al., 2014); and/or 2) HSF2, in a CBP- and EP300-dependent manner, as shown here in hPSFs. The delicate balance between these two arms of regulation, *i.e.* driven by HSF1 or HSF2, could be tipped depending on the cellular context, and potentially by the HSF1/HSF2 ratio (Östling et al., 2007; Sandqvist et al., 2009; Elsing et al., 2014). In conclusion, the repertoire of chaperones is altered in a chronic manner in RSTS cells already under physiological (unstressed) conditions, more extensively upon exposure to stress, and this alteration might confer the diseased cells vulnerability to proteostasis challenges.

The dysregulation of the HSF pathway in RSTS, including the markedly reduced HSF2 levels, might have several implications for this multifaceted disease. Indeed, RSTS is a neurodevelopmental disorder and HSF2, as a transcription factor involved in neurodevelopment both in normal and stress conditions, might contribute to the neurodevelopmental defects characteristic for RSTS. Strikingly, RSTS patients suffer from extreme vulnerability to airway infections, which is mainly due to defects in mounting a response to polysaccharides (Naimi et al., 2006; Herriot et al., 2016). Because the HSF pathway is involved in response to polysaccharides, inflammatory and immune responses, as well as lung protection against stress, its deregulation could also contribute to this aspect of the pathology (Xiao et al., 1999; Inouye et al., 2007; Wirth et al., 2003). Therefore, any imbalance in the delicate composition of the HSPs and other molecular chaperones under normal conditions and the triggering of HSF-driven stress-responses could profoundly influence this vulnerability to multiple severe health problems associated with the complex RSTS syndrome.

Rare diseases are in the center of growing interest based on the recent acceptance that they represent a global public health and economic problem. For example RSTS, despite its rarity (1:100,000 births) represents one on 300 patients institutionalized for intellectual disability (Spena et al., 2015). Moreover, they are conceptually considered as extreme components in the spectrum of a large diversity of diseases, and their study has the strong potential to highlight shared features in related common disorders, and thus of being transposable to other pathologies and deeply transform our ways to comprehend these pathologies. The functional impact of the regulation of HSF2 stability revealed in the context of RSTS might also represent an important reading key in related diseases, including neurodevelopmental disorders.

## EXPERIMENTAL PROCEDURES

### CONTACT FOR REAGENT AND RESOURCE SHARING

More detailed information and requests for resources and reagents should be directed to and will be fulfilled by the co-corresponding authors: Aurélie de THONEL, Lea SISTONEN, and Valérie MEZGER.

### Reagents and treatments

Proteasome inhibitor MG132 was used for 6 h (N2A, U20C_2KO) at a final concentration of 20μM. HDACs inhibitor VPA (Interchim, AYJ060) was used at 1 mM for 3 h. The HAT inhibitor C646 (Sigma-Aldrich; SML0002) was used at a final concentration of 20 or 40 μM for 4 h. For all chemicals, DMSO was used as vehicle (control).

Heat shock treatments were performed in water bath at 42 or 43°C for the indicated times.

### Table for antibodies

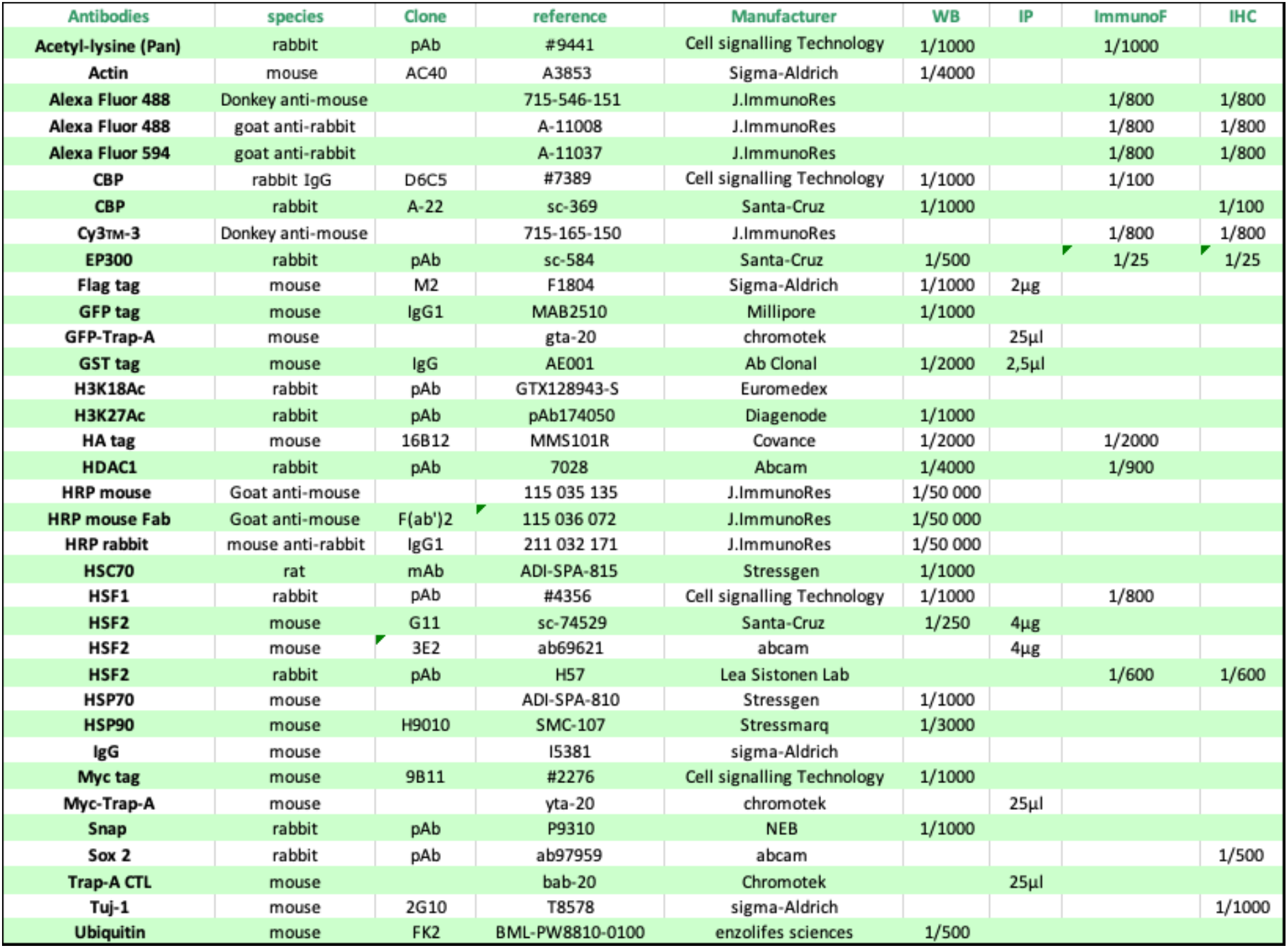

### Plasmid Table and constructs

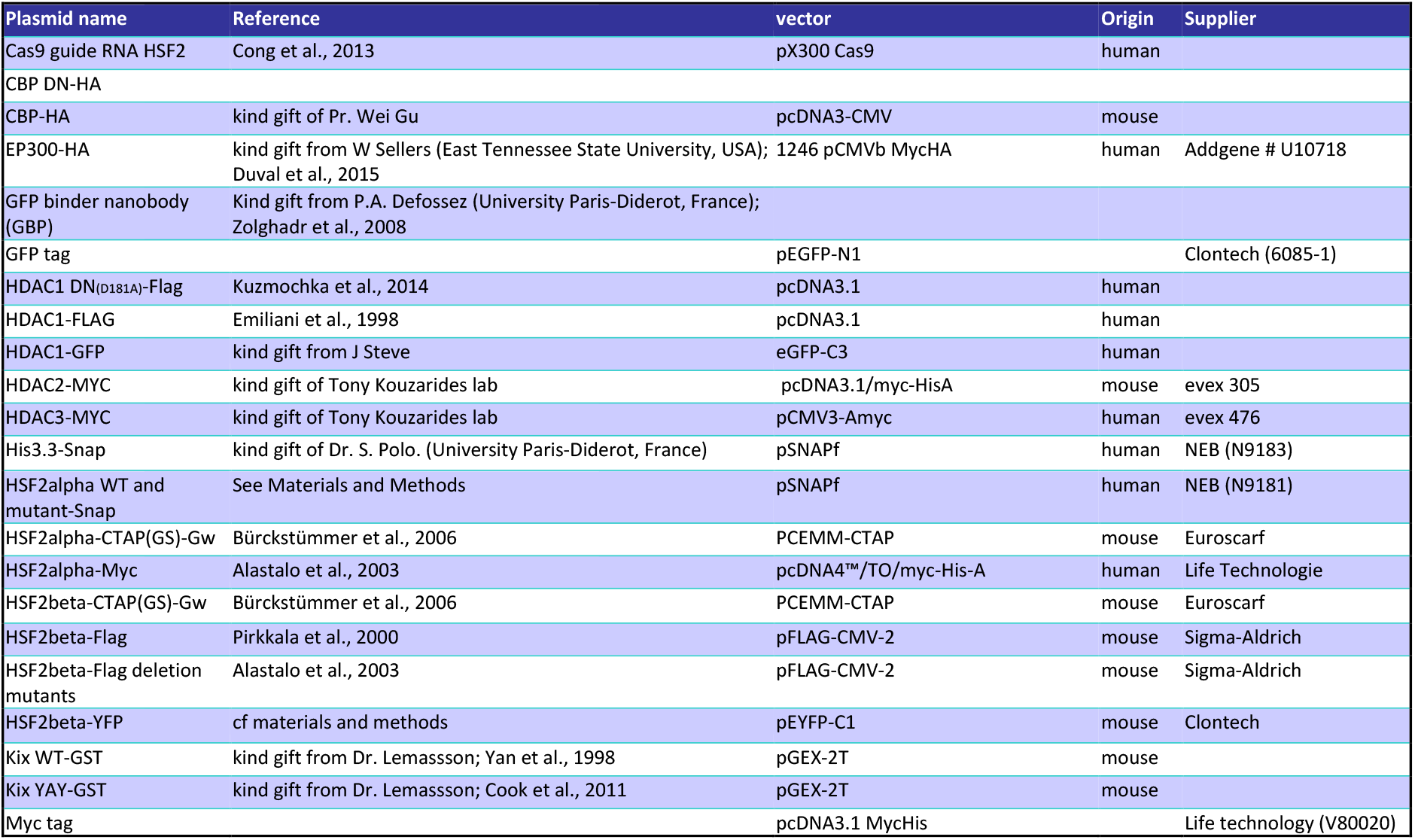

The human HSF2-Snap (WT/mutants) were constructed from the HSF2-Myc (WT/mutants) plasmid after digestion of the inserts by EcoRI and KpnI and cloning into the EcoRI and EcoRV sites in frame with the C-terminal Flag tag in pSNAPf plasmid using In-Fusion Kit (Clontech). The human HSF2-YFP was constructed by PCR and cloned into the XhoI and SalI sites in frame with the N-terminal YFP tag in EGFp-C1 plasmid using In-Fusion Kit (Clontech). All PCR-amplified products for both plasmids were sequenced to exclude the possibility of second site mutagenesis. The cDNA coding for the acetyltransferase domain of murine CBP (1097-1774) was a kind gift of Pr. Ricardo Dalla-Favera (Columbia University, New York) and was used to generate cDNA coding for key domains of CBP: Full HAT (1096-1700), HAT (1322-1700), RING (1205-1279), PHD (1280-1321), Bromodomain (1096-1205), later sub-cloned in pet28a plasmid (Invitrogen) in order to produce 6 His-tagged proteins.

### Patient material

Informed consent for skin biopsy and culture of hPSFs were obtained from the RSTS patients’parents (Patients 1 and 2, [P1] and [P2]; CHU, Robert Debré Hospital, Paris, France; Drs S. Passemard and A. Verloes), in accordance with the Declaration of Helsinki and approved by the local ethics committee, CPP Ile de France, agreement n° P10012 8. Human PSFs from healthy donors have been described in Yehezkel et al., 2008). hPSFs were grown in HAM’s F10 supplemented with 12% FBS in humidified atmosphere with 5 % CO2 at 37°C. See Figure S7A for a description of the deletion or mutation carried by the patients.

### Cell lines

Cell culture, transfections and treatments: murine Neuro2A (N2A, neuroblastoma, DSMZ # ACC 148), Hamster BHK (kindly provided by Dr.Leonhardt H and cultured as described (Herce et *al.,* 2013), human HEK 293T (ATCC®, CRL-11268™), U2OS (osteosarcoma, ATCC®, HTB-96™), U2OS-Crispr*HSF2*KO (2KO), SH-SY5Y (neuroblastoma, ATCC® CRL-2266™) and HeLa-S3 cells (kindly provided by Dr. Slimane Ait-Si-Ali and cultured as described (Yahi *et al*., 2008)) were grown in DMEM (Lonza Group Ltd.) supplemented with 4,5 g/L glucose and 10% fetal bovine serum (FBS, Life technology) in humidified atmosphere with 5 % CO2 at 37°C. RSTS lymphoblastoid cells were obtained from Dr. D Lacombe (CHU Bordeaux; patient 3 (P3)) and from CRB-Institut Médical Jérôme Lejeune, CRB-BioJeL (BB-0033-00016; May 05, 2018, Paris; patients 4 and 5 (P4 and P5)). Lymphoblastoid cells were grown in RPMI (Life technology) supplemented with 4,5 g/L glucose and 10% FBS with L-glutamine (Life technology) in humidified atmosphere with 5 % CO2 at 37°C. See Figure S7A for a description of the deletion or mutation carried by the lymphoblastoid cells. All cell lines were were tested to be mycoplasma free using Venor™ GeM Mycoplasma Detection Kit, PCR-based (Sigma-Aldrich).

### Mice model

Specific pathogen-free C57BL/N female mice were purchased from Janvier (Lyon, France) and maintained in sterile housing in accordance with the guidelines of the Ministère de la Recherche et de la Technologie (Paris, France). Rodent laboratory food and water were provided ad libitum. Experiments were performed in accordance with French and European guidelines for the care and use of laboratory animals. The project has been approved by the Animal Experimentation Ethical Committee Buffon (CEEA-40) and recorded under the following reference by the Ministère de l’Enseignement Supérieur et de la Recherche (#2016040414515579).

### Patients’ primary fibroblasts

For fibroblasts primary cells, The study was conducted in accordance with the Declaration of Helsinki with an approved written consent form for each patient (CPP ESTI: 2014/39; N°ID: 2014-A00968-39), and approval was obtained from the local ethics committee ESTI (license: NCT02873832).

### Generation of CRISPR/Cas9 *Hsf2*KO U2OS cells

Guide RNAs (gRNA) targeting the exon 1 of *HSF2* were designed using our own software (http://crispor.tefor.net/) and cloned into pMLM3636 guide RNA expression plasmid (Addgene 43860). U2OS cells were transfected with Cas9 and guide RNA expression plasmids using Amaxa electroporation as recommended by the manufacturer (Lonza). Cas9 expression plasmid was from the Church lab (Addgene 41815). One week after transfections, cells were seeded at single cell density. Clones were genotyped by DNA sequencing of PCR products spanning the targeted region of the *HSF2* gene. The selected U2OS clone presented 3 different outframe mutations on *HSF2* exon 1, each corresponding to a different allele (Figure S5B). Guide RNA sequence targeting the 1st AUG on *HSF2* exon 1: 5’-UGCGCCGCGUUAACAAUGAA-3’. Primers used for PCR cloning for validation: forward (hHSF2_Cr_ATG_F): 5’-AGTCGGCTCCTGGGATTG-3’ and reverse (hHSF2_Cr_ATG_R): 5’-AGTGAGGAGGCGGTTATTCAG-3’. The genomic sequences of the mutated h*HSF2* alleles and the resulting lack in HSF2 protein and HSF2 DNA-binding activity are described in Figure S5B.

### Production of KIX-GST, His-CBP domain and HSF2 proteins

*Escherichia coli* 21 (DE3) were transformed with the different 6His-tag CBP constructs for production of the different CBP domains as previously described (Duval et al, 2015). All proteins were stored in 20 mM Tris-HCl, 150 mM NaCl, pH 7.5 and kept at −80°C until use. *E. coli* BL21 bacteria were transformed with the different GST-KIX constructs (Cook et al., 2011) and grown in presence of ampicilline and chloramphenicol at 37°C (4-6h). Bacteria were then grown with 1 mM Isopropyl β-D-1-thiogalactopyranoside over-night. After centrifugation (4,400 rpm at 4°C), bacteria were lysed in PBS pH 8, 300 mM NaCl, Triton X100 1%, 1 mG/mL lysozyme, protease inhibitors under stirring at 4°C for 30 min. Bacteria were sonicated (BRANSON sonicator, power 20%, 10’’ ON/20’’OFF) and centrifugated at 4°C, 16000g for 30 min. Gluthatione sepharose 4B beads (G4510-10ML Sigma-Aldrich) were added to the cleared supernatants and subjected to rotation for 1 h 30 min at 4°C. The mixture was loaded into a column (Sigma-Aldrich) and washed with PBS/NaCl 300 mM pH8, then Tris 50 mM/NaCl 150 mM pH 8. Proteins were eluted with 5mL of elution buffer (50 mM Tris HCl pH8, 150 mM NaCl, 10 mM GSH). The protein concentration was measured using Bradford method. *In vitro* transcription and translation reactions were performed using a TNT T7 coupled reticulocyte lysate system as recommended by the supplier (Promega, Charbonnieres-les-Bains, France). 1 µG of the plasmid DNA template was transcribed and the protein was translated at 30°C for 90 min.

### Immunoprecipitation and Western blotting

Protein extracts from cells were prepared using a modified Laemmli buffer (5% sodium dodecyl sulphate, 10% glycerol, 32.9 mM Tris-HCl pH 6.8) supplemented with protease inhibitors (Sigma-Aldrich). Brain tissues were prepared with a lysis buffer (Hepes 10 mM ph 7.9; NaCl 0.4 M, EGTA 0.1 M; glycerol 5%, dithiothreitol (DTT) 1 mM, PMSF 1 mM, protease inhibitor (Sigma-Aldrich), phosphatase inhibitor (Roche)). Then, 30 μg of proteins from lysates were subjected to migration on 8–12 % acrylamide gels and transferred on to polyvinylidene difluoride membranes (GE Healthcare Europe GmbH) in borate buffer (50 mM Tris-HCl and 50 mM borate) for 1 h 45 at constant voltage (48 V). The membranes were incubated with primary antibodies overnight at 4°C, then washed in Tris-buffered saline–Tween 0.1% and incubated for 1 h with horseradish peroxidase (HRP)-coupled secondary antibody (Jackson Immunoresearch). The signal was revealed using a chemiluminescent reagent (Pierce® ECL Plus Western Blotting Substrate, Thermo Scientific) and was detected using hyperfilm (HyperfilmTM ECL, Amersham Biosciences) and a film processor (Konica Minolta). Poly-ubiquitinated HSF2 was detected as described in Ahlskog et al., 2010.

#### For immunoprecipitation of exogenous proteins, using GFP/Myc-Trap

GFP-Trap®-A (ChromoTek) contains a small recombinant fragment of alpaca anti-GFP-antibody, covalently coupled to the surface of agarose beads. It enables purification of any protein of interest fused to GFP, eGFP, YFP, CFP or Venus. HEK 293 cells were transfected by a combination YFP- or Myc-tagged hHSF2 and HA-tagged EP300, CBP (WT or DN) or GFP-tagged HDAC1, or mock vector, with XtremGENE HP Reagent (Sigma-Aldrich) following manufacturer’s instructions. Cells were lysed in Lysis buffer (50 mM Hepes pH 8, 100 mM NaCl, 5 mM EDTA, Triton X-100 0.5 %, Glycerol 10 %, VPA (1 mM), DTT 1 mM, PMSF 1 mM, proteases inhibitors, phosphatase inhibitors (Roche)) and then, HSF2 was immunoprecipitated using anti-GFP- or anti-Myc-trap antibody, or as a control Trap®-A control (ChromoTek). Immunoprecipitated proteins were run on a 8 % SDS-polyacrylamide gel, followed by an immunodetection of CBP or EP300 protein using anti-HA antibody. The amount of immunoprecipitated HSF2 was determined after reblot of the IP membrane with an anti-GFP or anti-Myc antibody. The amount of HSF2 and CBP or EP300 proteins, in the input samples, were detected with anti-GFP or Myc and anti-HA antibodies, respectively.

#### For immunoprecipitation of endogenous proteins

Brain cortices or organoids, or cells (SHSY5Y, N2a) were lysed 30min in Lysis buffer A (25 mM Hepes pH 8, 100 mM NaCl, 5 mM EDTA, Triton X-100 0,5%, 1 mM VPA, 1 mM PMSF, proteases inhibitors, phosphatase inhibitors (Roche)). After centrifugation (15 min, 12 000g) and preclearing, cell lysates were subjected to immunoprecipitation overnight using an anti-mouse HSF2 (Santa-Cruz) and a non relevant IgG (Sigma-aldrich) as a negative control that were pre-incubated 1h at RT with Protein G UltraLink Resin beads (53132, Pierce). Protein complexes were then washed 4 times in wash buffer (25 mM Tris HCl pH7.5, 150 mM NaCl, 1 mM EDTA, Triton X-100 0.1 % Glycerol 10 %, 1 mM VPA, 1 mM PMSF, proteases inhibitors, phosphatase inhibitors (Roche)), and suspended in 2x Laemmli buffer. After boiling, the immunoprecipitates were resolved in 8% SDS-PAGE and immunoblots were performed using an anti-rabbit pan acetyl-Lysine, anti-mouse HSF2 (Santa-Cruz), EP300 (Santa-Cruz) and CBP (CST). The amount of HSF2 and CBP or EP300 proteins in the input samples were detected with anti-mouse HSF2 and anti-rabbit CBP (CST) or EP300 (Santa-Cruz) antibodies.

### Biolayer Interferometry

For *in vitro* protein-protein interaction experiments, we used biolayer interferometry technology (Octet Red, Forté-Bio, USA). Recombinant HSF2 (TP310751 Origen) was desalted (ZebaTM Spin Desalting Columns, 7K molecular-weight cutoff, 0.5 ml (1034–1164, Fisher Scientific, Germany)) and biotinylated at a molar ratio biotin/protein (3:1) for 30 min at room temperature (EZ-Link NHS-PEG4-Biotin (1189-1195, Fisher Scientific, Germany)). Excess Biotin was removed using ZebaTM Spin Desalting Columns. Biotinylated recombinant HSF2 was used as a ligand and immobilized at 100 nM on streptavidin biosensors after dilution in phosphate-buffered saline (PBS; 600s). Interactions with desalted analytes diluted in PBS at 100 nM (recombinant CBP domains 6His-tag Full HAT, Bromodomain (BD), RING), or HSP70 as a positive control (ADI-SPP-555, Enzo-Life Sciences)) were analysed after association (600 s). All sensorgrams were corrected for baseline drift by subtracting a control sensor exposed to running buffer only. For Kd determination, it was calculated with a 1:1 stoichiometry model using a global fit with Rmax unlinked by sensor (FortéBio, Data analysis software version 7.1.0.89).

### Purification of HSF2 from HEK293 cells and mass spectrometry analysis of acetylated lysines

The protocole used is the same as for HSF1, as described by Westerheide et al., 2009. Briefly, HEK 293 cells were transfected with mouse HSF2-beta Flag with or without CMV-EP300, treated with trichostatin A or nicotinamide as indicated 18 h prior harvesting and lysed in RIPA buffer. HSF2-Flag was immunoprecipitated, using α-Flag M2 affinity gel beads (Sigma F2426), and eluted with Flag peptide. Purified mHSF2-Flag was separated by SDS-PAGE, excised from the gel, digested with trypsin, and subjected to tandem mass spectrometric analysis by a hybrid quadrupole time-of-flight instrument (QSTAR, Applied Biosystems, Foster City, CA) equipped with a nanoelectrospray source. MS/MS spectra were searched against the IPI mouse sequence database (68,222 entries; version 3.15) using Mascot (Matrix Science, Boston, MA; version 1.9.05) and X! Tandem (www.thegpm.org; version 2006.04.01.2) database 4 search algorithms. Mascot and X! Tandem were searched with a fragment and precursor ion mass tolerance of 0.3 Da assuming the digestion enzyme trypsin with the possibility of one missed cleavage. Carbamidomethylation of cysteine was included as a fixed modification whereas methionine oxidation, N-terminal protein and lysine acetylation were included as variable modifications in the database search. Peptide identifications were accepted at greater than 95.0% probability as determined by the Peptide Prophet algorithm (7) and validated by manual inspection of the MS/MS spectra, as shown in Table S2. Related to Figure 2C and S2A and Table S1.

### Tandem Affinity Purification (TAP) and M/S identification of HSF2 partners in HeLa cells

We performed retroviral transduction to establish HeLa-S3 cell lines expressing double-tagged HSF2 proteins. *Hsf2-*alpha and *Hsf2-*beta cDNA, cloned from E16 mouse brain were inserted in vector PCEMM-CTAP (Euroscarf P30536; Bürckstümmer et al., 2006) with CMV-driven expression of insert and GFP used as an indirect reporter (IRES; Figure S6A). PCEMM-CTAP allows sequential immunoprecipitation of the tagged protein through two tags: protein G (able to bind IgG) and the streptavidin-binding peptide (GS-TAP; Figure S6A). These two tag-modules are separated by two clivage sites for the Tobacco Etch virus (TEV) protease. Retrovirus production (*Moloney murine leukemia* virus type) and cell transduction with CTAP-empty (no insert), CTAP-Hsf2alpha and CTAP-Hsf2beta were performed as described (Sandrin and Cosset, 2006). GFP-positive Hela-S3 clones were then isolated by clonal dilution, selected by FACS (INFLUX 500, BD BioSciences, IMAGOSEINE Platform, Jacques Monod Institute, Paris) and amplified. By FACS we could isolate four cell sub-populations according to the intensity of the GFP signal to identify a population of cells expressing the recombinant tagged protein at levels similar to that of the endogenous HSF2 protein. GFP-positive Hela-S3 clones stably expressing GS-HSF2alpha, GS-HSF2beta or empty vector, were grown in floating cultures (spinners) and 10 G of cells (3 x 10^9^ cells) per cell line was collected as described (Fristch et al. 2010). Nuclear extracts were prepared as described (Fristch et al. 2010; except the DNase 1 treatment was omitted). Total nuclear extracts were incubated with IgG-agarose beads overnight (Sigma; A2909). Then beads were incubated twice with TEV enzyme (Invitrogen, ref. 12575023)) for 45 min transferred to Poly-prep columns (Bio-rad). Eluates were collected in TEV buffer. These eluates were then incubated with Dynabeads My-one streptavidin for 1 h, washed once, and eluted using 5 mM D-biotin. Proteins were concentrated using TCA/acetone precipitation and dissolved in Laemmli sample buffer. Samples from three independent experiments were sequentially sent to TAPLIN Biological Mass Spectrometry Facility (Harvard Medical School, Boston, MA, USA) for MS (LC/MS/MS) analysis as in Fristch et al. (2010).

### Immunoprecipitation and MS analysis of the endogenous HSF2 protein from E17 mouse cortices

We also analyzed by MS the partners of the endogenous protein HSF2 from E17 fetal cortices. Brain cortices were mechanically lysed into 4.5 volumes of the following buffer: 10 mM Hepes (pH 7.9), 400 mM NaCl, 5 %, glycerol, 100 mM EGTA) and two cycles of freezing and thawing in liquid nitrogen, and centrifuged at 20,000 g for 30 min. Supernatants were used to immunoprecipitate HSF2 using a monoclonal antibody (Abcam) or PBS as a negative control, and protein G agarose (Roche). Immunoprecipitates were analyzed on SDS-PAGE gel bands, containing the HSF2 protein (staining with colloidal blue) were sent to TAPLIN Biological Mass Spectrometry Facility (Harvard Medical School, Boston, MA, USA) for MS (LC/MS/MS) analysis.

### Immunohistochemistry and immunofluorescence

Human organoids sections were prepared as described in Lancaster et al. (2014). Antigen retrieval was performed using citrate buffer (0.1 M Tri-sodium citrate pH 6.0, 10% glycerol, Tween 0,05%) for 1 h at 70°C. Slices were saturated for 30 min with 3% horse serum in PBS -Triton 0,1% and incubated with primary antibody overnight at 4°C. After washing in PBS-Tween 0,1%, slices were incubated with corresponding secondary antibody and DAPI (1 µg/ml) for 1 h at room temperature. Imaging was performed with a confocal microscope Leica TCS SP5 (IMAGOSEINE Imaging Platform in Institut Jacques Monod) and processed with Fiji software.

Cells in basal or heat shock conditions were fixed in 4 % paraformaldehyde on coverslip and stained with HSF1 (CST), HSF2 (Santa-Cruz) or EP300 (Santa-Cruz) antibodies followed by a staining with the corresponding mouse or rabbit fluorescent secondary antibody (Jackson Immunoresearch). Microscopy images were taken on an inverted microscope Leica DMI 6000 and images were analyzed using Fiji software.

### SNAP-Tag labelling of HSF2 molecules and analysis of protein decay

CRISPR/Cas9 *Hsf2*KO U2OS cells were transfected with SNAP-HSF2 WT, -HSF2 3KQ or -HSF2 3KR constructs (Xtrem-Gen HP, Sigma-Aldrich), incubated in the presence of the cell-permeable SNAP-Cell® Oregon green fluorescent substrate (1,25 mM) and then with SNAP-Cell® Block (0,5mM) during the pulse chase according to the manufacturer’s instructions (New England Biolabs). Cells were lysed in modified Laemmli buffer (5% SDS, 10% glycerol, 32.9 mM Tris-HCl pH 6.8) supplemented with 1 mM DTT (Sigma-Aldrich), and their extracts (15 µG) were run on 10% SDS-PAGE. Gels were then scanned on a Typhoon Trio imager (GE Healthcare; excitation 532 nm, emission 580 nm, PMT 700 V) for determination of signal intensity of the covalently-bound fluorescent products as described in (Sanial et al., 2017).

### Fluorescence three-hybrid assay

Fluorescence three-hybrid assay was performed according to Herce *et al*. (2013). BHK cells were transfected with constructs expressing expressing YFP-HSF2, CBP-HA, or EP300-HA, and GBP-LacI, using different combinations (ratio 1:1.5:2) at 70-80 % confluency using reverse transfection by Lipofectamine 2000 (ThermoFisher Scientific), as indicated. Medium was changed after 4 hours for all transfection. After 24 h, the cells were fixed in 4 % paraformaldehyde on coverslip and stained with mouse anti-HA (Covance) or rabbit anti-CBP antibody (Santa-Cruz), followed by a staining with mouse or rabbit fluorescent secondary antibody (Jackson Immunoresearch), respectively. Confocal microscopy images were taken on a confocal microscope Leica TCS SP5 (IMAGOSEINE Imaging Platform in Institut Jacques Monod) and images were analyzed using Fiji software.

### Modeling of the HR-A/B domain and KIX domain of CBP

Prediction of secondary structure of the HSF2 HR-A/B domain was performed using Psipred (http://bioinf.cs.ucl.ac.uk/psipred/) and nps@ (https://npsa-prabi.ibcp.fr/). The tertiary structure of the same domain was predicted using http://petitjeanmichel.free.fr/itoweb.petitjean.freeware.html. Sequence similarity between human HSF2 HR-A/B (Sandqvist et al., 2009), lipoprotein Lpp56 of *E. coli* (Shu et al., 2000), yeast transcriptional factor GCN4 (mutated on some residues to stabilize heptad repeats; Shu et al., 1999) and murine PTRF (Polymerase I and Transcript-Release Factor) and human ATF2 (Activating Transcription Factor 2; a member of the ATF/CREB family) that is known to interact with CBP/EP300 (Bordoli et al., 2001) was explored using *Uniprot* (Pundir et al., 2017; https://www.uniprot.org/) and *ClustalW* (Thompson et al., 1994; https://www.ebi.ac.uk/Tools/msa/clustalw2/). Step 1: Based on this sequence similarity a sequence alignment of H-RA/B was developed (using *Uniprot and ClustalW*; see Figure S4E), a structural model of the monomer HRA/B was generated using using *Modeller* (v9.19 (Eswar et al., 2006; https://salilab.org/modeller/), verified by *ERRAT* (Colovos et al., 1993; http://servicesn.mbi.ucla.edu/ERRAT/) and *RESprox* (Berjanskii et al., 2012; http://www.resprox.ca/) and Ramachandran plot (see Figure S4F Ramachandran et al., 1963) followed by the development of the trimer using *SymmDock* (Schneidman-Duhovny *et al*., 2005; v Beta 1.0; http://bioinfo3d.cs.tau.ac.il/SymmDock/). Step 2: the interaction between HR-A/B and the KIX domain (pdb: 2LXT; Brüschweiler et al., 2013) was simulated and compared using *Zdock* (v 2.1; Pierce et *al.*, 2014) and *Firedock* (Mashiach et al., 2008; Odoux et al., 2016). More precisely, the ten best results generated by *Zdock* and *Firedock*, and scored according to their Root Mean Square Deviation (RMSD), which were compared thanks to a visualization program ICM (Fernandez-Recio et al, 2005). The mutation of the key residuse involved in the interaction between HR-A/B and the KIX domain have been performed using PyMOL (v2.0) and the docking was done as described above.

### RP-UFLC-based separation and quantification of CBP substrate peptides (HSF2) and their acetylated forms

For acetylation assays, we synthetized several 5-fluorescein amidite (5-FAM)-conjugated peptide substrates based on the human HSF2 sequence and containing various lysine residues of interest (Proteogenix):

- 5-FAM-SGIVK82QERD-NH2, referred to as K82 peptide
- 5-FAM-SSAQ135VQIR-NH2, referred to as K135 peptide
- 5-FAM-SLRRK197RPLL-NH2, referred to as K197 peptide

We also synthesized acetylated versions of these HSF2 peptides as standards. Samples containing HSF2 peptides and their acetylated forms were separated by RP-UFLC (Shimadzu) using Shim-pack XR-ODS column 2.0 x 100 mm 12 nm pores at 40°C. The mobile phase used for the separation consisted of 2 solvents: A was water with 0.12 % trifluoacetic acid (TFA) and B was acetonitrile with % TFA. Separation was performed by an isocratic flow depending on the peptide:

- 80% A/20 % B, rate of 1 ml/min for K82 and K135
- 77% A/23 % B, rate of 1 ml/min for K197

HSF2 peptide (substrate) and their acetylated forms (products) were monitored by fluorescence emission (λ= 530 nm) after excitation at λ= 485 nm and quantified by integration of the peak absorbance area, employing a calibration curve established with various known concentrations of peptides.

### *In vitro* acetyltransferase assay

To determine the activity of recombinant CBP-Full HAT on HSF2 peptides, we used 96-wells ELISA plate (Thermofisher) and assays were performed in a total volume of 50 µL of acetyltransferase buffer (50 mM Tris pH8, 50 mM NaCl) with 500 nM CBP-Full HAT, 50 µM HSF2 peptides and 1 mM DTT. Reaction was then started with the addition of 100 µM Acetyl-CoA (AcCoA) and the mixture was incubated 20 min at room temperature. 50 µL of HClO4 (15 % in water, v/v) was used to stop the reaction and 10 µL of the mixture were injected into the RP-UFLC column for analysis. For time course studies, aliquots of the mother solution were collected at different time points and quenched with 50 µL of HClO4 prior to RP-UFLC analysis.

### Statistics

Data are displayed as means ± standard deviation (SD). GraphPad Prism 8 (GraphPad Software, La Jolla, CA, USA) was used for statistical analyses. Statistical significance was assessed using Wilcoxon matched-pairs signed rank test for two groups with paired values (Figure 5–6,S6) or the Mann-Whitney test for two groups (Figure 7,S7). *p*–values below 0.05 are considered statistically significant.

## Acknowledgments

We warmly thank the patients and their families for their participation in this study. We thank Slimane AIT-SI-ALI for helpful discussions and comments on the manuscript, Anne PLESSIS (Jacques Monod Institute, Paris, France) for helpful discussions on setting the SNAP-TAG technology, Anne VANET (Jacques Monod Institute, Paris, France) for helpful discussions on the HSF2 structural modelling. We thank Lauriane FRITSCH and Slimane AIT-SI-ALI (UMR7216, for HeLa-S3 cells and growth conditions for TAP-TAG analyses). We thank the Imaging Platform IMAGOSEINE and especially Nicole BOGETTO for her help in sorting the GFP-positive HeLa-S3 cells. We are grateful to Heinrich LEONHARDT (Ludwig-Maximilians University, Munich, Germany) for F3H cellular and molecular tools and Pierre-Antoine DEFOSSEZ and Laure FERRY (UMR7216) for helpful guidance in F3H and GFP-Trap experiments, Isabelle LEMASSON (East Carolina University, USA) for the KIX-GST constructs, and Sophie POLO (UMR7216) for SNAP-TAG vectors. We are grateful to Vincent El Ghouzzi for the kind gift of human induced pluripotent stem cells (hIPSCs). We thank Isabelle COUPRY and Benoit ARVEILER (CHU de Bordeaux, France) for primary skin fibroblasts from healthy donors. We thank the Institut Médical Jérôme Lejeune for the gift of Lymphoblastoid cells (patients 4 and 5). We are grateful to Delphine BOHL, and Stéphane BLANCHARD from Pasteur Institute (Rétrovirus et Transfert Génétique, INSERM U622) for their help in producing the retroviruses for TAP-TAG experiments in Hela-S3 cells. We thank Laure FERRY (UMR7216) and the Epigenomics Platform, as well as Sandra PIQUET (UMR7216) and the Microscopy Platform (UMR7216) for access to instruments and technical advice, and Clara GIANFERMI (UMR7216) for microscopy pictures of organoids and nSBs. We thank Isabelle Le PARCO and the staff from the Buffon animal housing facility at the Jacques Monod Institute (Paris Diderot University, Paris, France) and the *Bioprofiler* Platform at the UMR8251 Biologie Fonctionnelle et Adaptative for *in vitro* acetylation assays.

## Funding information

VM was funded by the CNRS (Projet International de Coopération Scientifique PICS 2013-2015) for her collaboration with LS and by the Short Researcher Mobility France Embassy/MESRI-Finnish Society of Sciences and Letters; the Agence Nationale de la Recherche («NeuroHSF», Programme Neurosciences, Neurologie and Psychiatrie ANR-06-NEURO-024-01 and « HSF-EPISAME », SAMENTA ANR-13-SAMA-0008-01). LS was funded by the Academy of Finland, Sigrid Jusélius Foundation, Magnus Ehrnrooth Foundation and Cancer Foundation Finland. RA was supported by PhD Fellowships from Neuropôle Ile-de France and Fondation ARC, FM by PhD Fellowships from the CNRS and Fondation pour la Recherche Médicale and from a Postdoc Fellowship from SAMENTA ANR-13-SAMA-0008-01), AD by a Ministère de l’Enseignement supérieur, de la Recherche et de l’Innovation (MESRI) Doctoral Fellowship, and ADT, DSD, and DB by the PICS travel grant, and GP and MH by Master 2 Intership Fellowships from SAMENTA ANR-13-SAMA-0008-01. JB was supported by a PhD Fellowship from Région Ile-de-France (Cancéropôle IDF) and University Paris Diderot. JKA was supported by Magnus Ehrnrooth foundation. MCP was supported by the Turku Doctoral Network in Molecular Biosciences and Magnus Ehrnrooth Foundation. The supporting bodies played no role in any aspect of study design, analysis, interpretation or decision to publish this data.

## LIST OF SUPPLEMENTARY FIGURES

Supplemental Information includes 7 figures. Figure S1 to S7 are relative to Figure 1 to 7.

## LIST OF SUPPLEMENTAL TABLES

**Table S1.** Summary of the HSF2 acetylated peptides identified by MS

**Table S2.** Original tables and spectra corresponding to the HSF2 acetylated peptides identified by MS

**Table S3. Optimal docking area (ODA)/ tableau des scores de docking/mutation inefficace sur HSF2.** Related to Experimental procedures.

