## Supplemental data for "CBP/EP300 acetylates and stabilizes the stress-responsive Heat Shock Factor 2, a process compromised in Rubinstein-Taybi syndrome"

**SUPPLEMENTARY DATA**

**LEGENDS OF SUPPLEMENTAL FIGURES**

**Figure S1.** (related to Figure 1)

**(A) *HSF2, EP300, and CBP (CREBBP) mRNAs are expressed along the differentiation process of human brain organoids*.** Graph corresponding to the RNA-Seq data from two independent sets of D20 embryoid bodies (EB_D20_A and _B), and organoids at D40 (ORG_D40_A and _B) and D60 (ORG_D60_A and _B). The amount of mRNAs is expressed as RPKM (reads/kb/million mapped reads).

**(B) *HSF2 is expressed in areas containing NPCs and neurons.*** Upper and lower panels (a and f) whole view of a section of two different human organoids. The white rectangle indicates the magnified areas in (b-e) and (g to j) in the two panels. (b,g) phase contrasts. (c,h) DAPI staining.

(Upper panels) HSF2 is expressed (d; red) in the DAPI-dense area (c) and in the area of Tuj1-positive neurons (e; green). Scale bar (a): 500 µm. As indicated by thin rectangles in (c, d, and e), (c1, d1, and e1) and (c2, d2, and e2) the magnified areas in the zones of low DAPI density-high Tuj1 signal and high DAPI density-low Tuj1 signal, correspond to neurons (arrows) and NPCs (arrowhead), respectively, as shown in the lower panels.

(Lower panels) DAPI-dense areas (h) correspond to Sox2-positive (i; red), which are also Tuj1-negative cells (j), that is, to NPCs. Scale bars: upper panels (a) 500 µm, (b-e) 50 µm; lower panels, (a,f) 300 µm, (b-e, g-j) 50 µm.

**(C)** ***CBP is expressed in areas containing NPCs and neurons***. CBP staining corresponds to cells of dense DAPI staining (NPCs) and also some Tuj1 positive cells (green; neurons). Scale bar: 50 µm.

**(D)** ***HSF2, CBP, EP300 are expressed at all stages of mouse cortical development***. WB analysis of forebrain extracts from E11 to E17 of gestation. The expression profile of HSF2 is in line with our previous data (Rallu et al., 1997; Kallio et al., 2002; Chang et al., 2006; El Fatimy et al. 2014) and that of CBP and EP300 are also in line with their reported patterns (Kawasaki et al. 1998, Yao et al., 1998 Partanen et al., 1999, and Bhattacherjee et al; 2009). Tel: telencephalon.

**(E) *HSF2 interacts with EP300 and is present in an acetylated form in the E10 mouse cortex.*** (Left upper panels) Co-immunoprecipitation of endogenous EP300 and CBP proteins, using anti-HSF2 antibody. (Right upper panels) inputs. (Lower panels) Acetylation of the immunoprecipitated HSF2 protein from E10 cortical extracts. The blot was incubated with anti-pan-AcK antibody (WB:AcK; left) and then re-incubated with anti-HSF2 antibody (WB:HSF2; right)). Brackets: HSF2 forms of higher molecular weights are typically detected in the mouse cortex (El Fatimy et al., 2014), possibly corresponding to additional post-translational modifications, like sumoylation (Anckar et al. 2011).

**(F)** **Endogenous HSF2 is acetylated in the human neural precursor cell line SHSY-5Y**. Endogenous HSF2 is immunoprecipitated from SHSY-5Y cells pre-treated with MG132 (20 µM for 6 h) to increase HSF2 protein levels and optimize its detection (Mathew et al., 2008; Ahlskog et al., 2010). Its acetylation status was explored using anti-Pan-AcK antibody, as in (**E**).

**Figure S2. HSF2 is acetylated by CBP and EP300 in normal conditions.** (related to Figure 2).

**(A) *Positioning of the acetylated peptides identified by MS in the mHSF2β protein***. Each peptide is represented as a line and numbered in red. The color code corresponds to the different HSF2 domains, as in Figure 2C. The peptides and their sequences are also listed in Table S1.

**(B) *Comparative impact of the mutations of single residues*** ***(K82, K128, K135, and K197), and of the doublet K209/K210 on HSF2 global acetylation by EP300***. HEK 293 cells were co-transfected with EP300-HA and HSF2-Myc WT or a mutated HSF2 on the indicated lysine residues. HSF2 was immunoprecipitated using an anti-Myc antibody and its acetylation was detected using a pan-acetyl-lysine (AcK) antibody. HSC70, loading control.

**(C) *Impact of the combined mutations of 3 lysines (K128, K135, and K197)*** ***or 4 lysines* K82, K218, K135, and K197)** ***into glutamines (3KQ or 4KQ) or arginines (3KR or 4KR) on the acetylation levels of HSF2*.** HEK 293 cells were co-transfected with CBP-HA and the different HSF2-Myc constructs (wild-type (WT) or indicated mutants). Actin, loading control.

**(D) *Kinetics of* in vitro *acetylation of HSF2 peptides containing the K82 residue*.** RP–UFLC experiments. As in Figure 2E. Note that we could not perform this experiment on the HSF2 peptide containing K128, due to its insolubility.

**(E) *Control experiments for the determination of the elution profiles*** of the K82 and acetylated K82 (AcK82), K135 and AcK135, and K197 and AcK197 peptides, using non-acetylated and synthetically acetylated commercial peptides. Separation was monitored by RP-UFLC.

**(F)** ***Control experiments for Acetyl-CoA-dependent acetylation of HSF2 K82, K135 and K197 peptide in the presence of CBP-Full HAT.*** HSF2 peptide substrates were incubated in the presence of recombinant purified CBP-Full HAT and with or without acetyl-CoA for 20 minutes. Acetylated products were then analysed by RP-UFLC.

**Figure S3. HSF2 interacts with CBP and EP300 in normal conditions.** (related to Figure 3).

**(A) *Determination of the Kd of CBP Full-HAT domain affinity to HSF2.*** Biolayer interferometry to determine association and dissociation curves of CBP Full-HAT domain (concentration range from 12.5 μM to 400 nM) with biotinylated HSF2 immobilized on streptavidin sensor tips (blue curves). As a positive control the binding profile of HSP70 (100 nM) is plotted (red curve).

**(B) *Interaction between endogenous HSF2 and CBP/EP300 proteins in neural cells.***

N2A cells were treated or not (-) for 3 hours with valproic acid, a Class I HDAC inhibitor (VPA, 1 mM). HSF2 and co-precipitated CBP (upper panels) or EP300 proteins (lower panels) were detected by WB. Inputs: Total amounts of proteins in the extracts. Actin, loading control. *: IgG heavy chain. Representative immunoblots (n=3 experiments).

**(C)** ***F3H control experiments for the visualization of interaction between HSF2-YFP and exogenous CBP-HA***. As in Figure 3F (upper panels). Scale bar: 10 µm.

**(D)** ***F3H control experiments for the visualization of interaction between HSF2-YFP and endogenous CBP***. As in Figure 3F (lower panels). Scale bar: 10 µm.

**(E) *F3H control experiments for the visualization of interaction between HSF2-YFP and exogenous EP300-HA*.** As in Figure 3G. Scale bar: 10 µm.

**Figure S4. Identification of the HSF2 domains that interact with CBP** (related to Figure 4).

**(A)** ***Qualitative summary of the impact of the deletion of different functional domains of Flag-HSF2 on its acetylation status and interaction with CBP-HA***, corresponding to experimental data shown in (B-D). (+++) strong; (+) moderate; (+/-) low acetylation or interaction; (ND) non detectable: in that case, the Flag tag is not recognized by the antibody, likely because it is masked by the aberrant conformation of the truncated HSF2 protein; (-*) not observed (the interaction might be very labile in the absence of the TAD domain, a typical docking site for CBP in many transcription factors).

**(B and C) *Determination of the HSF2 domains necessary for HSF2 acetylation by CBP-HA***

HEK 293 cells were transfected with CBP-HA and the WT HSF2-Flag or its different deleted forms.

**(B)** (Upper panels) Immunoprecipitated WT or deleted HSF2-Flag forms, described in (**A**), were checked for their acetylation status, as determined by WB using an anti-pan-acetyl-lysine antibody (WB: AcK) and the membrane was re-incubated with anti-Flag antibody (WB: Flag) to verify the equivalent expression levels of the different HSF2 constructs. (Middle panels) Inputs, total amounts of proteins in the extracts. (Lower panels) WT HSF2-Flag is not acetylated in the absence of CBP-HA (negative control). Representative immunoblots.

**(C)** Representative immunoblot of HSF2 acetylation by CBP-HA comparing WT Flag-HSF2 with the HSF2 deleted forms described in (**A**). (upper panel) CBP-HA expression. The HSF2 acetylation status was determined by WB using an anti-pan-acetyl-lysine antibody (WB: AcK) as in (**B**). Asterisks point to the acetylated forms of the WT and deleted HSF2-Flag. HSC70, loading control.

(n=3 independent experiments).

**(D) *Determination of HSF2 domains involved in the interaction with CBP.***

Co-immunoprecipitation of CBP-HA with WT or deleted HSF2-Flag was checked using anti-HA antibody (upper panel) and anti-Flag (lower panel) antibody to verify the comparable expression level of HSF2 constructs (as in **B**).

**(E)** ***A sequence alignment of HR-A/B*** (from aa. 121 to 201) between human HSF2 HR-A/B, lipoprotein Lpp56 of *E. coli*, yeast transcription factor GCN4 (mutated on some residues to generate stabilized heptad repeats) and murine PTRF and human ATF2 transcription factors (Shu *et al.,* 1999, 2000; Sandqvist *et al.,* 2009). The alignment was developed, using *Discovery Studio* and *Clustal W multiple sequence alignment program*. Sign code: ":" indicates that one of the 'strong' groups of residues is fully conserved; "*" marks the positions which have a single, fully conserved residue; "." indicates that one of the 'weaker' groups is fully conserved.

**(F)** ***Ramachandran plot*** (*Discovery studio*) showing the good quality of the triple coiled-coil model structure of the HR-A/B determined based on sequence similarity of the proteins cited in (**E**). Relative to Figure 4B.

**(G) In silico** ***analysis of the single*** ***mutations*** ***Y650A in the CBP KIX domain, F181, V183, K177, and Q180A in the HSF2 KIX recognition motif***. The mutations Y650A (*Zdock* analysis), K177A and Q180A hamper the interaction between the HRA/B domain and KIX domain, while F181A and V183A have no effect (*Firedock* analysis). Relative to Figure 4D and E.

**Figure S5.** (related to Figure 5).

**(A) *Decreased acetylation levels of lysine residue K18 of histone H3 (H3K18Ac) assess C646 efficiency in inhibiting CBP/EP300 activity***. HSC70, loading control. Relative to Figure 5A.

**(B)** ***Sequence of the HSF2 alleles mutated in U2OS cells using the CRISPR/Cas9 KO strategy and corresponding protein products***.

**(C) *Representative immunoblot analysis of HSF2 content in the CRISPR/Cas9 Hsf2KO cells***. U2OS parental and SHSY-5Y cell extracts were loaded as positive controls for the detection of HSF2. Actin, loading control.

**(D)** ***Myc-HSF2WT, HSF2 3KQ, HSF2 3KR, and HSF2 4KR are ectopically expressed at similar levels in 2KO cells***. Representative immunoblot analysis of transient transfection experiments with the corresponding constructs. Actin, loading control.

**(E)** ***Ectopically expressed*** ***Myc-HSF2WT, and mutant HSF2 3KQ, HSF2 3KR, and HSF2 4KR* *exhibit HSF2 DNA-binding activity* ex vivo**. Gel-shift analysis of HSF1 and HSF2 DNA-binding activity in *2*KO cells, expressing Myc-tagged HSF2WT, HSF2 3KQ, HSF2 3KR, or HSF2 4KR, (or Myc (mock) as a negative control). The presence of HSF1 and/or HSF2 in the HSF-HSE complex was analyzed by supershifting (arrowhead) with anti-HSF1 (α1; black arrowhead) or anti-HSF2 (α2; white arrowhead), or anti-Myc antibodies (α-Myc; blue arrowhead). N2A cells were loaded as positive controls for anti-HSF2 antibodies. Notably, *2*KO cells expressing HSF2WT, HSF2 3KQ, HSF2 3KR, and HSF2 4KR proteins allow the formation of a HSE-HSF complex, which is mainly supershifted by anti-HSF2 (but almost not by anti-HSF1), whereas, as expected, mocked transfected *2*KO cells are devoid of HSF2 activity. HSF-HSE: HSF-HSE complexes. CHBA: constitutive HSE-binding activity, which is not carried by HSFs (Mosser et al., 1988; Abravaya et al., 1991); NS: non-specific DNA-protein complex; free: unbound double-stranded HSE oligonucleotide.

**(F)** ***Myc-HSF2WT, HSF2 3KQ, or HSF2 3KR proteins localize into nuclear-stress bodies upon HS.*** Immunofluorescence analysis of the ability of the WT or mutated HSF2 to localize to nSBs when ectopically expressed in U2OS cells, in response to a 1h HS at 42°C, using anti-Myc antibodies. Scale bar: 20 µm.

**Figure S6. Impact of HDAC1 on HSF2 levels in normal and stress conditions.** (related to Figure 6).

**(A) *Schematic representation of the two-step TAP-TAG approach for identification of HSF2 partners in the nucleus of HeLa-S3 cells*** (Bürckstümmer et al., 2006) transfected with PCEMM-CTAP-HSF2α or PCEMM-CTAP-HSF2β, using the G-protein (in yellow) and streptavidin binding peptide (SBP in red), as dual affinity tags (GS-TAP), sequentially. Relative to Figure 6A.

**(B)** ***Analysis of whole cell (Tot), cytosolic (Cyto), and nuclear (Nuc) extracts*** from the HeLa-S3-CTAP-empty, HeLa-S3-CTAP-HSF2α and HeLa-S3-CTAP-HSF2β cell populations by WB. Enrichment in histones in nuclear fractions by Coomassie blue was verified in a first step (data not shown). Relative to Figure 6A.

**(C) *Silver staining of SDS-PAGE analyses of final double-TAP-TAG eluates*** from HeLa-S3-CTAP-empty, HeLa-S3-CTAP-HSF2α and HeLa-S3-CTAP-HSF2β nuclear extracts. Relative to Figure 6A.

**(D)** ***Identification of HDAC1 and 2 as HSF2 protein partners by MS in the E17 mouse brain cortex***. (Upper panel) Colloidal blue staining of the gel, containing HSF2 immunoprecipitates from the E17 fetal telencephalon, prior to MS analysis. The rectangles delimit the gel bands that were cut and subjected to MS. (Lower panel) number of unique peptides from each identified protein and their UniProt Knowledgebase (UniProtKB) codes are indicated.

**(E) *F3H control experiments for*** ***the visualization of the interaction between HSF2-Flag and exogenous HDAC1-GFP***. As in Figure 6B. Scale bar: 10 µm.

**(F) *HDAC8 does not affect HSF2 acetylation levels, in contrast to HDAC1.*** n=3 independent experiments. Relative to Figure 6D.

**(G) *Endogenous HSF2 levels are decreased upon HS in N2A cells***. Representative immunoblot and quantification of n= 3 experiments. n=3 independent experiments.

**(H) *Assessment experiment for the efficiency of the HDAC inhibitor, VPA, in N2A cells***. Representative immunoblot for H3K27 acetylation (AcH3K27). (n=3 independent experiments). Relative to Figure 6F.

**(I) *Expression of dominant-negative HDAC1 prevents the accumulation of poly-ubiquinated HSF2 upon HS***. HEK 293 cells were cotransfected with HSF2-Myc, EP300-HA, and Flag-HDAC1 or dnHDAC1, and subjected or not to HS 42°C (30 min). Representative immunoblot of polyubiquinated HSF2 levels after Myc-HSF2 immunoprecipitation. n=3 independent experiments.

**Figure S7. Altered HSF2 protein levels and dysregulation of the stress response in RSTS.** (related to Figure 7).

**(A) *Description of the mutations or deletions present in RSTS patients*.** The scheme of the genomic organization of the genes are taken from [https ://www.ncbi.nlm.nih.gov/nuccore/NM_001429.3](https://www.ncbi.nlm.nih.gov/nuccore/NM_001429.3)

**RSTS hPSFs were isolated from two patients (P1 and P2):**

**Patient P1- hPSF RSTS*_CBP_***: Mutation pLys1139X (exon 18) located in the catalytic CBP HAT domain, needed for its acetyltransferase activity. This mutation did not affect CBP or EP300 levels and a marked impact on H3K27 acetylation was observed (Figure S7B).

**Patient P2- hPSF RSTS*_EP300_***: Deletion of at least 10 kb (chr22:41533381.41543435) and to the maximum of 11 kb (chr22:41532396.41543758) confirmed by quantitative multiplex fluorescent (QMF)-PCR. This mutation leads to a truncated protein, with deletion of a binding site, the KIX domain, which is essential for the interaction between HSF2 and EP300/CBP proteins. We show that hPSF exhibit a decreased level in H3K27 acetylation compared to HD as expected in this cell model (Jin, PMID: 21131905; Figure S7B).

**Lymphoblastoid cells were derived from three patients (P3-P5):**

**Patient P3- LB RSTS*_EP300_***: Mutation pTrp1649X (Exon 30) in *EP300 gene* which gives rise to a stop codon in the HAT domain leading to a C-terminal deletion. This mutation leads to >50% decrease of EP300 protein level with a predicted altered activity (MutPred2).

**Patient P4- LB RSTS*_CBP_***: Deletion in 5’ (2 first exons) in *CREBBP gene*. This mutation leads to a truncated protein and decrease in CBP protein levels.

**Patient P5- LB RSTS*_CBP_***: Mutation p.Arg1498* (exon 27) located in the catalytic CBP HAT domain with formation of a stop codon leading to a C-terminal deletion. This mutation leads to a decrease of CBP protein level.

**(B) *Reduction of H3K27 acetylation (AcH3K27) in hPSFs from RSTS patients compared to HD***. (Upper panels) RSTS*_CBP_* [P1]. (Lower panels) RSTS*_EP300_* [P2].

**(C) *VPA does not increase HSF2 levels in RSTS_CBP_ [P1] hPSFs, as in RSTS_EP300_ [P2]*** hPSFs Relative to Figure 7A and B.

**(D)** ***Assessment of the efficiency of VPA in RSTS_CBP_ hPSFs (patient P1)***, in increasing in H3K27 acetylation. Related to Figure 7B.

**(E) *HD and RSTS_EP300_ hPSFs (patient P2)*** ***contain similar levels of HDAC1*** by immunofluorescence detection (left) and WB (right). Scale Bar, 10 µM.

**(F) *Formation of nSBs, with HSF1 and EP300 expression in HD and RSTS_EP300_ hPSFs, upon HS*.** Representative immunofluorescence experiment, in control (CTR) or HS conditions (1 hour at 42°C). Scale bar: 20 µm. Relative to Figure 7D and E. Scale Bar, 10 µM.

**(G)** ***Altered formation of nSBs by HS in*** ***RSTS*_EP300_ or *RSTS*_CBP_ *LBs.*** (Left panel) Representative pictures of cells in HS conditions (1 hour at 43°C). (Right panel) Quantification of the percentage of cells positive for nSBs, from 100 – 150 cells in n = 3 different experiments. RSTS LBs were compared to HD LBs. Standard deviation. *, p < 0.05. Scale Bar, 10 µM.
