## Supplemental Figures for "CBP/EP300 acetylates and stabilizes the stress-responsive Heat Shock Factor 2, a process compromised in Rubinstein-Taybi syndrome"

### Suppl. Figure S1

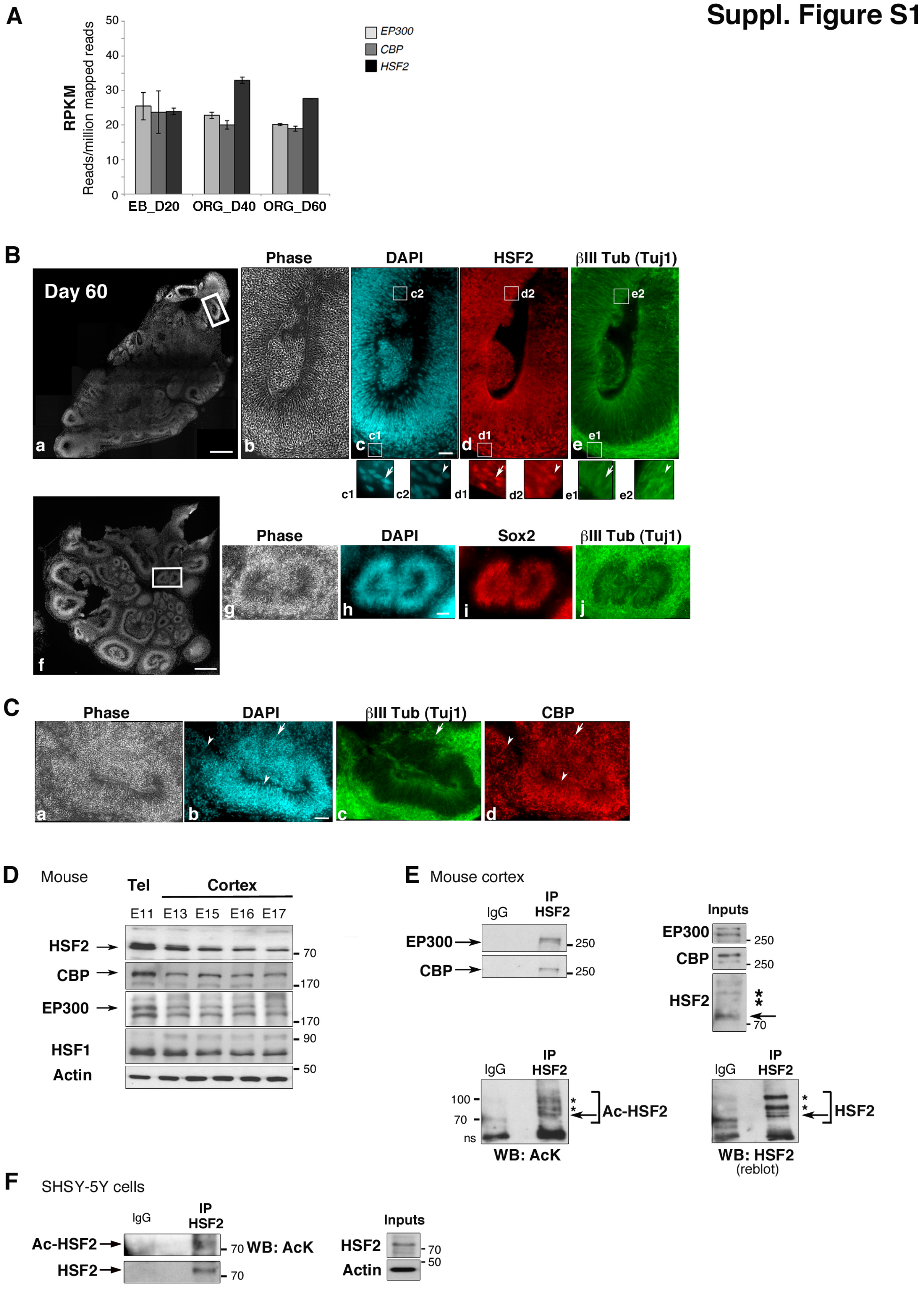

### Suppl. Figure S2

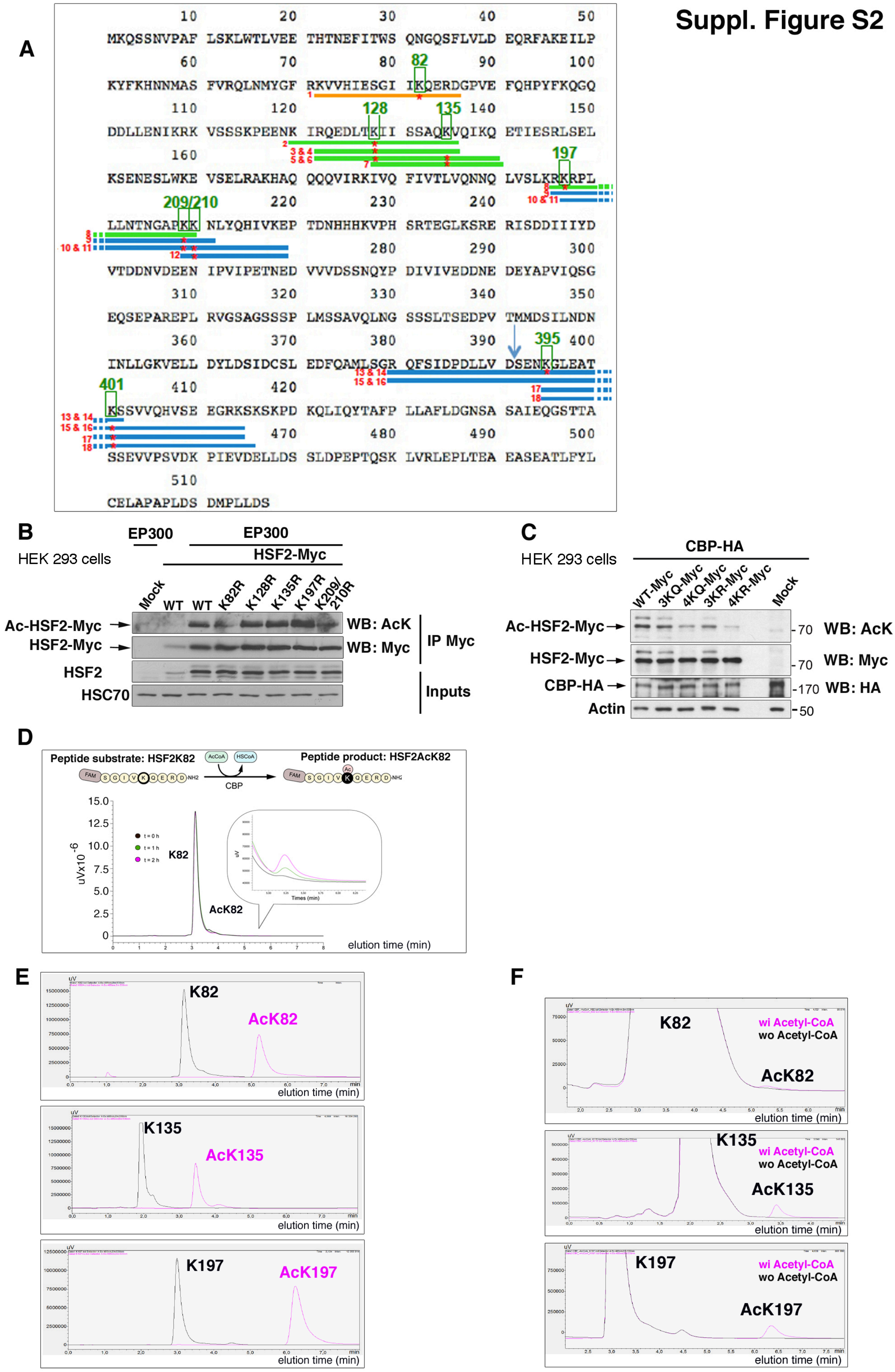

### Suppl. Figure S3

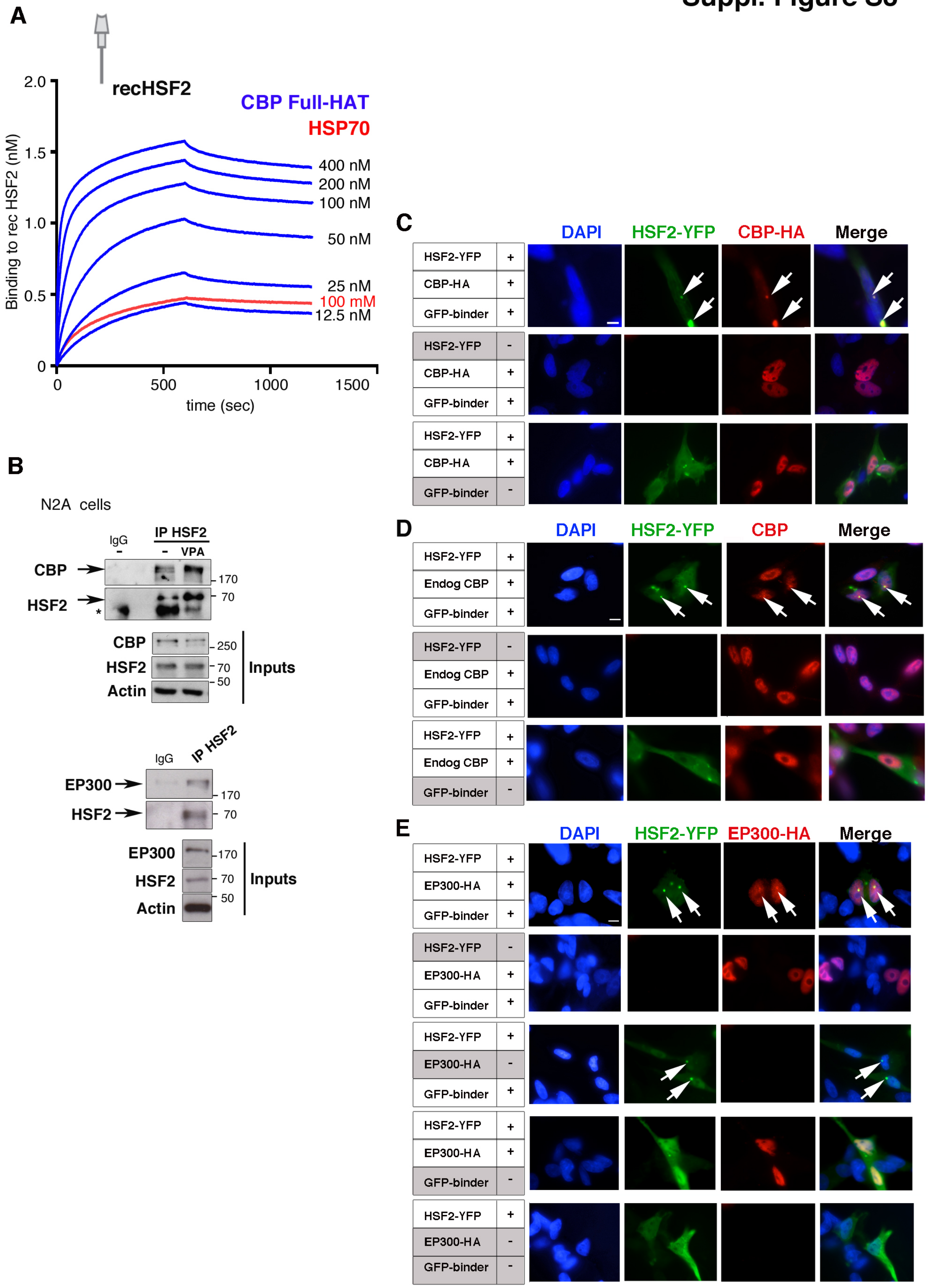

### Suppl. Figure S4

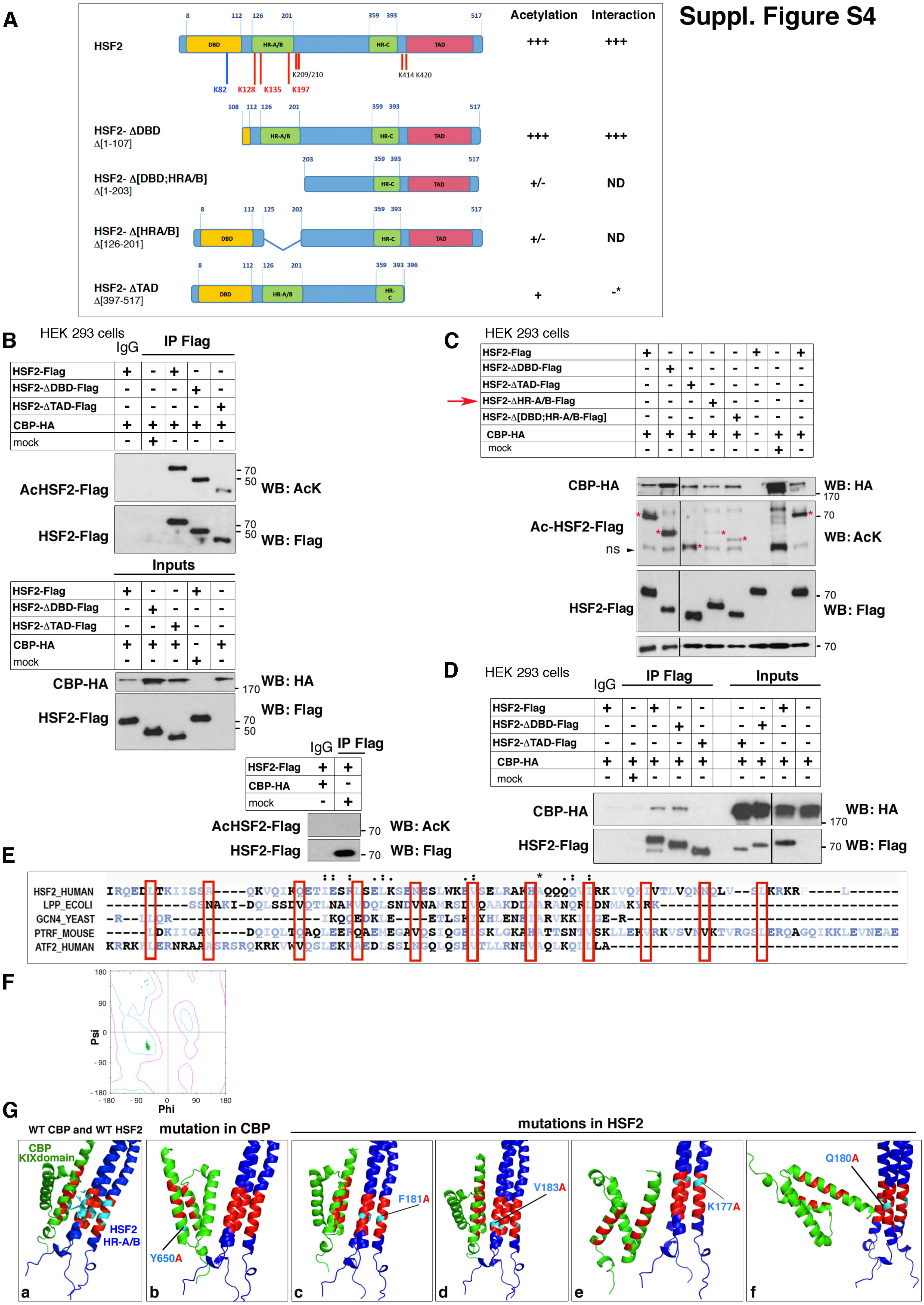

### Suppl. Figure S5

**A**

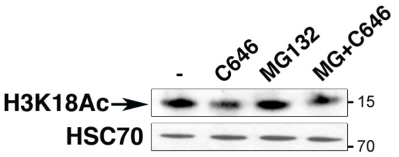


**B**

| | Genomic sequence | mutation | Protein product |
|---|---|---|---|
| WT | TGCGCCGCGTTAACAATGAAGCAGAGTTCGAACGTGCCGGCTTTCCTCAG CAAGCTGTGGACGCTTGTGGAGGAAACCCACACTAACGAGTTCATCACCT | | HSF WT protein |
| Allele 1 | TGCGCCGCGTTAACAATGAAG~~C~~AGAGTTCGAACGTGCCGGCTTTCCTCAG CAAGCTGTGGACGCTTGTGGAGGAAACCCACACTAACGAGTTCATCACCT | 1-nt deletion | 61-aa truncated protein (only the first six aa correspond to HSF2) |
| Allele 2 | TGCGCCGCGTTAACAATGAAGCAGAGTTCGAAC~~G~~TGCCGGCTT**C**CCTCAG CAAGCTGTGGACGCTTGTGGAGGAAACCCACACTAACGAGTTCATCACCT | 1nt deletion + 1 substitution | 61-aa truncated protein (only the first six aa correspond to HSF2) |
| Allele 3 | TGCGCCGCGTTAACAATGAAGCAGAGTTCGAACG~~TGCCGGCTTTCCT~~ ~~CAGCAAG~~CTGTGGACGCTTGTGGAGGAAACCCACACTAACGAGTTCATC ACCT | 20nt deletion | 16-aa truncated protein (only the first six aa correspond to HSF2) |

**C**

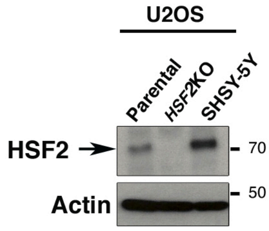


**D**

U2OS *HSF2*KO cells

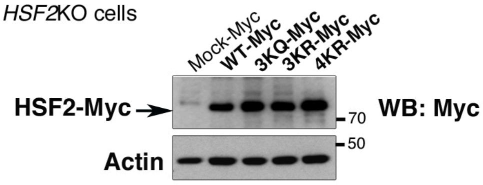


**E**

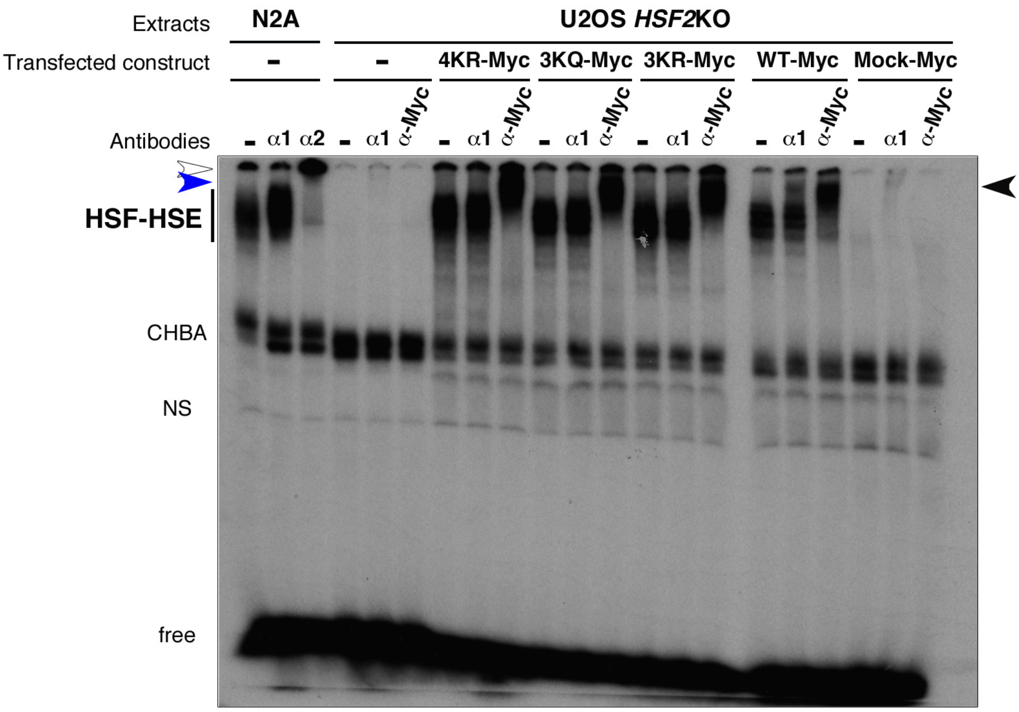


**F** U2OS cells - HS (42°C)

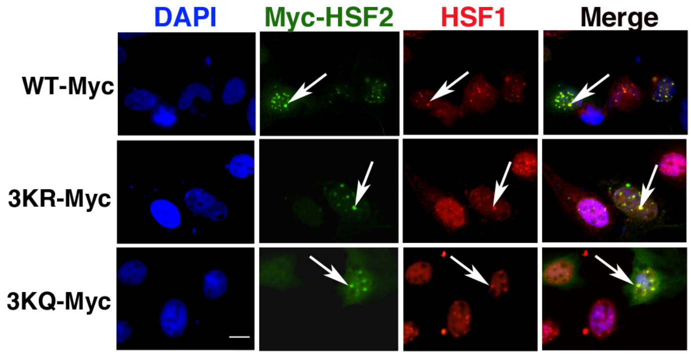

### Suppl. Figure S6

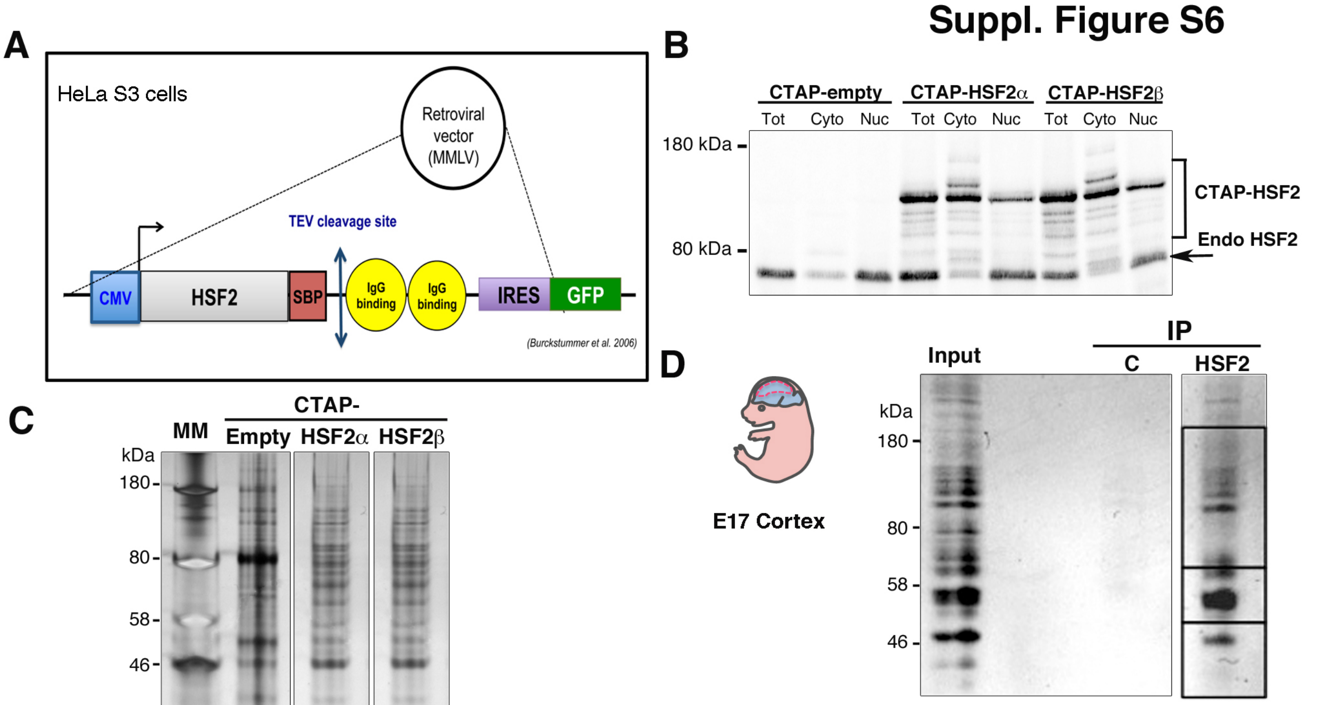


| Sample | Number of peptides | Protein match | UniprotKB code |
|---|---|---|---|
| IP HSF2 from mouse Cortex | 3 | HDAC2_MOUSE | Q3TMT1 |
| | 2 | HDAC1_MOUSE | O09106 |

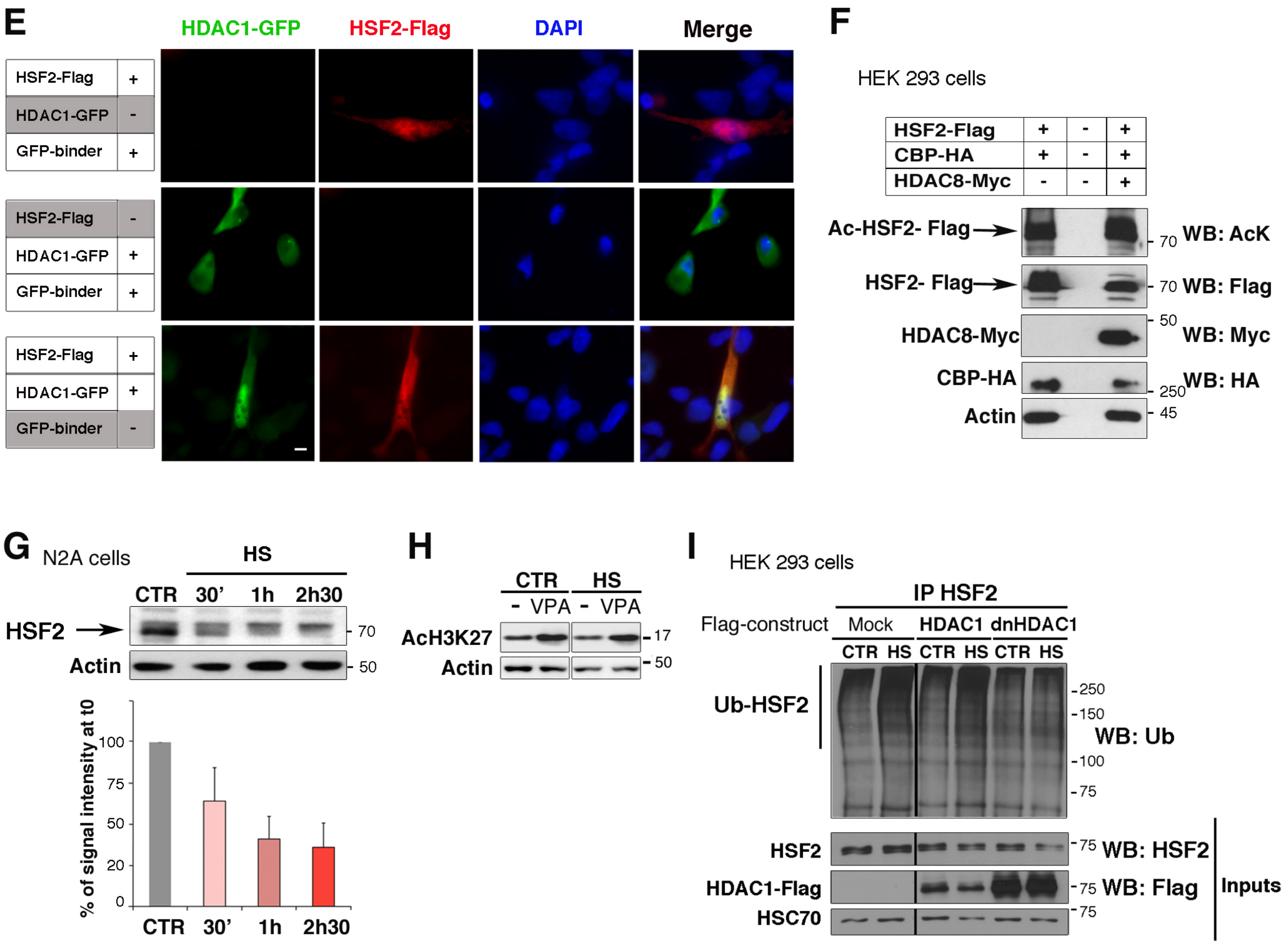

### Suppl. Figure S7

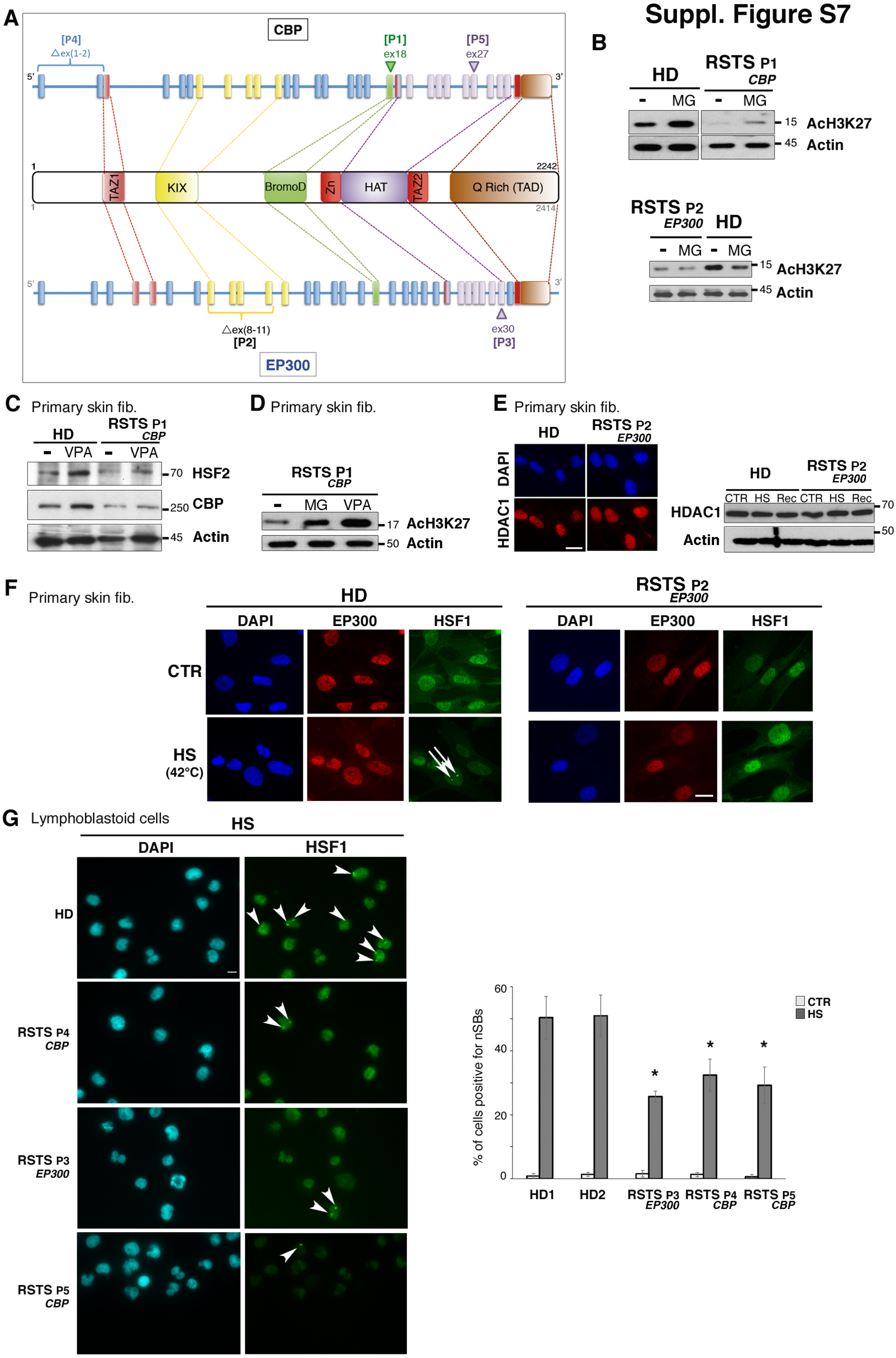
