## Supplemental Table S1 for "CBP/EP300 acetylates and stabilizes the stress-responsive Heat Shock Factor 2, a process compromised in Rubinstein-Taybi syndrome"

**Table S1. Summary of the acetylated peptides found in the mass spectrometry analysis of the mHSF2b isoform.** (relative to Figure 2C and Figure S2A)

The acetylated lysine residue and its position are highlighted in bold. The peptides are ranked from the N-terminal to C-terminal extremities of the mHSF2**b** protein. Some of the peptides are listed more than one time because they have been found several times in the same M/S experiment. Peptides have been attributed numbers (#), ranked from their N-terminal extremity of the HSF2 protein (relative to Figure S2A). Colors correspond to the HSF2 protein domains, schematized in Figure 2C.

| # | **Sequence (Acetylated lysine in bold)** | **Position** of the peptide**/ acetylated lysine** | **Domain of localization of the acetylated K** |
|---|---|---|---|
| 1 | (K)VVHIESGII**K**QER(D) | 72-86/**82** | DBD |
| 2 | (K)IRQEDLT**K**IISSAQK(V) | 120-136/**128** | HR-AB |
| 3 | (R)QEDLT**K**IISSAQK(V) | 122-136/**128** | |
| 4 | (R)QEDLT**K**IISSAQK(V) | 122-136/**128** | |
| 5 | (R)QEDLT**K**IISSAQ**K**VQIK(Q) | 122-139/**128**/**135** | |
| 6 | (R)QEDLT**K**IISSAQ**K**VQIK(Q) | 122-139/**128**/**135** | |
| 7 | (K)IISSAQ**K**VQIK(Q) | 128-140/**135** | |
| 8 | (R)**K**RPLLLNTNGAPK(K) | 196-210/**197** | |
| 9 | (R)KRPLLLNTNGAP**K**K(N) | 196-210/**209** | Just downstream HR-A/B |
| 10 | (K)RPLLLNTNGAP**KK**NLYQHIVK(E) | 197-218/**209**/**210** | |
| 11 | (K)RPLLLNTNGAP**KK**NLYQHIVK(E) | 197-218/**209**/**210** | |
| 12 | (K)**K**NLYQHIVK(E) | 209-219/**210** | |
| 13 | (R)QFSIDPDLLVDSEN**K**GLEATK(S) | 381-401/**395** | |
| 14 | (R)QFSIDPDLLVDSEN**K**GLEATK(S) | 381-401/**395** | |
| 15 | (R)QFSIDPDLLVDSENKGLEAT**K**SSVVQHVSEEGR(K) | 380-415/**401** | |
| 16 | (R)QFSIDPDLLVDSENKGLEAT**K**SSVVQHVSEEGR(K) | 380-415/**401** | |
| 17 | (K)GLEAT**K**SSVVQHVSEEGR(K) | 395-414/**401** | Between HR-C and before AD |
| 18 | (K)GLEAT**K**SSVVQHVSEEGRK(S) | 395-415/**401** | |
