## Supplemental Table S2 for "CBP/EP300 acetylates and stabilizes the stress-responsive Heat Shock Factor 2, a process compromised in Rubinstein-Taybi syndrome"

**Table S2 : Original tables and spectra corresponding to the HSF2 acetylated peptides identified by MS**

# #1 K82

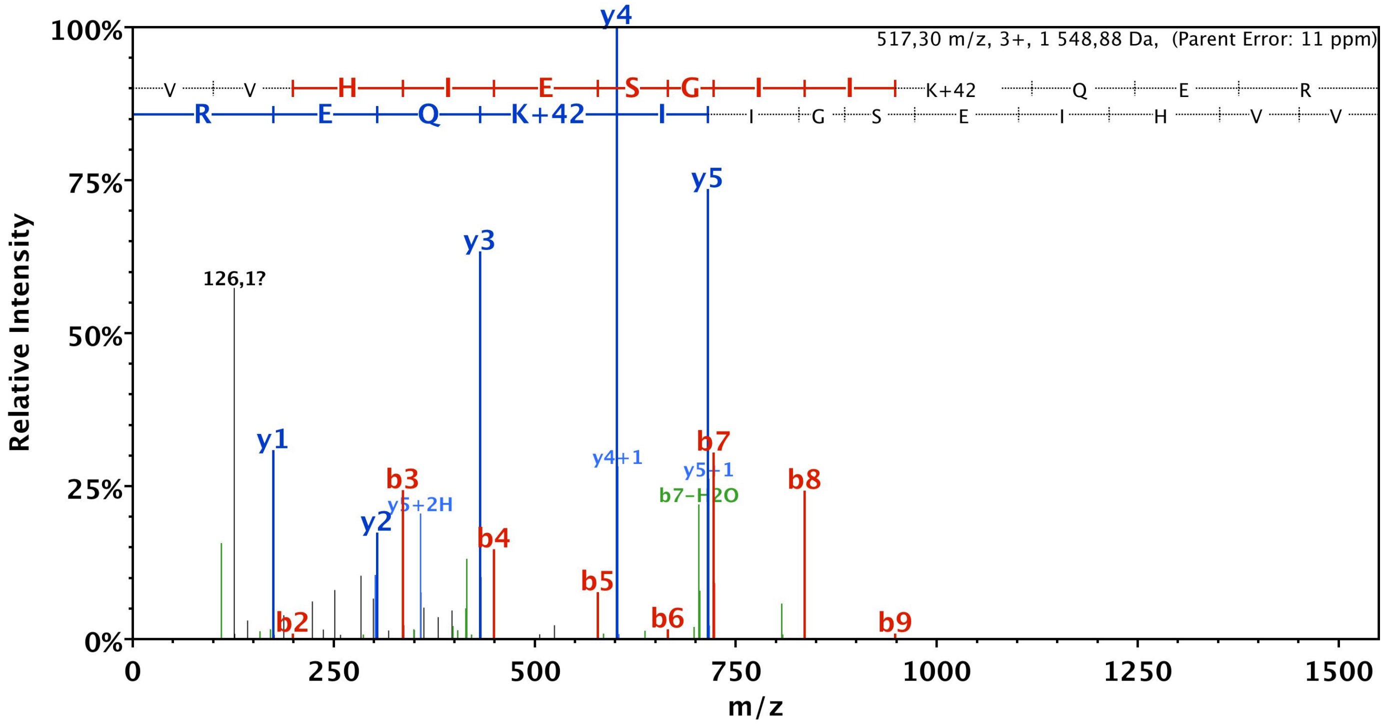


| B | B Ions | B+2H | B-NH3 | B-H2O | AA | Y Ions | Y+2H | Y-NH3 | Y-H2O | Y |
|---|---|---|---|---|---|---|---|---|---|---|
| 1 | 100,1 | 50,5 | | | V | 1 549,9 | 775,4 | 1 532,8 | 1 531,9 | 13 |
| 2 | 199,1 | 100,1 | | | V | 1 450,8 | 725,9 | 1 433,8 | 1 432,8 | 12 |
| 3 | 336,2 | 168,6 | | | H | 1 351,7 | 676,4 | 1 334,7 | 1 333,7 | 11 |
| 4 | 449,3 | 225,1 | | | I | 1 214,7 | 607,8 | 1 197,6 | 1 196,7 | 10 |
| 5 | 578,3 | 289,7 | | 560,3 | E | 1 101,6 | 551,3 | 1 084,6 | 1 083,6 | 9 |
| 6 | 665,4 | 333,2 | | 647,4 | S | 972,5 | 486,8 | 955,5 | 954,5 | 8 |
| 7 | 722,4 | 361,7 | | 704,4 | G | 885,5 | 443,3 | 868,5 | 867,5 | 7 |
| 8 | 835,5 | 418,2 | | 817,5 | I | 828,5 | 414,8 | 811,5 | 810,5 | 6 |
| 9 | 948,6 | 474,8 | | 930,5 | I | 715,4 | 358,2 | 698,4 | 697,4 | 5 |
| 10 | 1 118,7 | 559,8 | 1 101,6 | 1 100,6 | K+42 | 602,3 | 301,7 | 585,3 | 584,3 | 4 |
| 11 | 1 246,7 | 623,9 | 1 229,7 | 1 228,7 | Q | 432,2 | 216,6 | 415,2 | 414,2 | 3 |
| 12 | 1 375,8 | 688,4 | 1 358,7 | 1 357,7 | E | 304,2 | 152,6 | 287,1 | 286,2 | 2 |
| 13 | 1 549,9 | 775,4 | 1 532,8 | 1 531,9 | R | 175,1 | 88,1 | 158,1 | | 1 |

# #2 K128

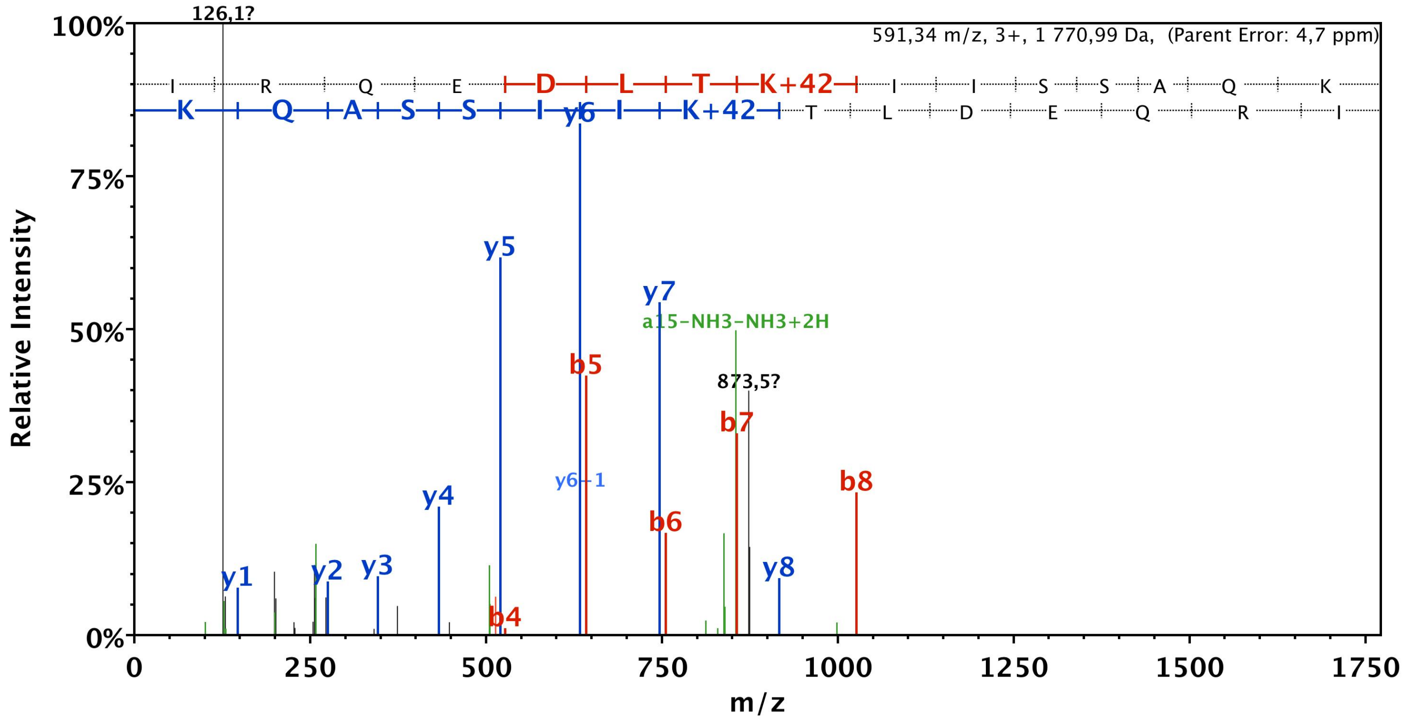


| B | B Ions | B+2H | B-NH3 | B-H2O | AA | Y Ions | Y+2H | Y-NH3 | Y-H2O | Y |
|---|---|---|---|---|---|---|---|---|---|---|
| 1 | 114,1 | 57,5 | | | I | 1 772,0 | 886,5 | 1 755,0 | 1 754,0 | 15 |
| 2 | 270,2 | 135,6 | 253,2 | | R | 1 658,9 | 830,0 | 1 641,9 | 1 640,9 | 14 |
| 3 | 398,3 | 199,6 | 381,2 | | Q | 1 502,8 | 751,9 | 1 485,8 | 1 484,8 | 13 |
| 4 | 527,3 | 264,2 | 510,3 | 509,3 | E | 1 374,7 | 687,9 | 1 357,7 | 1 356,7 | 12 |
| 5 | 642,3 | 321,7 | 625,3 | 624,3 | D | 1 245,7 | 623,4 | 1 228,7 | 1 227,7 | 11 |
| 6 | 755,4 | 378,2 | 738,4 | 737,4 | L | 1 130,7 | 565,8 | 1 113,7 | 1 112,7 | 10 |
| 7 | 856,5 | 428,7 | 839,4 | 838,4 | T | 1 017,6 | 509,3 | 1 000,6 | 999,6 | 9 |
| 8 | 1 026,6 | 513,8 | 1 009,5 | 1 008,5 | K+42 | 916,5 | 458,8 | 899,5 | 898,5 | 8 |
| 9 | 1 139,6 | 570,3 | 1 122,6 | 1 121,6 | I | 746,4 | 373,7 | 729,4 | 728,4 | 7 |
| 10 | 1 252,7 | 626,9 | 1 235,7 | 1 234,7 | I | 633,4 | 317,2 | 616,3 | 615,3 | 6 |
| 11 | 1 339,8 | 670,4 | 1 322,7 | 1 321,7 | S | 520,3 | 260,6 | 503,2 | 502,3 | 5 |
| 12 | 1 426,8 | 713,9 | 1 409,8 | 1 408,8 | S | 433,2 | 217,1 | 416,2 | 415,2 | 4 |
| 13 | 1 497,8 | 749,4 | 1 480,8 | 1 479,8 | A | 346,2 | 173,6 | 329,2 | | 3 |
| 14 | 1 625,9 | 813,4 | 1 608,9 | 1 607,9 | Q | 275,2 | 138,1 | 258,1 | | 2 |
| 15 | 1 772,0 | 886,5 | 1 755,0 | 1 754,0 | K | 147,1 | 74,1 | 130,1 | | 1 |

# #3 K128

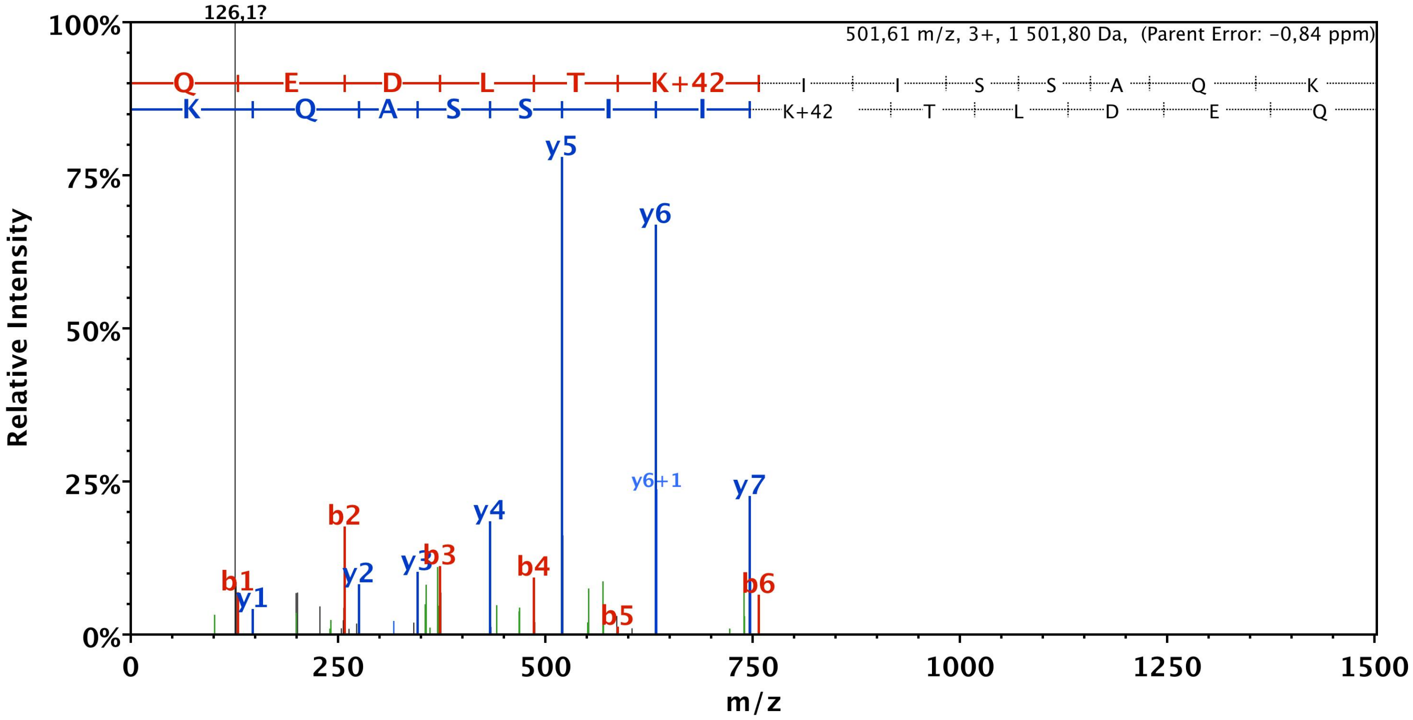


| B | B Ions | B+2H | B-NH3 | B-H2O | AA | Y Ions | Y+2H | Y-NH3 | Y-H2O | Y |
|---|---|---|---|---|---|---|---|---|---|---|
| 1 | 129,1 | 65,0 | 112,0 | | Q | 1 502,8 | 751,9 | 1 485,8 | 1 484,8 | 13 |
| 2 | 258,1 | 129,6 | 241,1 | 240,1 | E | 1 374,7 | 687,9 | 1 357,7 | 1 356,7 | 12 |
| 3 | 373,1 | 187,1 | 356,1 | 355,1 | D | 1 245,7 | 623,4 | 1 228,7 | 1 227,7 | 11 |
| 4 | 486,2 | 243,6 | 469,2 | 468,2 | L | 1 130,7 | 565,8 | 1 113,7 | 1 112,7 | 10 |
| 5 | 587,3 | 294,1 | 570,2 | 569,3 | T | 1 017,6 | 509,3 | 1 000,6 | 999,6 | 9 |
| 6 | 757,4 | 379,2 | 740,3 | 739,4 | K+42 | 916,5 | 458,8 | 899,5 | 898,5 | 8 |
| 7 | 870,5 | 435,7 | 853,4 | 852,4 | I | 746,4 | 373,7 | 729,4 | 728,4 | 7 |
| 8 | 983,5 | 492,3 | 966,5 | 965,5 | I | 633,4 | 317,2 | 616,3 | 615,3 | 6 |
| 9 | 1 070,6 | 535,8 | 1 053,5 | 1 052,6 | S | 520,3 | 260,6 | 503,2 | 502,3 | 5 |
| 10 | 1 157,6 | 579,3 | 1 140,6 | 1 139,6 | S | 433,2 | 217,1 | 416,2 | 415,2 | 4 |
| 11 | 1 228,6 | 614,8 | 1 211,6 | 1 210,6 | A | 346,2 | 173,6 | 329,2 | | 3 |
| 12 | 1 356,7 | 678,9 | 1 339,7 | 1 338,7 | Q | 275,2 | 138,1 | 258,1 | | 2 |
| 13 | 1 502,8 | 751,9 | 1 485,8 | 1 484,8 | K | 147,1 | 74,1 | 130,1 | | 1 |

# #4 K128

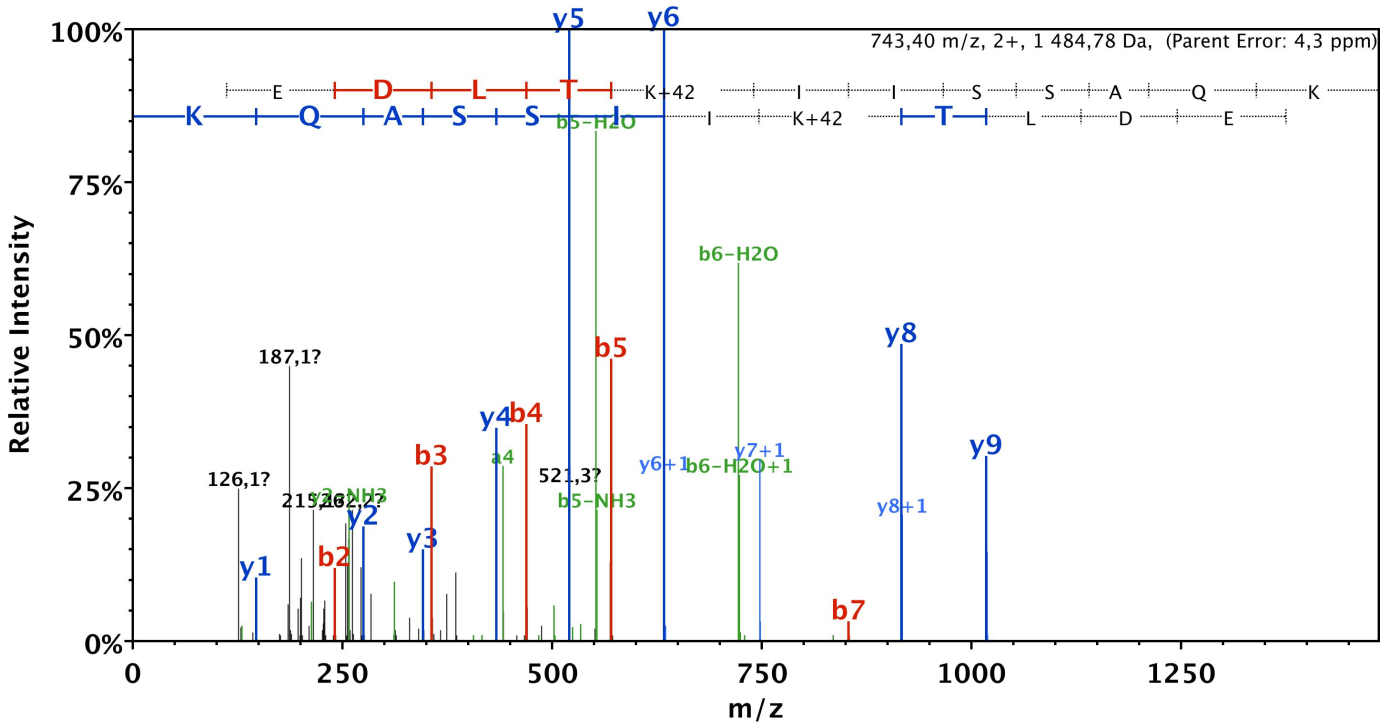


| B | B Ions | B+2H | B-NH3 | B-H2O | AA | Y Ions | Y+2H | Y-NH3 | Y-H2O | Y |
|---|---|---|---|---|---|---|---|---|---|---|
| 1 | 112,0 | | 95,0 | | Q-17 | 1 485,8 | 743,4 | 1 468,8 | 1 467,8 | 13 |
| 2 | 241,1 | | 224,1 | 223,1 | E | 1 374,7 | 687,9 | 1 357,7 | 1 356,7 | 12 |
| 3 | 356,1 | | 339,1 | 338,1 | D | 1 245,7 | 623,4 | 1 228,7 | 1 227,7 | 11 |
| 4 | 469,2 | | 452,2 | 451,2 | L | 1 130,7 | 565,8 | 1 113,7 | 1 112,7 | 10 |
| 5 | 570,2 | | 553,2 | 552,2 | T | 1 017,6 | 509,3 | 1 000,6 | 999,6 | 9 |
| 6 | 740,3 | 370,7 | 723,3 | 722,3 | K+42 | 916,5 | 458,8 | 899,5 | 898,5 | 8 |
| 7 | 853,4 | 427,2 | 836,4 | 835,4 | I | 746,4 | 373,7 | 729,4 | 728,4 | 7 |
| 8 | 966,5 | 483,8 | 949,5 | 948,5 | I | 633,4 | 317,2 | 616,3 | 615,3 | 6 |
| 9 | 1 053,5 | 527,3 | 1 036,5 | 1 035,5 | S | 520,3 | | 503,2 | 502,3 | 5 |
| 10 | 1 140,6 | 570,8 | 1 123,6 | 1 122,6 | S | 433,2 | | 416,2 | 415,2 | 4 |
| 11 | 1 211,6 | 606,3 | 1 194,6 | 1 193,6 | A | 346,2 | | 329,2 | | 3 |
| 12 | 1 339,7 | 670,3 | 1 322,6 | 1 321,7 | Q | 275,2 | | 258,1 | | 2 |
| 13 | 1 485,8 | 743,4 | 1 468,8 | 1 467,8 | K | 147,1 | | 130,1 | | 1 |

# #5 K128/135

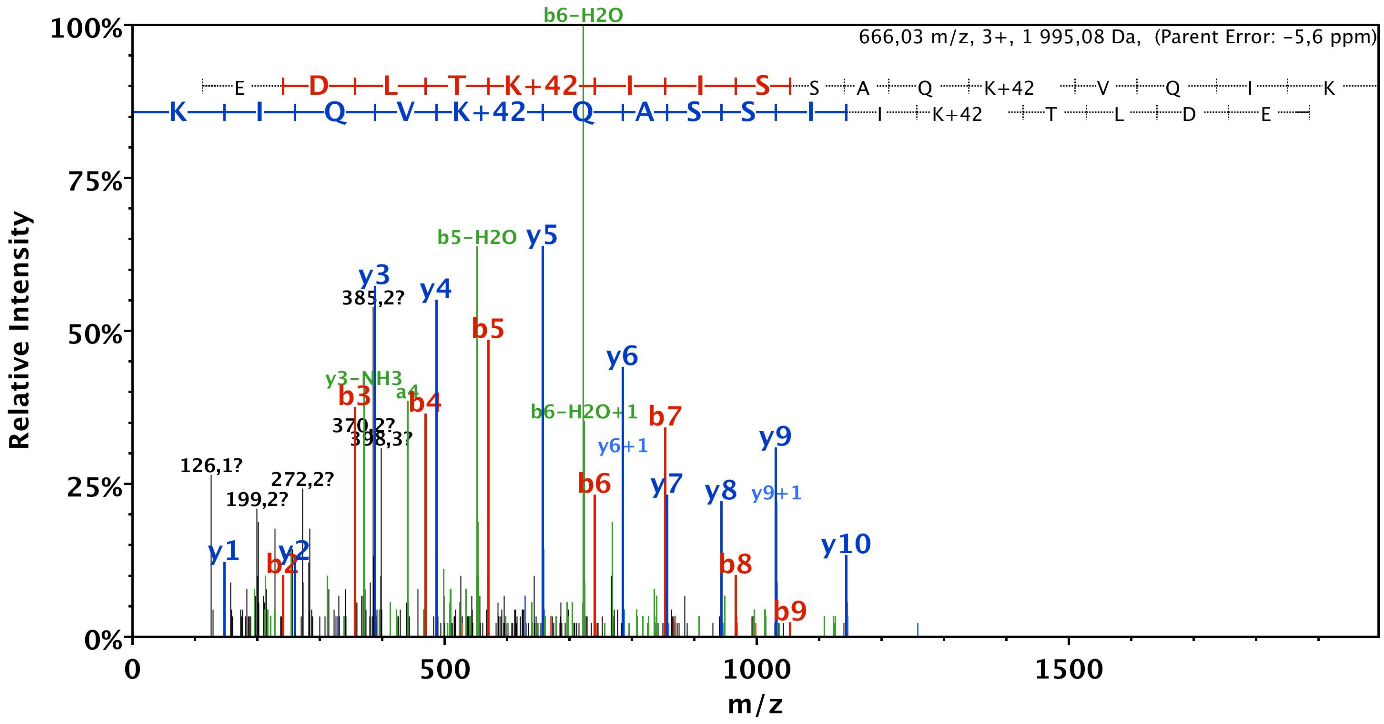


| B | B Ions | B+2H | B-NH3 | B-H2O | AA | Y Ions | Y+2H | Y-NH3 | Y-H2O | Y |
|---|---|---|---|---|---|---|---|---|---|---|
| 1 | 112,0 | 56,5 | 95,0 | | Q-17 | 1 996,1 | 998,6 | 1 979,1 | 1 978,1 | 17 |
| 2 | 241,1 | 121,0 | 224,1 | 223,1 | E | 1 885,1 | 943,0 | 1 868,0 | 1 867,1 | 16 |
| 3 | 356,1 | 178,6 | 339,1 | 338,1 | D | 1 756,0 | 878,5 | 1 739,0 | 1 738,0 | 15 |
| 4 | 469,2 | 235,1 | 452,2 | 451,2 | L | 1 641,0 | 821,0 | 1 624,0 | 1 623,0 | 14 |
| 5 | 570,2 | 285,6 | 553,2 | 552,2 | T | 1 527,9 | 764,5 | 1 510,9 | 1 509,9 | 13 |
| 6 | 740,3 | 370,7 | 723,3 | 722,3 | K+42 | 1 426,9 | 713,9 | 1 409,8 | 1 408,9 | 12 |
| 7 | 853,4 | 427,2 | 836,4 | 835,4 | I | 1 256,8 | 628,9 | 1 239,7 | 1 238,7 | 11 |
| 8 | 966,5 | 483,8 | 949,5 | 948,5 | I | 1 143,7 | 572,3 | 1 126,6 | 1 125,7 | 10 |
| 9 | 1 053,5 | 527,3 | 1 036,5 | 1 035,5 | S | 1 030,6 | 515,8 | 1 013,6 | 1 012,6 | 9 |
| 10 | 1 140,6 | 570,8 | 1 123,6 | 1 122,6 | S | 943,6 | 472,3 | 926,5 | 925,5 | 8 |
| 11 | 1 211,6 | 606,3 | 1 194,6 | 1 193,6 | A | 856,5 | 428,8 | 839,5 | | 7 |
| 12 | 1 339,7 | 670,3 | 1 322,6 | 1 321,7 | Q | 785,5 | 393,2 | 768,5 | | 6 |
| 13 | 1 509,8 | 755,4 | 1 492,8 | 1 491,8 | K+42 | 657,4 | 329,2 | 640,4 | | 5 |
| 14 | 1 608,8 | 804,9 | 1 591,8 | 1 590,8 | V | 487,3 | 244,2 | 470,3 | | 4 |
| 15 | 1 736,9 | 869,0 | 1 719,9 | 1 718,9 | Q | 388,3 | 194,6 | 371,2 | | 3 |
| 16 | 1 850,0 | 925,5 | 1 833,0 | 1 832,0 | I | 260,2 | 130,6 | 243,2 | | 2 |
| 17 | 1 996,1 | 998,6 | 1 979,1 | 1 978,1 | K | 147,1 | 74,1 | 130,1 | | 1 |

# #6 K128/135

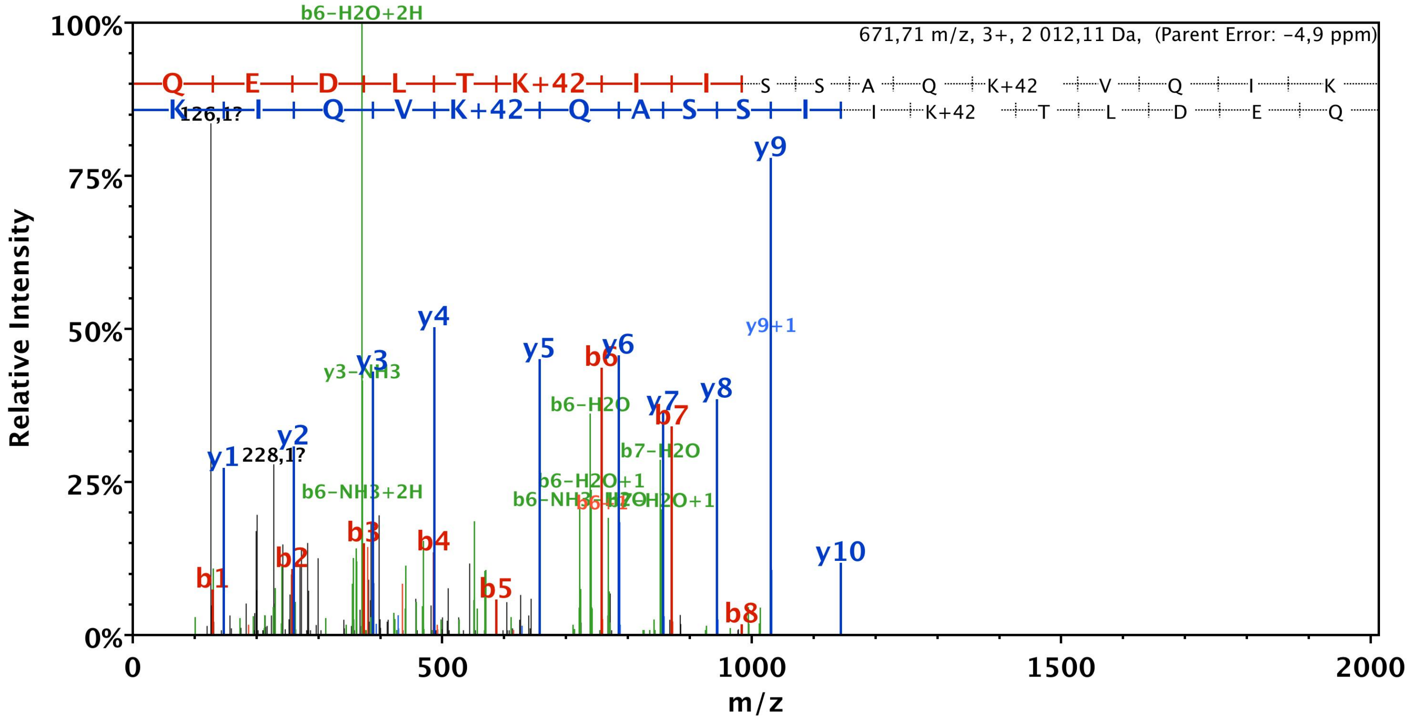


| B | B Ions | B+2H | B-NH3 | B-H2O | AA | Y Ions | Y+2H | Y-NH3 | Y-H2O | Y |
|---|---|---|---|---|---|---|---|---|---|---|
| 1 | 129,1 | 65,0 | 112,0 | | Q | 2 013,1 | 1 007,1 | 1 996,1 | 1 995,1 | 17 |
| 2 | 258,1 | 129,6 | 241,1 | 240,1 | E | 1 885,1 | 943,0 | 1 868,0 | 1 867,1 | 16 |
| 3 | 373,1 | 187,1 | 356,1 | 355,1 | D | 1 756,0 | 878,5 | 1 739,0 | 1 738,0 | 15 |
| 4 | 486,2 | 243,6 | 469,2 | 468,2 | L | 1 641,0 | 821,0 | 1 624,0 | 1 623,0 | 14 |
| 5 | 587,3 | 294,1 | 570,2 | 569,3 | T | 1 527,9 | 764,5 | 1 510,9 | 1 509,9 | 13 |
| 6 | 757,4 | 379,2 | 740,3 | 739,4 | K+42 | 1 426,9 | 713,9 | 1 409,8 | 1 408,9 | 12 |
| 7 | 870,5 | 435,7 | 853,4 | 852,4 | I | 1 256,8 | 628,9 | 1 239,7 | 1 238,7 | 11 |
| 8 | 983,5 | 492,3 | 966,5 | 965,5 | I | 1 143,7 | 572,3 | 1 126,6 | 1 125,7 | 10 |
| 9 | 1 070,6 | 535,8 | 1 053,5 | 1 052,6 | S | 1 030,6 | 515,8 | 1 013,6 | 1 012,6 | 9 |
| 10 | 1 157,6 | 579,3 | 1 140,6 | 1 139,6 | S | 943,6 | 472,3 | 926,5 | 925,5 | 8 |
| 11 | 1 228,6 | 614,8 | 1 211,6 | 1 210,6 | A | 856,5 | 428,8 | 839,5 | | 7 |
| 12 | 1 356,7 | 678,9 | 1 339,7 | 1 338,7 | Q | 785,5 | 393,2 | 768,5 | | 6 |
| 13 | 1 526,8 | 763,9 | 1 509,8 | 1 508,8 | K+42 | 657,4 | 329,2 | 640,4 | | 5 |
| 14 | 1 625,9 | 813,4 | 1 608,8 | 1 607,9 | V | 487,3 | 244,2 | 470,3 | | 4 |
| 15 | 1 753,9 | 877,5 | 1 736,9 | 1 735,9 | Q | 388,3 | 194,6 | 371,2 | | 3 |
| 16 | 1 867,0 | 934,0 | 1 850,0 | 1 849,0 | I | 260,2 | 130,6 | 243,2 | | 2 |
| 17 | 2 013,1 | 1 007,1 | 1 996,1 | 1 995,1 | K | 147,1 | 74,1 | 130,1 | | 1 |

# #7 K135

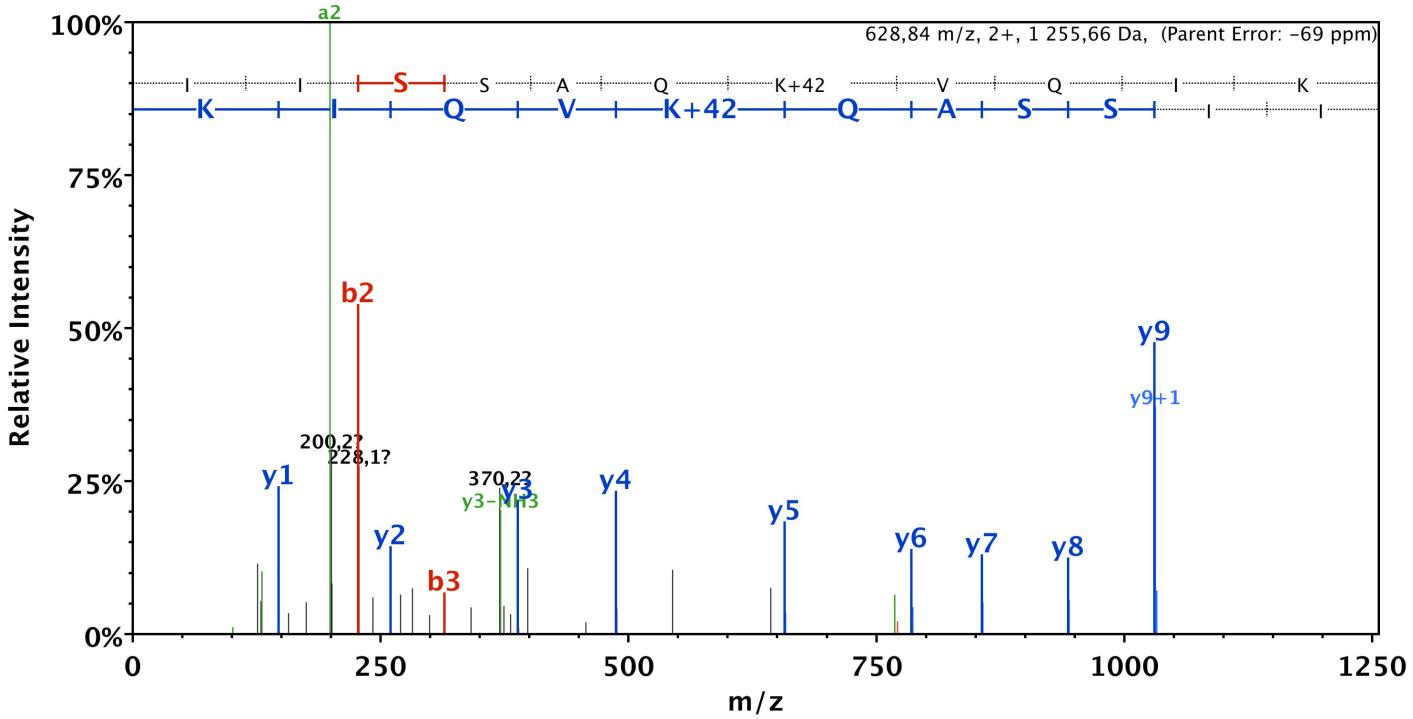


| B | B Ions | B+2H | B-NH3 | B-H2O | AA | Y Ions | Y+2H | Y-NH3 | Y-H2O | Y |
|---|---|---|---|---|---|---|---|---|---|---|
| 1 | 114,1 | | | | I | 1 256,8 | 628,9 | 1 239,7 | 1 238,7 | 11 |
| 2 | 227,2 | | | | I | 1 143,7 | 572,3 | 1 126,6 | 1 125,7 | 10 |
| 3 | 314,2 | | | 296,2 | S | 1 030,6 | 515,8 | 1 013,6 | 1 012,6 | 9 |
| 4 | 401,2 | | | 383,2 | S | 943,6 | 472,3 | 926,5 | 925,5 | 8 |
| 5 | 472,3 | | | 454,3 | A | 856,5 | 428,8 | 839,5 | | 7 |
| 6 | 600,3 | 300,7 | 583,3 | 582,3 | Q | 785,5 | 393,2 | 768,5 | | 6 |
| 7 | 770,4 | 385,7 | 753,4 | 752,4 | K+42 | 657,4 | 329,2 | 640,4 | | 5 |
| 8 | 869,5 | 435,3 | 852,5 | 851,5 | V | 487,3 | | 470,3 | | 4 |
| 9 | 997,6 | 499,3 | 980,5 | 979,6 | Q | 388,3 | | 371,2 | | 3 |
| 10 | 1 110,7 | 555,8 | 1 093,6 | 1 092,6 | I | 260,2 | | 243,2 | | 2 |
| 11 | 1 256,8 | 628,9 | 1 239,7 | 1 238,7 | K | 147,1 | | 130,1 | | 1 |

# #8 K197

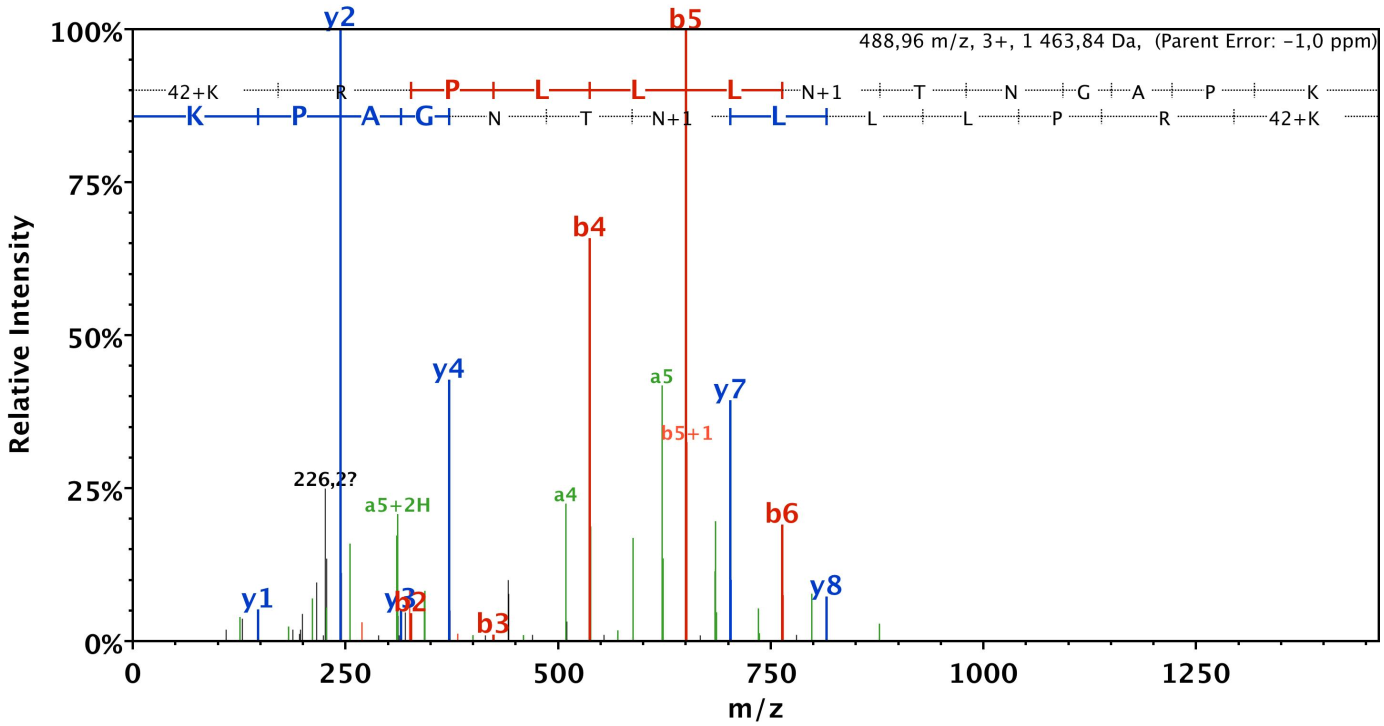


| B | B Ions | B+2H | B-NH3 | B-H2O | AA | Y Ions | Y+2H | Y-NH3 | Y-H2O | Y |
|---|---|---|---|---|---|---|---|---|---|---|
| 1 | 171,1 | 86,1 | 154,1 | | K+42 | 1 464,9 | 732,9 | 1 447,8 | 1 446,8 | 13 |
| 2 | 327,2 | 164,1 | 310,2 | | R | 1 294,7 | 647,9 | 1 277,7 | 1 276,7 | 12 |
| 3 | 424,3 | 212,6 | 407,2 | | P | 1 138,6 | 569,8 | 1 121,6 | 1 120,6 | 11 |
| 4 | 537,4 | 269,2 | 520,3 | | L | 1 041,6 | 521,3 | 1 024,6 | 1 023,6 | 10 |
| 5 | 650,4 | 325,7 | 633,4 | | L | 928,5 | 464,8 | 911,5 | 910,5 | 9 |
| 6 | 763,5 | 382,3 | 746,5 | | L | 815,4 | 408,2 | 798,4 | 797,4 | 8 |
| 7 | 878,5 | 439,8 | 861,5 | | N+1 | 702,3 | 351,7 | 685,3 | 684,3 | 7 |
| 8 | 979,6 | 490,3 | 962,6 | 961,6 | T | 587,3 | 294,2 | 570,3 | 569,3 | 6 |
| 9 | 1 093,6 | 547,3 | 1 076,6 | 1 075,6 | N | 486,3 | 243,6 | 469,2 | | 5 |
| 10 | 1 150,7 | 575,8 | 1 133,6 | 1 132,6 | G | 372,2 | 186,6 | 355,2 | | 4 |
| 11 | 1 221,7 | 611,4 | 1 204,7 | 1 203,7 | A | 315,2 | 158,1 | 298,2 | | 3 |
| 12 | 1 318,7 | 659,9 | 1 301,7 | 1 300,7 | P | 244,2 | 122,6 | 227,1 | | 2 |
| 13 | 1 464,9 | 732,9 | 1 447,8 | 1 446,8 | K | 147,1 | 74,1 | 130,1 | | 1 |

# #9 K209

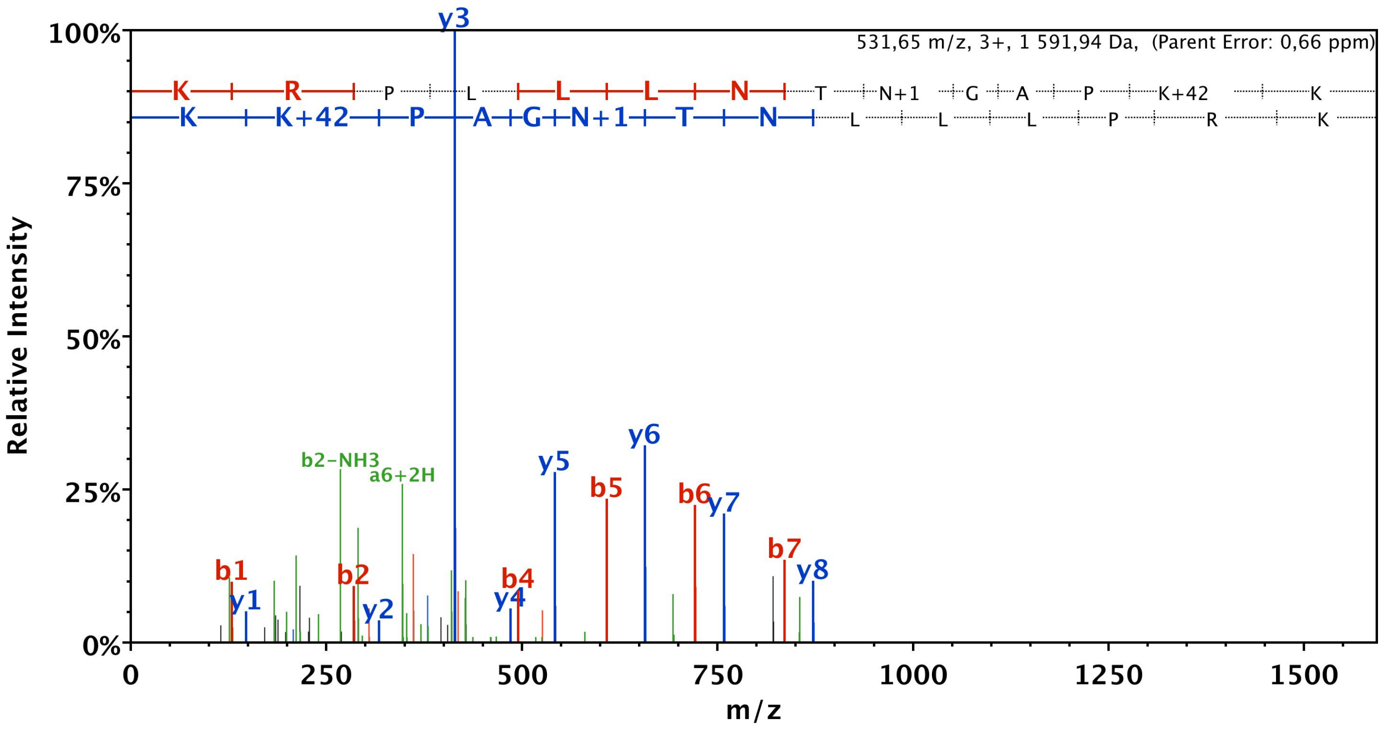


| B | B Ions | B+2H | B-NH3 | B-H2O | AA | Y Ions | Y+2H | Y-NH3 | Y-H2O | Y |
|---|---|---|---|---|---|---|---|---|---|---|
| 1 | 129,1 | 65,1 | 112,1 | | K | 1 592,9 | 797,0 | 1 575,9 | 1 574,9 | 14 |
| 2 | 285,2 | 143,1 | 268,2 | | R | 1 464,9 | 732,9 | 1 447,8 | 1 446,8 | 13 |
| 3 | 382,3 | 191,6 | 365,2 | | P | 1 308,8 | 654,9 | 1 291,7 | 1 290,7 | 12 |
| 4 | 495,3 | 248,2 | 478,3 | | L | 1 211,7 | 606,4 | 1 194,7 | 1 193,7 | 11 |
| 5 | 608,4 | 304,7 | 591,4 | | L | 1 098,6 | 549,8 | 1 081,6 | 1 080,6 | 10 |
| 6 | 721,5 | 361,3 | 704,5 | | L | 985,5 | 493,3 | 968,5 | 967,5 | 9 |
| 7 | 835,6 | 418,3 | 818,5 | | N | 872,4 | 436,7 | 855,4 | 854,4 | 8 |
| 8 | 936,6 | 468,8 | 919,6 | 918,6 | T | 758,4 | 379,7 | 741,4 | 740,4 | 7 |
| 9 | 1 051,6 | 526,3 | 1 034,6 | 1 033,6 | N+1 | 657,4 | 329,2 | 640,3 | | 6 |
| 10 | 1 108,6 | 554,8 | 1 091,6 | 1 090,6 | G | 542,3 | 271,7 | 525,3 | | 5 |
| 11 | 1 179,7 | 590,3 | 1 162,7 | 1 161,7 | A | 485,3 | 243,2 | 468,3 | | 4 |
| 12 | 1 276,7 | 638,9 | 1 259,7 | 1 258,7 | P | 414,3 | 207,6 | 397,2 | | 3 |
| 13 | 1 446,8 | 723,9 | 1 429,8 | 1 428,8 | K+42 | 317,2 | 159,1 | 300,2 | | 2 |
| 14 | 1 592,9 | 797,0 | 1 575,9 | 1 574,9 | K | 147,1 | 74,1 | 130,1 | | 1 |

# #10 K209/210

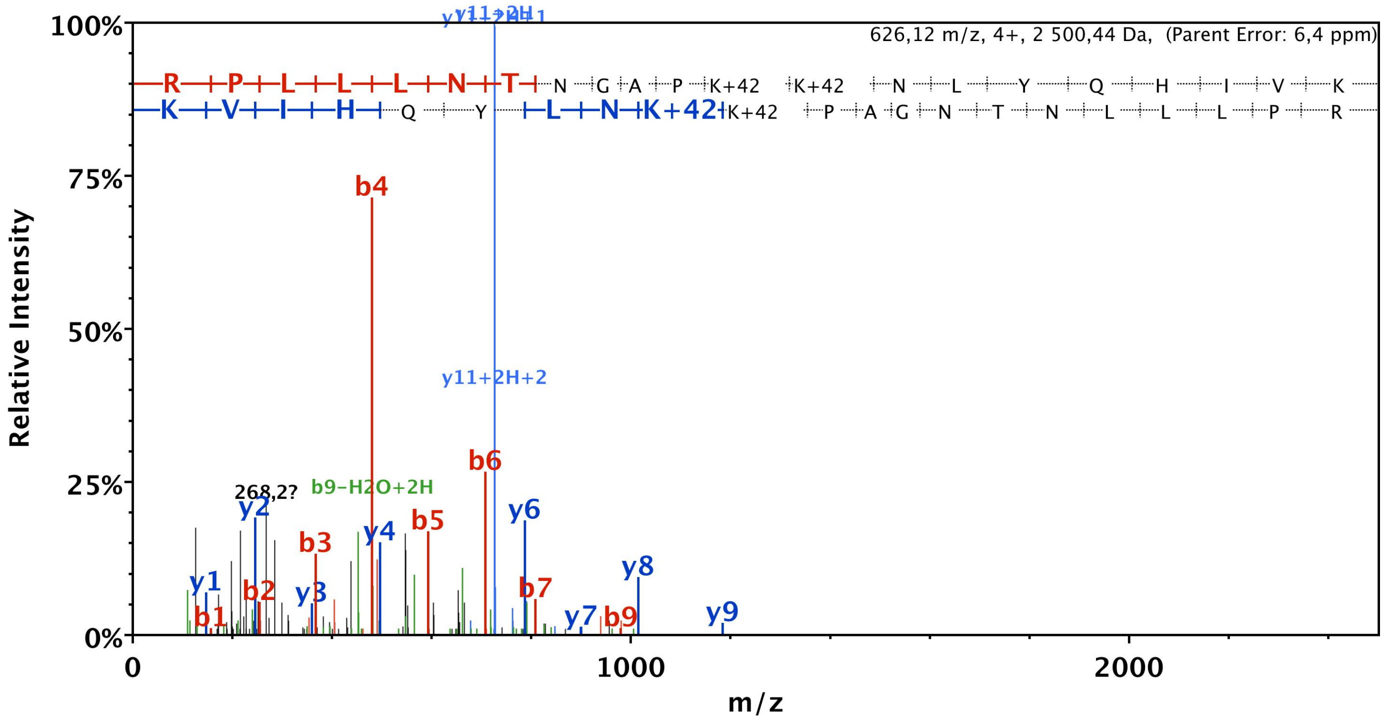


| B | B Ions | B+2H | B-NH3 | B-H2O | AA | Y Ions | Y+2H | Y-NH3 | Y-H2O | Y |
|---|---|---|---|---|---|---|---|---|---|---|
| 1 | 157,1 | 79,1 | 140,1 | | R | 2 501,4 | 1 251,2 | 2 484,4 | 2 483,4 | 21 |
| 2 | 254,2 | 127,6 | 237,1 | | P | 2 345,3 | 1 173,2 | 2 328,3 | 2 327,3 | 20 |
| 3 | 367,2 | 184,1 | 350,2 | | L | 2 248,3 | 1 124,6 | 2 231,3 | 2 230,3 | 19 |
| 4 | 480,3 | 240,7 | 463,3 | | L | 2 135,2 | 1 068,1 | 2 118,2 | 2 117,2 | 18 |
| 5 | 593,4 | 297,2 | 576,4 | | L | 2 022,1 | 1 011,6 | 2 005,1 | 2 004,1 | 17 |
| 6 | 707,5 | 354,2 | 690,4 | | N | 1 909,0 | 955,0 | 1 892,0 | 1 891,0 | 16 |
| 7 | 808,5 | 404,8 | 791,5 | 790,5 | T | 1 795,0 | 898,0 | 1 778,0 | 1 777,0 | 15 |
| 8 | 922,5 | 461,8 | 905,5 | 904,5 | N | 1 693,9 | 847,5 | 1 676,9 | | 14 |
| 9 | 979,6 | 490,3 | 962,5 | 961,6 | G | 1 579,9 | 790,5 | 1 562,9 | | 13 |
| 10 | 1 050,6 | 525,8 | 1 033,6 | 1 032,6 | A | 1 522,9 | 761,9 | 1 505,8 | | 12 |
| 11 | 1 147,7 | 574,3 | 1 130,6 | 1 129,6 | P | 1 451,8 | 726,4 | 1 434,8 | | 11 |
| 12 | 1 317,8 | 659,4 | 1 300,7 | 1 299,8 | K+42 | 1 354,8 | 677,9 | 1 337,8 | | 10 |
| 13 | 1 487,9 | 744,4 | 1 470,8 | 1 469,9 | K+42 | 1 184,7 | 592,8 | 1 167,7 | | 9 |
| 14 | 1 601,9 | 801,5 | 1 584,9 | 1 583,9 | N | 1 014,6 | 507,8 | 997,5 | | 8 |
| 15 | 1 715,0 | 858,0 | 1 698,0 | 1 697,0 | L | 900,5 | 450,8 | 883,5 | | 7 |
| 16 | 1 878,1 | 939,5 | 1 861,0 | 1 860,0 | Y | 787,4 | 394,2 | 770,4 | | 6 |
| 17 | 2 006,1 | 1 003,6 | 1 989,1 | 1 988,1 | Q | 624,4 | 312,7 | 607,4 | | 5 |
| 18 | 2 143,2 | 1 072,1 | 2 126,2 | 2 125,2 | H | 496,3 | 248,7 | 479,3 | | 4 |
| 19 | 2 256,3 | 1 128,6 | 2 239,2 | 2 238,3 | I | 359,3 | 180,1 | 342,2 | | 3 |
| 20 | 2 355,3 | 1 178,2 | 2 338,3 | 2 337,3 | V | 246,2 | 123,6 | 229,2 | | 2 |
| 21 | 2 501,4 | 1 251,2 | 2 484,4 | 2 483,4 | K | 147,1 | 74,1 | 130,1 | | 1 |

# #11 K209/210

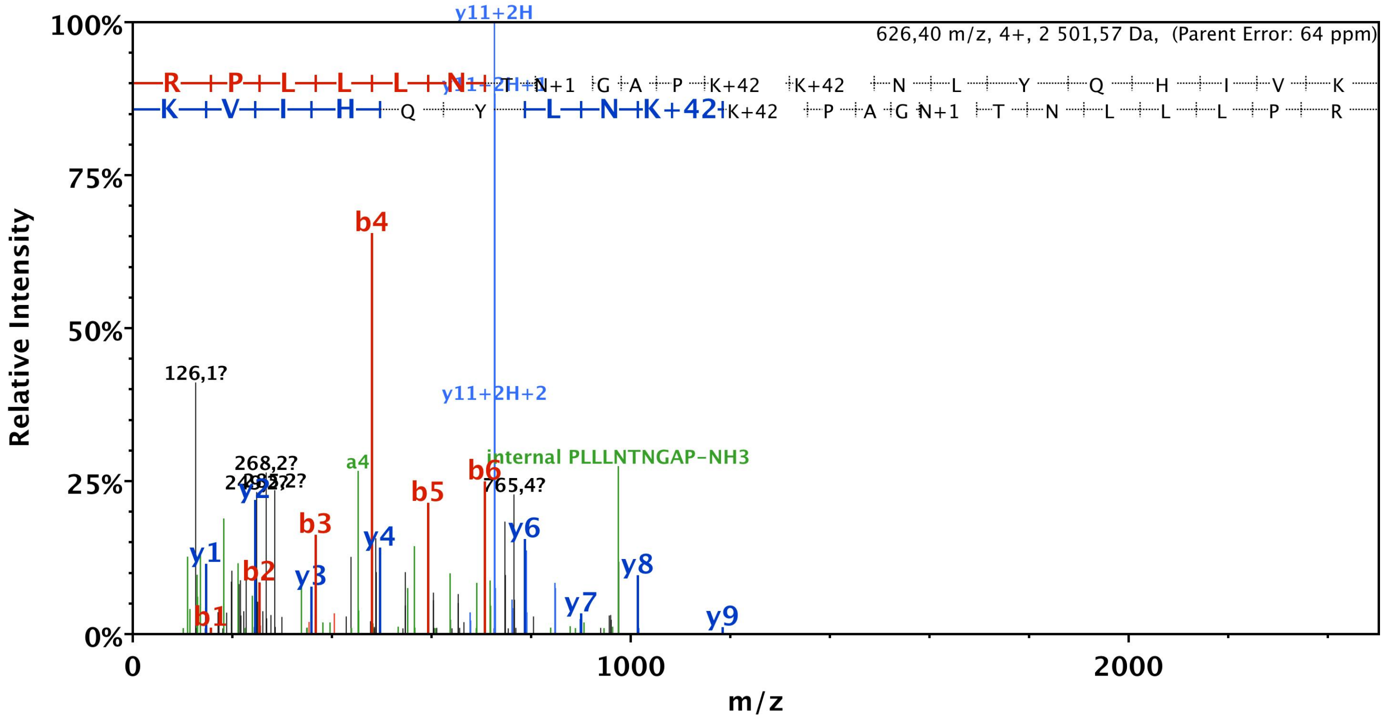


| B | B Ions | B+2H | B-NH3 | B-H2O | AA | Y Ions | Y+2H | Y-NH3 | Y-H2O | Y |
|---|---|---|---|---|---|---|---|---|---|---|
| 1 | 157,1 | 79,1 | 140,1 | | R | 2 502,4 | 1 251,7 | 2 485,4 | 2 484,4 | 21 |
| 2 | 254,2 | 127,6 | 237,1 | | P | 2 346,3 | 1 173,7 | 2 329,3 | 2 328,3 | 20 |
| 3 | 367,2 | 184,1 | 350,2 | | L | 2 249,3 | 1 125,1 | 2 232,2 | 2 231,3 | 19 |
| 4 | 480,3 | 240,7 | 463,3 | | L | 2 136,2 | 1 068,6 | 2 119,2 | 2 118,2 | 18 |
| 5 | 593,4 | 297,2 | 576,4 | | L | 2 023,1 | 1 012,1 | 2 006,1 | 2 005,1 | 17 |
| 6 | 707,5 | 354,2 | 690,4 | | N | 1 910,0 | 955,5 | 1 893,0 | 1 892,0 | 16 |
| 7 | 808,5 | 404,8 | 791,5 | 790,5 | T | 1 796,0 | 898,5 | 1 778,9 | 1 778,0 | 15 |
| 8 | 923,5 | 462,3 | 906,5 | 905,5 | N+1 | 1 694,9 | 848,0 | 1 677,9 | | 14 |
| 9 | 980,6 | 490,8 | 963,5 | 962,5 | G | 1 579,9 | 790,5 | 1 562,9 | | 13 |
| 10 | 1 051,6 | 526,3 | 1 034,6 | 1 033,6 | A | 1 522,9 | 761,9 | 1 505,8 | | 12 |
| 11 | 1 148,6 | 574,8 | 1 131,6 | 1 130,6 | P | 1 451,8 | 726,4 | 1 434,8 | | 11 |
| 12 | 1 318,7 | 659,9 | 1 301,7 | 1 300,7 | K+42 | 1 354,8 | 677,9 | 1 337,8 | | 10 |
| 13 | 1 488,9 | 744,9 | 1 471,8 | 1 470,8 | K+42 | 1 184,7 | 592,8 | 1 167,7 | | 9 |
| 14 | 1 602,9 | 802,0 | 1 585,9 | 1 584,9 | N | 1 014,6 | 507,8 | 997,5 | | 8 |
| 15 | 1 716,0 | 858,5 | 1 699,0 | 1 698,0 | L | 900,5 | 450,8 | 883,5 | | 7 |
| 16 | 1 879,0 | 940,0 | 1 862,0 | 1 861,0 | Y | 787,4 | 394,2 | 770,4 | | 6 |
| 17 | 2 007,1 | 1 004,1 | 1 990,1 | 1 989,1 | Q | 624,4 | 312,7 | 607,4 | | 5 |
| 18 | 2 144,2 | 1 072,6 | 2 127,1 | 2 126,2 | H | 496,3 | 248,7 | 479,3 | | 4 |
| 19 | 2 257,2 | 1 129,1 | 2 240,2 | 2 239,2 | I | 359,3 | 180,1 | 342,2 | | 3 |
| 20 | 2 356,3 | 1 178,7 | 2 339,3 | 2 338,3 | V | 246,2 | 123,6 | 229,2 | | 2 |
| 21 | 2 502,4 | 1 251,7 | 2 485,4 | 2 484,4 | K | 147,1 | 74,1 | 130,1 | | 1 |

# #12 K210

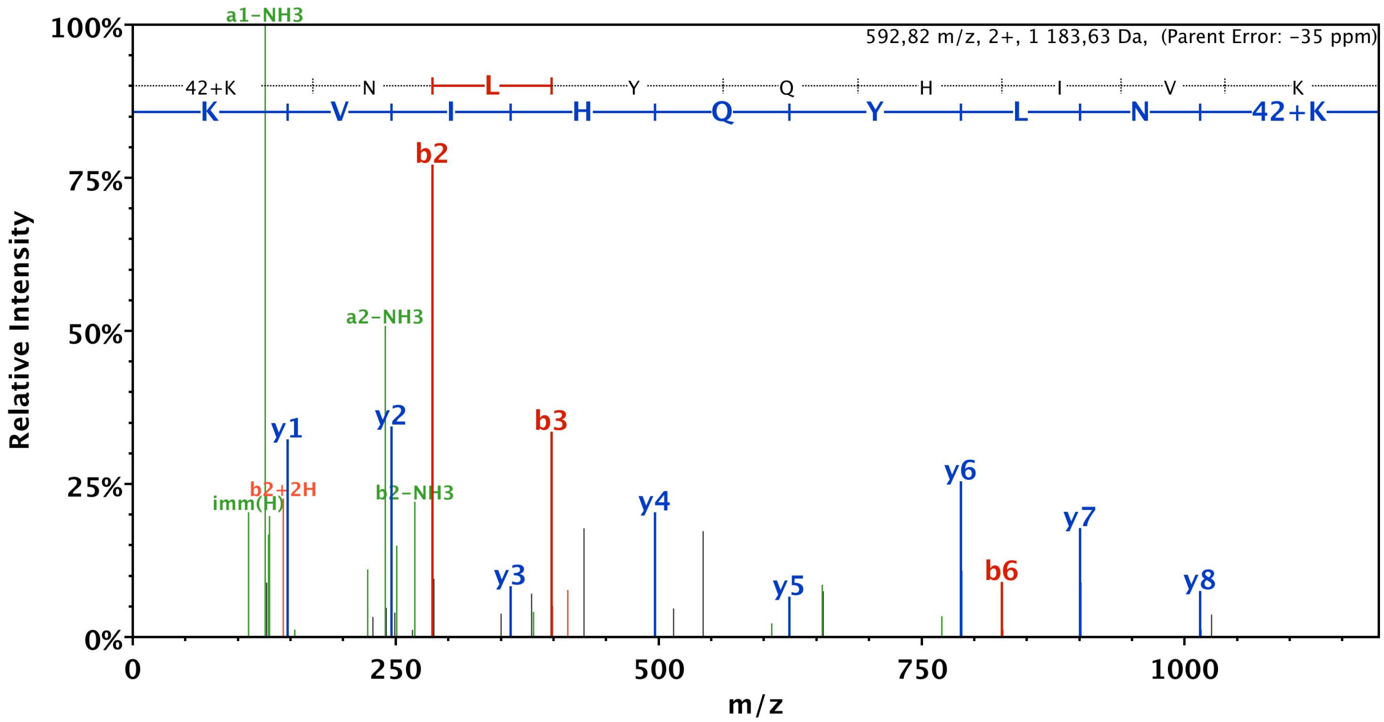


| B | B Ions | B+2H | B-NH3 | B-H2O | AA | Y Ions | Y+2H | Y-NH3 | Y-H2O | Y |
|---|---|---|---|---|---|---|---|---|---|---|
| 1 | 171,1 | 86,1 | 154,1 | | K+42 | 1 184,7 | 592,8 | 1 167,7 | 1 166,7 | 9 |
| 2 | 285,2 | 143,1 | 268,1 | | N | 1 014,6 | 507,8 | 997,5 | 996,6 | 8 |
| 3 | 398,2 | 199,6 | 381,2 | | L | 900,5 | 450,8 | 883,5 | 882,5 | 7 |
| 4 | 561,3 | 281,2 | 544,3 | 543,3 | Y | 787,4 | 394,2 | 770,4 | 769,4 | 6 |
| 5 | 689,4 | 345,2 | 672,3 | 671,4 | Q | 624,4 | 312,7 | 607,4 | 606,4 | 5 |
| 6 | 826,4 | 413,7 | 809,4 | 808,4 | H | 496,3 | 248,7 | 479,3 | 478,3 | 4 |
| 7 | 939,5 | 470,3 | 922,5 | 921,5 | I | 359,3 | | 342,2 | | 3 |
| 8 | 1 038,6 | 519,8 | 1 021,5 | 1 020,6 | V | 246,2 | | 229,2 | | 2 |
| 9 | 1 184,7 | 592,8 | 1 167,7 | 1 166,7 | K | 147,1 | | 130,1 | | 1 |

# #13 K395

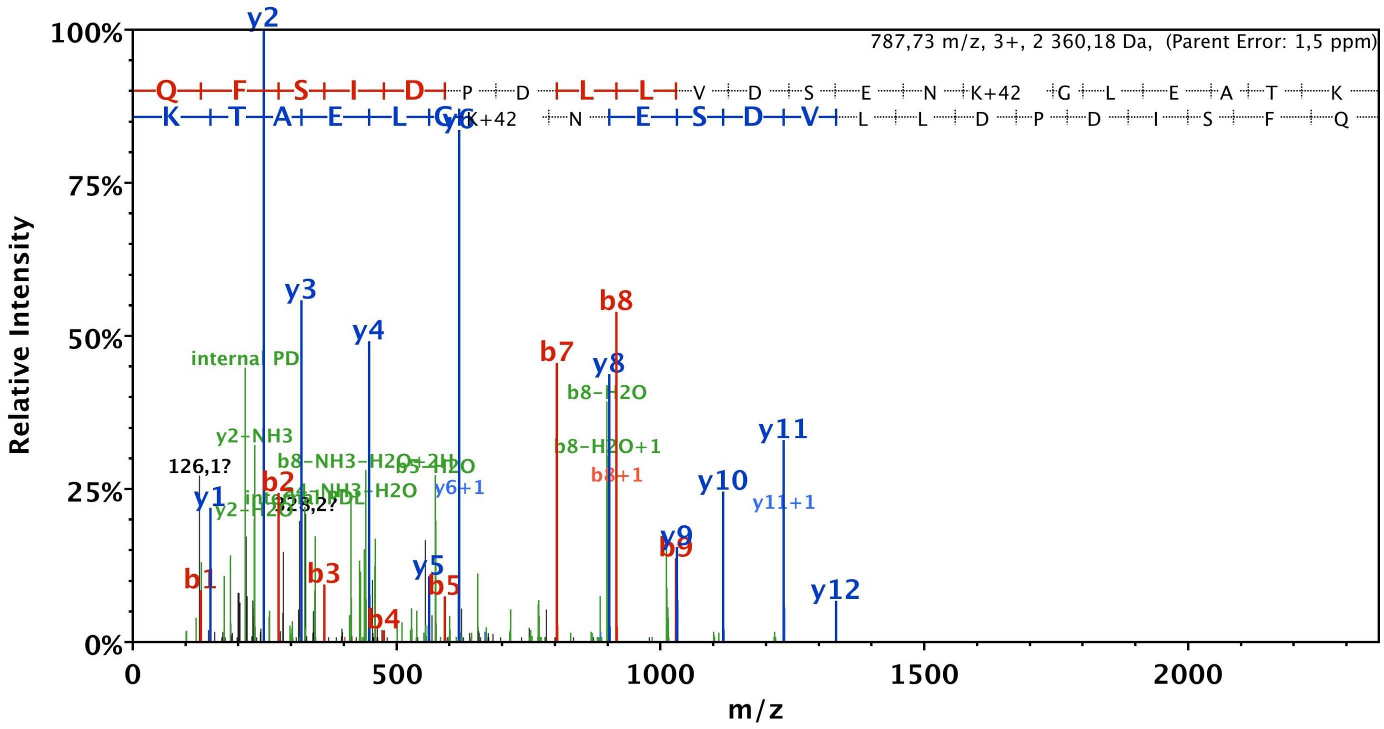


| B | B Ions | B+2H | B-NH3 | B-H2O | AA | Y Ions | Y+2H | Y-NH3 | Y-H2O | Y |
|---|---|---|---|---|---|---|---|---|---|---|
| 1 | 129,1 | 65,0 | 112,0 | | Q | 2 361,2 | 1 181,1 | 2 344,2 | 2 343,2 | 21 |
| 2 | 276,1 | 138,6 | 259,1 | | F | 2 233,1 | 1 117,1 | 2 216,1 | 2 215,1 | 20 |
| 3 | 363,2 | 182,1 | 346,1 | 345,2 | S | 2 086,1 | 1 043,5 | 2 069,0 | 2 068,0 | 19 |
| 4 | 476,3 | 238,6 | 459,2 | 458,2 | I | 1 999,0 | 1 000,0 | 1 982,0 | 1 981,0 | 18 |
| 5 | 591,3 | 296,1 | 574,3 | 573,3 | D | 1 885,9 | 943,5 | 1 868,9 | 1 867,9 | 17 |
| 6 | 688,3 | 344,7 | 671,3 | 670,3 | P | 1 770,9 | 886,0 | 1 753,9 | 1 752,9 | 16 |
| 7 | 803,4 | 402,2 | 786,3 | 785,3 | D | 1 673,9 | 837,4 | 1 656,8 | 1 655,8 | 15 |
| 8 | 916,4 | 458,7 | 899,4 | 898,4 | L | 1 558,8 | 779,9 | 1 541,8 | 1 540,8 | 14 |
| 9 | 1 029,5 | 515,3 | 1 012,5 | 1 011,5 | L | 1 445,7 | 723,4 | 1 428,7 | 1 427,7 | 13 |
| 10 | 1 128,6 | 564,8 | 1 111,6 | 1 110,6 | V | 1 332,7 | 666,8 | 1 315,6 | 1 314,7 | 12 |
| 11 | 1 243,6 | 622,3 | 1 226,6 | 1 225,6 | D | 1 233,6 | 617,3 | 1 216,6 | 1 215,6 | 11 |
| 12 | 1 330,7 | 665,8 | 1 313,6 | 1 312,6 | S | 1 118,6 | 559,8 | 1 101,5 | 1 100,6 | 10 |
| 13 | 1 459,7 | 730,4 | 1 442,7 | 1 441,7 | E | 1 031,5 | 516,3 | 1 014,5 | 1 013,5 | 9 |
| 14 | 1 573,7 | 787,4 | 1 556,7 | 1 555,7 | N | 902,5 | 451,8 | 885,5 | 884,5 | 8 |
| 15 | 1 743,8 | 872,4 | 1 726,8 | 1 725,8 | K+42 | 788,5 | 394,7 | 771,4 | 770,4 | 7 |
| 16 | 1 800,9 | 900,9 | 1 783,8 | 1 782,9 | G | 618,3 | 309,7 | 601,3 | 600,3 | 6 |
| 17 | 1 913,9 | 957,5 | 1 896,9 | 1 895,9 | L | 561,3 | 281,2 | 544,3 | 543,3 | 5 |
| 18 | 2 043,0 | 1 022,0 | 2 026,0 | 2 025,0 | E | 448,2 | 224,6 | 431,2 | 430,2 | 4 |
| 19 | 2 114,0 | 1 057,5 | 2 097,0 | 2 096,0 | A | 319,2 | 160,1 | 302,2 | 301,2 | 3 |
| 20 | 2 215,1 | 1 108,0 | 2 198,0 | 2 197,1 | T | 248,2 | 124,6 | 231,1 | 230,1 | 2 |
| 21 | 2 361,2 | 1 181,1 | 2 344,2 | 2 343,2 | K | 147,1 | 74,1 | 130,1 | | 1 |

# #14 K395

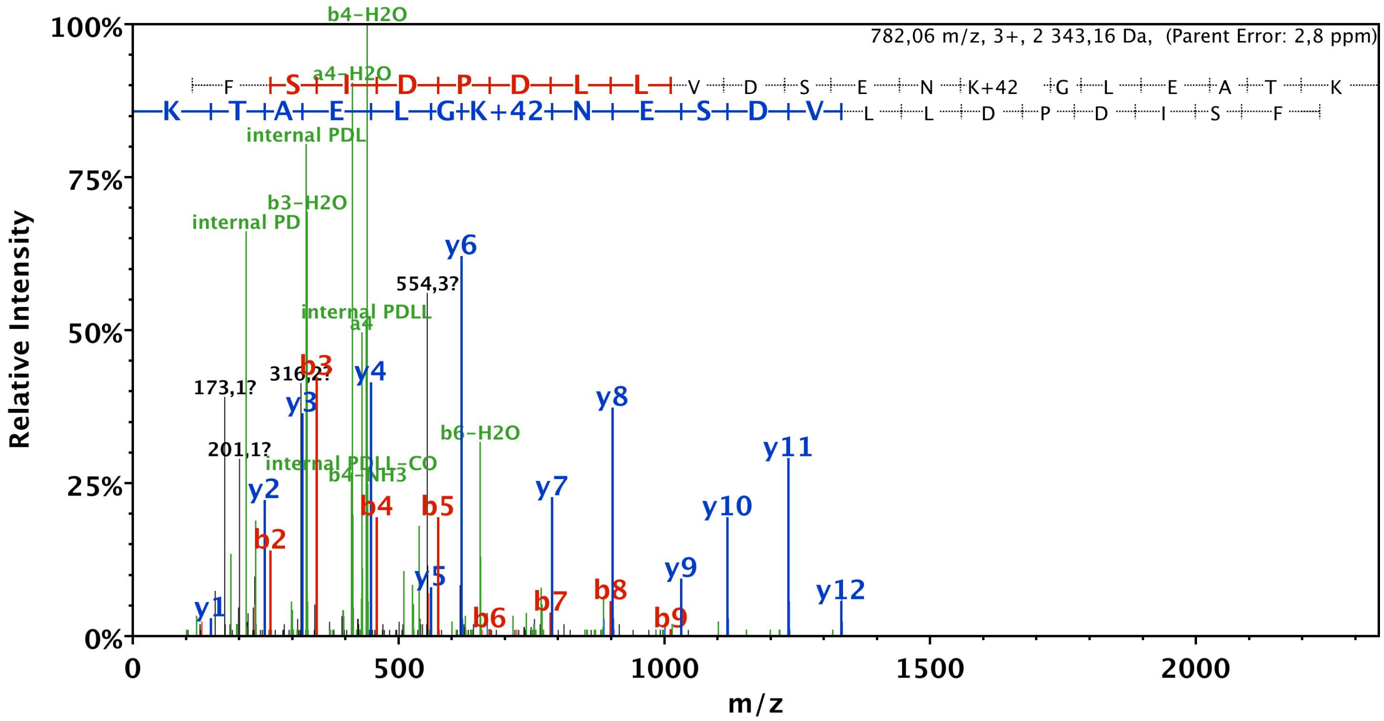


| B | B Ions | B+2H | B-NH3 | B-H2O | AA | Y Ions | Y+2H | Y-NH3 | Y-H2O | Y |
|---|---|---|---|---|---|---|---|---|---|---|
| 1 | 112,0 | 56,5 | 95,0 | | Q-17 | 2 344,2 | 1 172,6 | 2 327,1 | 2 326,1 | 21 |
| 2 | 259,1 | 130,1 | 242,1 | | F | 2 233,1 | 1 117,1 | 2 216,1 | 2 215,1 | 20 |
| 3 | 346,1 | 173,6 | 329,1 | 328,1 | S | 2 086,1 | 1 043,5 | 2 069,0 | 2 068,0 | 19 |
| 4 | 459,2 | 230,1 | 442,2 | 441,2 | I | 1 999,0 | 1 000,0 | 1 982,0 | 1 981,0 | 18 |
| 5 | 574,3 | 287,6 | 557,2 | 556,2 | D | 1 885,9 | 943,5 | 1 868,9 | 1 867,9 | 17 |
| 6 | 671,3 | 336,2 | 654,3 | 653,3 | P | 1 770,9 | 886,0 | 1 753,9 | 1 752,9 | 16 |
| 7 | 786,3 | 393,7 | 769,3 | 768,3 | D | 1 673,9 | 837,4 | 1 656,8 | 1 655,8 | 15 |
| 8 | 899,4 | 450,2 | 882,4 | 881,4 | L | 1 558,8 | 779,9 | 1 541,8 | 1 540,8 | 14 |
| 9 | 1 012,5 | 506,8 | 995,5 | 994,5 | L | 1 445,7 | 723,4 | 1 428,7 | 1 427,7 | 13 |
| 10 | 1 111,6 | 556,3 | 1 094,5 | 1 093,6 | V | 1 332,7 | 666,8 | 1 315,6 | 1 314,7 | 12 |
| 11 | 1 226,6 | 613,8 | 1 209,6 | 1 208,6 | D | 1 233,6 | 617,3 | 1 216,6 | 1 215,6 | 11 |
| 12 | 1 313,6 | 657,3 | 1 296,6 | 1 295,6 | S | 1 118,6 | 559,8 | 1 101,5 | 1 100,6 | 10 |
| 13 | 1 442,7 | 721,8 | 1 425,6 | 1 424,7 | E | 1 031,5 | 516,3 | 1 014,5 | 1 013,5 | 9 |
| 14 | 1 556,7 | 778,9 | 1 539,7 | 1 538,7 | N | 902,5 | 451,8 | 885,5 | 884,5 | 8 |
| 15 | 1 726,8 | 863,9 | 1 709,8 | 1 708,8 | K+42 | 788,5 | 394,7 | 771,4 | 770,4 | 7 |
| 16 | 1 783,8 | 892,4 | 1 766,8 | 1 765,8 | G | 618,3 | 309,7 | 601,3 | 600,3 | 6 |
| 17 | 1 896,9 | 949,0 | 1 879,9 | 1 878,9 | L | 561,3 | 281,2 | 544,3 | 543,3 | 5 |
| 18 | 2 026,0 | 1 013,5 | 2 008,9 | 2 008,0 | E | 448,2 | 224,6 | 431,2 | 430,2 | 4 |
| 19 | 2 097,0 | 1 049,0 | 2 080,0 | 2 079,0 | A | 319,2 | 160,1 | 302,2 | 301,2 | 3 |
| 20 | 2 198,1 | 1 099,5 | 2 181,0 | 2 180,0 | T | 248,2 | 124,6 | 231,1 | 230,1 | 2 |
| 21 | 2 344,2 | 1 172,6 | 2 327,1 | 2 326,1 | K | 147,1 | 74,1 | 130,1 | | 1 |

# #15 K401

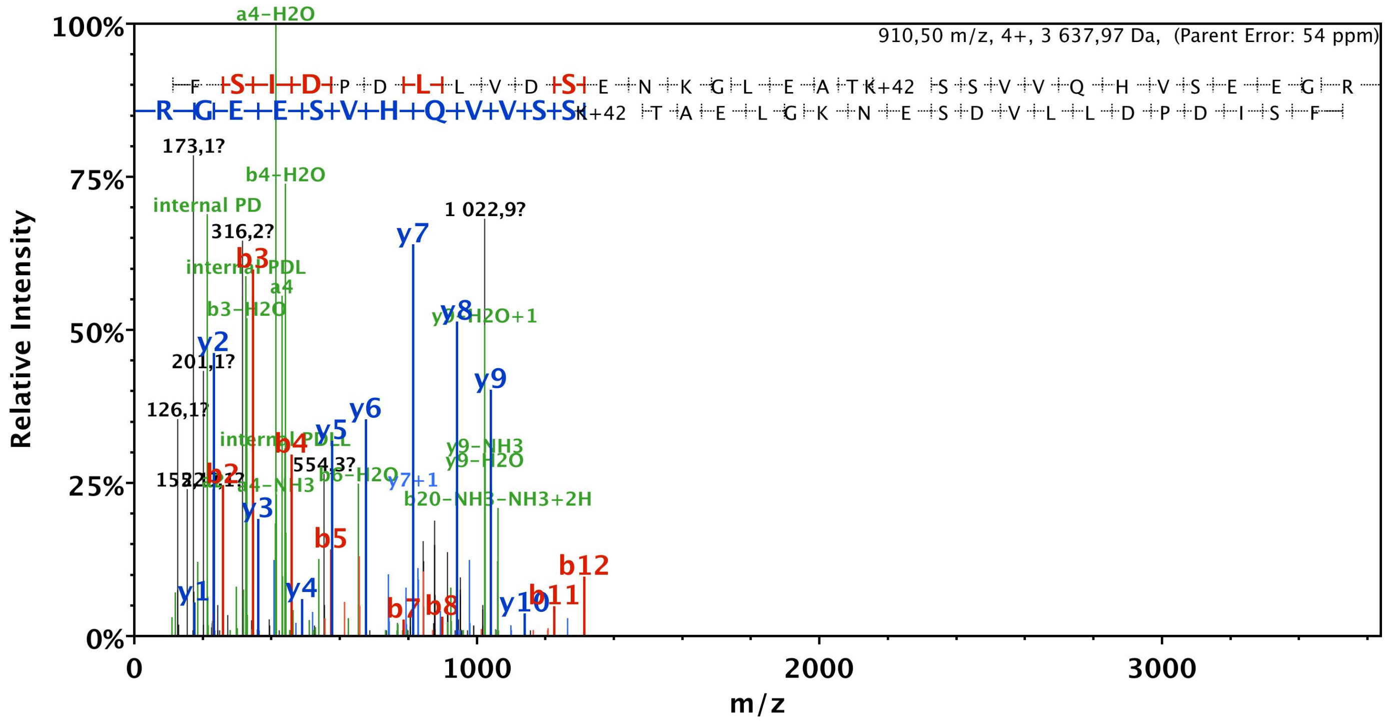


| B | B Ions | B+2H | B-NH3 | B-H2O | AA | Y Ions | Y+2H | Y-NH3 | Y-H2O | Y |
|---|---|---|---|---|---|---|---|---|---|---|
| 1 | 112,0 | 56,5 | 95,0 | | Q-17 | 3 638,8 | 1 819,9 | 3 621,8 | 3 620,8 | 33 |
| 2 | 259,1 | 130,1 | 242,1 | | F | 3 527,8 | 1 764,4 | 3 510,7 | 3 509,7 | 32 |
| 3 | 346,1 | 173,6 | 329,1 | 328,1 | S | 3 380,7 | 1 690,8 | 3 363,7 | 3 362,7 | 31 |
| 4 | 459,2 | 230,1 | 442,2 | 441,2 | I | 3 293,6 | 1 647,3 | 3 276,6 | 3 275,6 | 30 |
| 5 | 574,3 | 287,6 | 557,2 | 556,2 | D | 3 180,6 | 1 590,8 | 3 163,5 | 3 162,6 | 29 |
| 6 | 671,3 | 336,2 | 654,3 | 653,3 | P | 3 065,5 | 1 533,3 | 3 048,5 | 3 047,5 | 28 |
| 7 | 786,3 | 393,7 | 769,3 | 768,3 | D | 2 968,5 | 1 484,7 | 2 951,5 | 2 950,5 | 27 |
| 8 | 899,4 | 450,2 | 882,4 | 881,4 | L | 2 853,5 | 1 427,2 | 2 836,4 | 2 835,4 | 26 |
| 9 | 1 012,5 | 506,8 | 995,5 | 994,5 | L | 2 740,4 | 1 370,7 | 2 723,3 | 2 722,4 | 25 |
| 10 | 1 111,6 | 556,3 | 1 094,5 | 1 093,6 | V | 2 627,3 | 1 314,1 | 2 610,3 | 2 609,3 | 24 |
| 11 | 1 226,6 | 613,8 | 1 209,6 | 1 208,6 | D | 2 528,2 | 1 264,6 | 2 511,2 | 2 510,2 | 23 |
| 12 | 1 313,6 | 657,3 | 1 296,6 | 1 295,6 | S | 2 413,2 | 1 207,1 | 2 396,2 | 2 395,2 | 22 |
| 13 | 1 442,7 | 721,8 | 1 425,6 | 1 424,7 | E | 2 326,2 | 1 163,6 | 2 309,1 | 2 308,2 | 21 |
| 14 | 1 556,7 | 778,9 | 1 539,7 | 1 538,7 | N | 2 197,1 | 1 099,1 | 2 180,1 | 2 179,1 | 20 |
| 15 | 1 684,8 | 842,9 | 1 667,8 | 1 666,8 | K | 2 083,1 | 1 042,0 | 2 066,1 | 2 065,1 | 19 |
| 16 | 1 741,8 | 871,4 | 1 724,8 | 1 723,8 | G | 1 955,0 | 978,0 | 1 938,0 | 1 937,0 | 18 |
| 17 | 1 854,9 | 928,0 | 1 837,9 | 1 836,9 | L | 1 898,0 | 949,5 | 1 880,9 | 1 880,0 | 17 |
| 18 | 1 984,0 | 992,5 | 1 966,9 | 1 965,9 | E | 1 784,9 | 892,9 | 1 767,9 | 1 766,9 | 16 |
| 19 | 2 055,0 | 1 028,0 | 2 038,0 | 2 037,0 | A | 1 655,8 | 828,4 | 1 638,8 | 1 637,8 | 15 |
| 20 | 2 156,0 | 1 078,5 | 2 139,0 | 2 138,0 | T | 1 584,8 | 792,9 | 1 567,8 | 1 566,8 | 14 |
| 21 | 2 326,1 | 1 163,6 | 2 309,1 | 2 308,1 | K+42 | 1 483,7 | 742,4 | 1 466,7 | 1 465,7 | 13 |
| 22 | 2 413,2 | 1 207,1 | 2 396,2 | 2 395,2 | S | 1 313,6 | 657,3 | 1 296,6 | 1 295,6 | 12 |
| 23 | 2 500,2 | 1 250,6 | 2 483,2 | 2 482,2 | S | 1 226,6 | 613,8 | 1 209,6 | 1 208,6 | 11 |
| 24 | 2 599,3 | 1 300,1 | 2 582,3 | 2 581,3 | V | 1 139,6 | 570,3 | 1 122,6 | 1 121,6 | 10 |
| 25 | 2 698,3 | 1 349,7 | 2 681,3 | 2 680,3 | V | 1 040,5 | 520,8 | 1 023,5 | 1 022,5 | 9 |
| 26 | 2 826,4 | 1 413,7 | 2 809,4 | 2 808,4 | Q | 941,4 | 471,2 | 924,4 | 923,4 | 8 |
| 27 | 2 963,5 | 1 482,2 | 2 946,4 | 2 945,5 | H | 813,4 | 407,2 | 796,4 | 795,4 | 7 |
| 28 | 3 062,5 | 1 531,8 | 3 045,5 | 3 044,5 | V | 676,3 | 338,7 | 659,3 | 658,3 | 6 |
| 29 | 3 149,6 | 1 575,3 | 3 132,5 | 3 131,6 | S | 577,3 | 289,1 | 560,2 | 559,2 | 5 |
| 30 | 3 278,6 | 1 639,8 | 3 261,6 | 3 260,6 | E | 490,2 | 245,6 | 473,2 | 472,2 | 4 |
| 31 | 3 407,6 | 1 704,3 | 3 390,6 | 3 389,6 | E | 361,2 | 181,1 | 344,2 | 343,2 | 3 |
| 32 | 3 464,7 | 1 732,8 | 3 447,6 | 3 446,7 | G | 232,1 | 116,6 | 215,1 | | 2 |
| 33 | 3 638,8 | 1 819,9 | 3 621,8 | 3 620,8 | R | 175,1 | 88,1 | 158,1 | | 1 |

# #16 K401

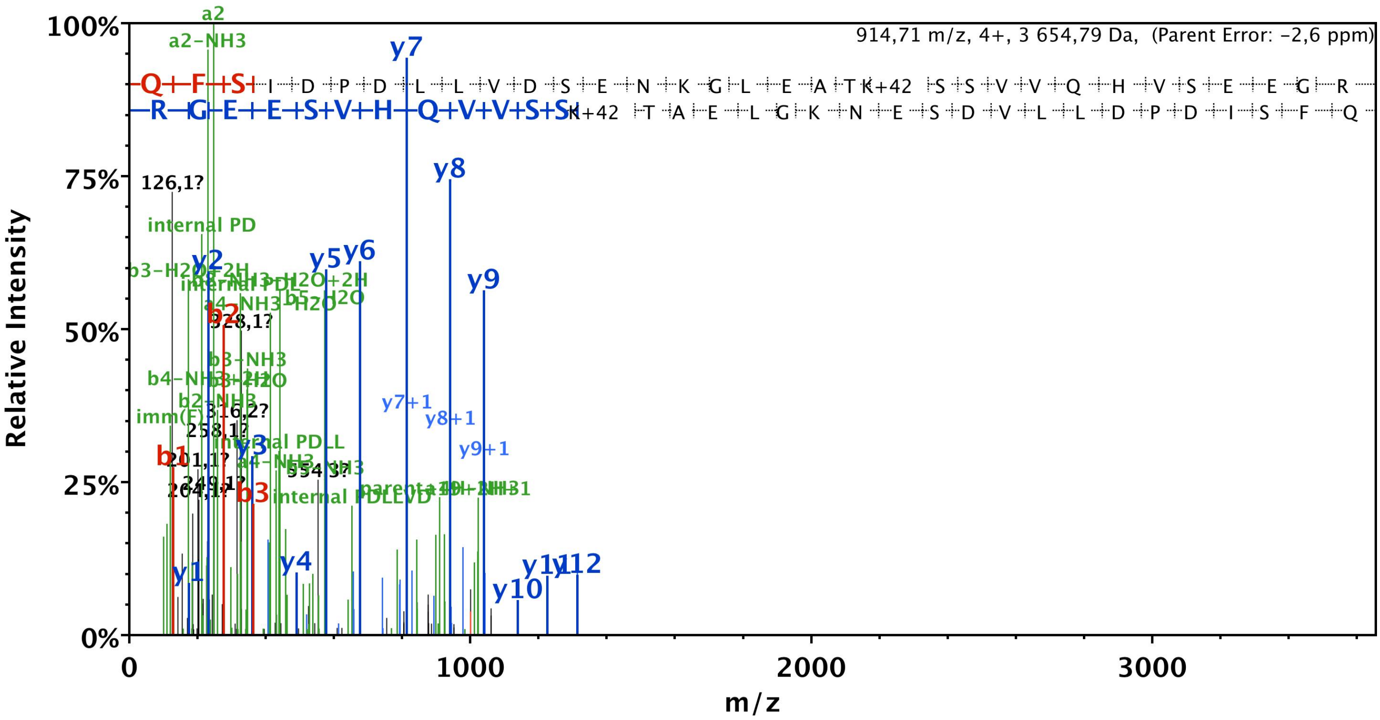


| B | B Ions | B+2H | B-NH3 | B-H2O | AA | Y Ions | Y+2H | Y-NH3 | Y-H2O | Y |
|---|---|---|---|---|---|---|---|---|---|---|
| 1 | 129,1 | 65,0 | 112,0 | | Q | 3 655,8 | 1 828,4 | 3 638,8 | 3 637,8 | 33 |
| 2 | 276,1 | 138,6 | 259,1 | | F | 3 527,8 | 1 764,4 | 3 510,7 | 3 509,7 | 32 |
| 3 | 363,2 | 182,1 | 346,1 | 345,2 | S | 3 380,7 | 1 690,8 | 3 363,7 | 3 362,7 | 31 |
| 4 | 476,3 | 238,6 | 459,2 | 458,2 | I | 3 293,6 | 1 647,3 | 3 276,6 | 3 275,6 | 30 |
| 5 | 591,3 | 296,1 | 574,3 | 573,3 | D | 3 180,6 | 1 590,8 | 3 163,5 | 3 162,6 | 29 |
| 6 | 688,3 | 344,7 | 671,3 | 670,3 | P | 3 065,5 | 1 533,3 | 3 048,5 | 3 047,5 | 28 |
| 7 | 803,4 | 402,2 | 786,3 | 785,3 | D | 2 968,5 | 1 484,7 | 2 951,5 | 2 950,5 | 27 |
| 8 | 916,4 | 458,7 | 899,4 | 898,4 | L | 2 853,5 | 1 427,2 | 2 836,4 | 2 835,4 | 26 |
| 9 | 1 029,5 | 515,3 | 1 012,5 | 1 011,5 | L | 2 740,4 | 1 370,7 | 2 723,3 | 2 722,4 | 25 |
| 10 | 1 128,6 | 564,8 | 1 111,6 | 1 110,6 | V | 2 627,3 | 1 314,1 | 2 610,3 | 2 609,3 | 24 |
| 11 | 1 243,6 | 622,3 | 1 226,6 | 1 225,6 | D | 2 528,2 | 1 264,6 | 2 511,2 | 2 510,2 | 23 |
| 12 | 1 330,7 | 665,8 | 1 313,6 | 1 312,6 | S | 2 413,2 | 1 207,1 | 2 396,2 | 2 395,2 | 22 |
| 13 | 1 459,7 | 730,4 | 1 442,7 | 1 441,7 | E | 2 326,2 | 1 163,6 | 2 309,1 | 2 308,2 | 21 |
| 14 | 1 573,7 | 787,4 | 1 556,7 | 1 555,7 | N | 2 197,1 | 1 099,1 | 2 180,1 | 2 179,1 | 20 |
| 15 | 1 701,8 | 851,4 | 1 684,8 | 1 683,8 | K | 2 083,1 | 1 042,0 | 2 066,1 | 2 065,1 | 19 |
| 16 | 1 758,9 | 879,9 | 1 741,8 | 1 740,8 | G | 1 955,0 | 978,0 | 1 938,0 | 1 937,0 | 18 |
| 17 | 1 871,9 | 936,5 | 1 854,9 | 1 853,9 | L | 1 898,0 | 949,5 | 1 880,9 | 1 880,0 | 17 |
| 18 | 2 001,0 | 1 001,0 | 1 984,0 | 1 983,0 | E | 1 784,9 | 892,9 | 1 767,9 | 1 766,9 | 16 |
| 19 | 2 072,0 | 1 036,5 | 2 055,0 | 2 054,0 | A | 1 655,8 | 828,4 | 1 638,8 | 1 637,8 | 15 |
| 20 | 2 173,1 | 1 087,0 | 2 156,0 | 2 155,1 | T | 1 584,8 | 792,9 | 1 567,8 | 1 566,8 | 14 |
| 21 | 2 343,2 | 1 172,1 | 2 326,1 | 2 325,2 | K+42 | 1 483,7 | 742,4 | 1 466,7 | 1 465,7 | 13 |
| 22 | 2 430,2 | 1 215,6 | 2 413,2 | 2 412,2 | S | 1 313,6 | 657,3 | 1 296,6 | 1 295,6 | 12 |
| 23 | 2 517,2 | 1 259,1 | 2 500,2 | 2 499,2 | S | 1 226,6 | 613,8 | 1 209,6 | 1 208,6 | 11 |
| 24 | 2 616,3 | 1 308,7 | 2 599,3 | 2 598,3 | V | 1 139,6 | 570,3 | 1 122,6 | 1 121,6 | 10 |
| 25 | 2 715,4 | 1 358,2 | 2 698,3 | 2 697,4 | V | 1 040,5 | 520,8 | 1 023,5 | 1 022,5 | 9 |
| 26 | 2 843,4 | 1 422,2 | 2 826,4 | 2 825,4 | Q | 941,4 | 471,2 | 924,4 | 923,4 | 8 |
| 27 | 2 980,5 | 1 490,7 | 2 963,5 | 2 962,5 | H | 813,4 | 407,2 | 796,4 | 795,4 | 7 |
| 28 | 3 079,6 | 1 540,3 | 3 062,5 | 3 061,5 | V | 676,3 | 338,7 | 659,3 | 658,3 | 6 |
| 29 | 3 166,6 | 1 583,8 | 3 149,6 | 3 148,6 | S | 577,3 | 289,1 | 560,2 | 559,2 | 5 |
| 30 | 3 295,6 | 1 648,3 | 3 278,6 | 3 277,6 | E | 490,2 | 245,6 | 473,2 | 472,2 | 4 |
| 31 | 3 424,7 | 1 712,8 | 3 407,6 | 3 406,7 | E | 361,2 | 181,1 | 344,2 | 343,2 | 3 |
| 32 | 3 481,7 | 1 741,4 | 3 464,7 | 3 463,7 | G | 232,1 | 116,6 | 215,1 | | 2 |
| 33 | 3 655,8 | 1 828,4 | 3 638,8 | 3 637,8 | R | 175,1 | 88,1 | 158,1 | | 1 |

# #17 K401

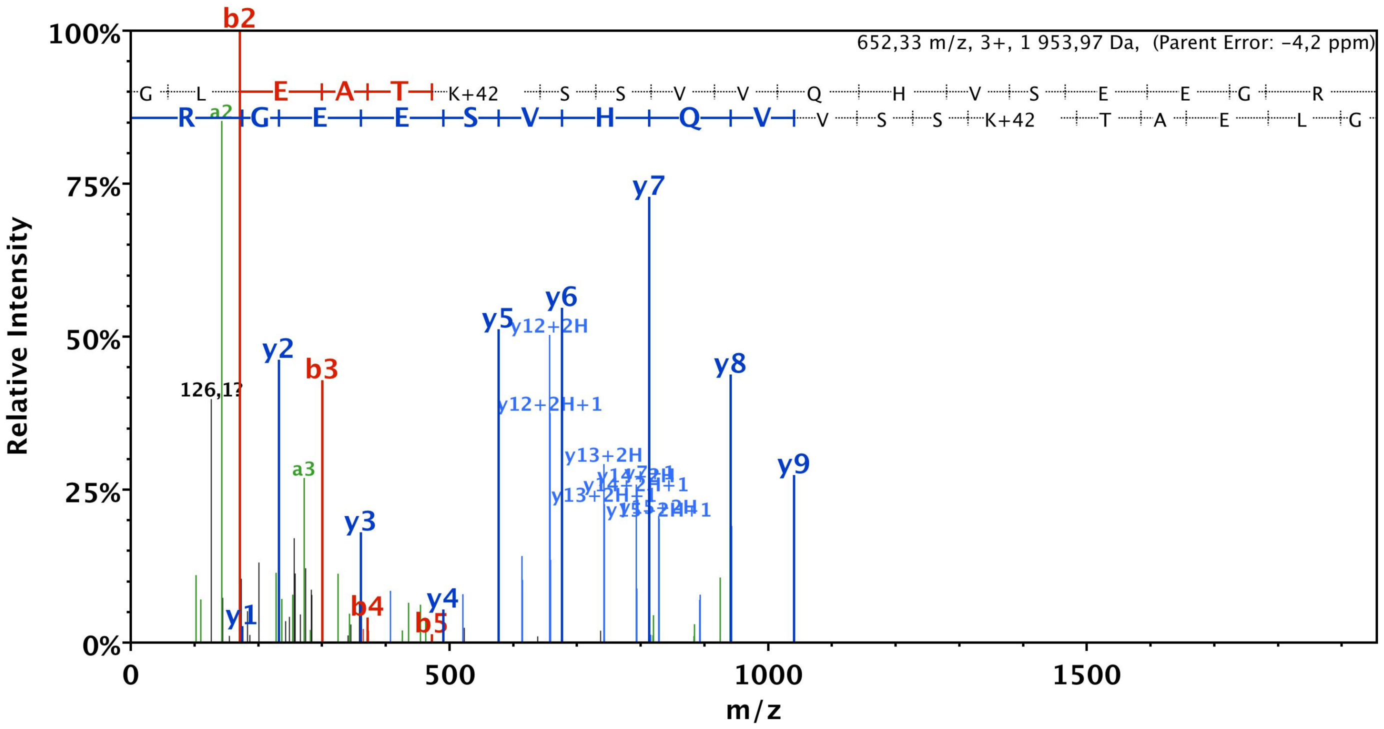


| B | B Ions | B+2H | B-NH3 | B-H2O | AA | Y Ions | Y+2H | Y-NH3 | Y-H2O | Y |
|---|---|---|---|---|---|---|---|---|---|---|
| 1 | 58,0 | 29,5 | | | G | 1 955,0 | 978,0 | 1 938,0 | 1 937,0 | 18 |
| 2 | 171,1 | 86,1 | | | L | 1 898,0 | 949,5 | 1 880,9 | 1 880,0 | 17 |
| 3 | 300,2 | 150,6 | | 282,1 | E | 1 784,9 | 892,9 | 1 767,9 | 1 766,9 | 16 |
| 4 | 371,2 | 186,1 | | 353,2 | A | 1 655,8 | 828,4 | 1 638,8 | 1 637,8 | 15 |
| 5 | 472,2 | 236,6 | | 454,2 | T | 1 584,8 | 792,9 | 1 567,8 | 1 566,8 | 14 |
| 6 | 642,3 | 321,7 | 625,3 | 624,3 | K+42 | 1 483,7 | 742,4 | 1 466,7 | 1 465,7 | 13 |
| 7 | 729,4 | 365,2 | 712,4 | 711,4 | S | 1 313,6 | 657,3 | 1 296,6 | 1 295,6 | 12 |
| 8 | 816,4 | 408,7 | 799,4 | 798,4 | S | 1 226,6 | 613,8 | 1 209,6 | 1 208,6 | 11 |
| 9 | 915,5 | 458,2 | 898,5 | 897,5 | V | 1 139,6 | 570,3 | 1 122,6 | 1 121,6 | 10 |
| 10 | 1 014,5 | 507,8 | 997,5 | 996,5 | V | 1 040,5 | 520,8 | 1 023,5 | 1 022,5 | 9 |
| 11 | 1 142,6 | 571,8 | 1 125,6 | 1 124,6 | Q | 941,4 | 471,2 | 924,4 | 923,4 | 8 |
| 12 | 1 279,7 | 640,3 | 1 262,6 | 1 261,7 | H | 813,4 | 407,2 | 796,4 | 795,4 | 7 |
| 13 | 1 378,7 | 689,9 | 1 361,7 | 1 360,7 | V | 676,3 | 338,7 | 659,3 | 658,3 | 6 |
| 14 | 1 465,8 | 733,4 | 1 448,7 | 1 447,8 | S | 577,3 | 289,1 | 560,2 | 559,2 | 5 |
| 15 | 1 594,8 | 797,9 | 1 577,8 | 1 576,8 | E | 490,2 | 245,6 | 473,2 | 472,2 | 4 |
| 16 | 1 723,8 | 862,4 | 1 706,8 | 1 705,8 | E | 361,2 | 181,1 | 344,2 | 343,2 | 3 |
| 17 | 1 780,9 | 890,9 | 1 763,8 | 1 762,9 | G | 232,1 | 116,6 | 215,1 | | 2 |
| 18 | 1 955,0 | 978,0 | 1 938,0 | 1 937,0 | R | 175,1 | 88,1 | 158,1 | | 1 |

# #18 K401

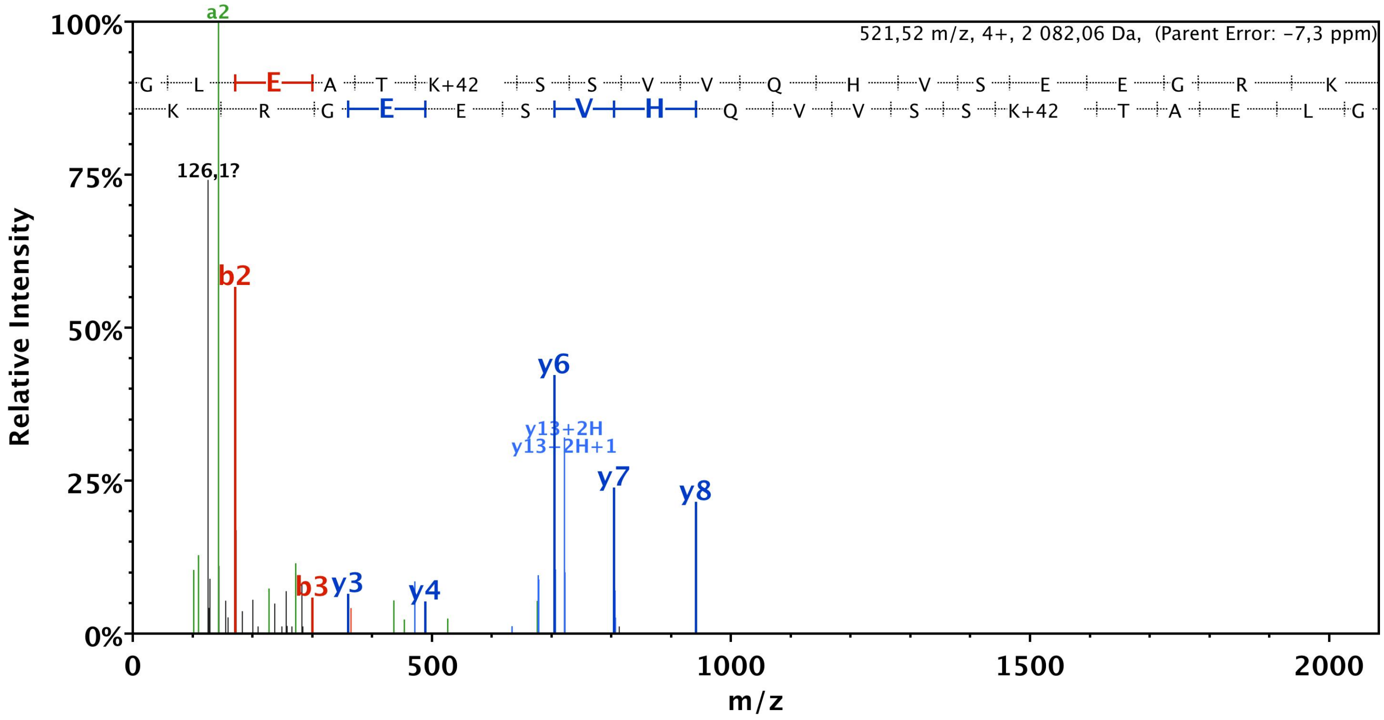


| B | B Ions | B+2H | B-NH3 | B-H2O | AA | Y Ions | Y+2H | Y-NH3 | Y-H2O | Y |
|---|---|---|---|---|---|---|---|---|---|---|
| 1 | 58,0 | 29,5 | | | G | 2 083,1 | 1 042,0 | 2 066,1 | 2 065,1 | 19 |
| 2 | 171,1 | 86,1 | | | L | 2 026,1 | 1 013,5 | 2 009,0 | 2 008,0 | 18 |
| 3 | 300,2 | 150,6 | | 282,1 | E | 1 913,0 | 957,0 | 1 895,9 | 1 895,0 | 17 |
| 4 | 371,2 | 186,1 | | 353,2 | A | 1 783,9 | 892,5 | 1 766,9 | 1 765,9 | 16 |
| 5 | 472,2 | 236,6 | | 454,2 | T | 1 712,9 | 856,9 | 1 695,9 | 1 694,9 | 15 |
| 6 | 642,3 | 321,7 | 625,3 | 624,3 | K+42 | 1 611,8 | 806,4 | 1 594,8 | 1 593,8 | 14 |
| 7 | 729,4 | 365,2 | 712,4 | 711,4 | S | 1 441,7 | 721,4 | 1 424,7 | 1 423,7 | 13 |
| 8 | 816,4 | 408,7 | 799,4 | 798,4 | S | 1 354,7 | 677,9 | 1 337,7 | 1 336,7 | 12 |
| 9 | 915,5 | 458,2 | 898,5 | 897,5 | V | 1 267,7 | 634,3 | 1 250,6 | 1 249,7 | 11 |
| 10 | 1 014,5 | 507,8 | 997,5 | 996,5 | V | 1 168,6 | 584,8 | 1 151,6 | 1 150,6 | 10 |
| 11 | 1 142,6 | 571,8 | 1 125,6 | 1 124,6 | Q | 1 069,5 | 535,3 | 1 052,5 | 1 051,5 | 9 |
| 12 | 1 279,7 | 640,3 | 1 262,6 | 1 261,7 | H | 941,5 | 471,2 | 924,5 | 923,5 | 8 |
| 13 | 1 378,7 | 689,9 | 1 361,7 | 1 360,7 | V | 804,4 | 402,7 | 787,4 | 786,4 | 7 |
| 14 | 1 465,8 | 733,4 | 1 448,7 | 1 447,8 | S | 705,4 | 353,2 | 688,3 | 687,3 | 6 |
| 15 | 1 594,8 | 797,9 | 1 577,8 | 1 576,8 | E | 618,3 | 309,7 | 601,3 | 600,3 | 5 |
| 16 | 1 723,8 | 862,4 | 1 706,8 | 1 705,8 | E | 489,3 | 245,1 | 472,3 | 471,3 | 4 |
| 17 | 1 780,9 | 890,9 | 1 763,8 | 1 762,9 | G | 360,2 | 180,6 | 343,2 | | 3 |
| 18 | 1 937,0 | 969,0 | 1 919,9 | 1 919,0 | R | 303,2 | 152,1 | 286,2 | | 2 |
| 19 | 2 083,1 | 1 042,0 | 2 066,1 | 2 065,1 | K | 147,1 | 74,1 | 130,1 | | 1 |
