## Supplemental Table S3 for "CBP/EP300 acetylates and stabilizes the stress-responsive Heat Shock Factor 2, a process compromised in Rubinstein-Taybi syndrome"

**Table S3. Docking study between the HR-A/B domain of HSF2 and the KIX domain of CBP/EP300**

Docking study - between HR-A/B domain of HSF2 and KIX domain of CBP/EP300 - was processed using *ZDOCK (score) and* FireDock *(energetic score)* and the best *pose* of the complex were sorted in the absence of mutation (belong to the Top 3 for Zdock and Top 1 for Firedock) and in the presence of mutations, within the KIX domain (Y650), or in the HR-A/B domain (V183A, F181A, Q180A, K177A). In red, mutations that impair the interaction between the HR-A/B domain of HSF2 and the KIX domain of CBP/EP300.

| **Complex** | **Zdock** | **Firedock** |
|---|---|---|
| **Complex HR-A/B_KIX** | **14.66** | **-13.04** |
| **Mutation Y650A** | 14.38 | **0.70** |
| Mutation V183A | 14.10 | -5.42 |
| Mutation F181A | 14.22 | -6.16 |
| Mutation Q180A | 14.38 | -2.36 |
| Mutation K177A | 14.16 | -8.33 |
